# Somatic Mutations in Collagens are Associated with a Distinct Tumor Environment and Overall Survival in Gastric Cancer

**DOI:** 10.1101/2021.01.19.426938

**Authors:** Alexander S. Brodsky, Jay Khurana, Kevin S. Guo, Elizabeth Y. Wu, Dongfang Yang, Ayesha S. Siddique, Ian Y. Wong, Ece D. Gamsiz Uzun, Murray B. Resnick

## Abstract

**Background:** Gastric cancer is a heterogeneous disease with poorly understood genetic and microenvironmental factors. Mutations in collagen genes are associated with genetic diseases that compromise tissue integrity, but their role in tumor progression has not been extensively reported. Aberrant collagen expression has been long associated with malignant tumor growth, invasion, chemoresistance, and patient outcomes. We hypothesized that somatic mutations in collagens could functionally alter the tumor extracellular matrix.

**Methods:** We used publicly available datasets including The Tumor Cancer Genome Atlas (TCGA) to interrogate somatic mutations in collagens in stomach adenocarcinomas. To demonstrate that collagens were significantly mutated above background mutation rates, we used a moderated Kolmogorov-Smirnov test along with combination analysis with a bootstrap approach to define the background accounting for mutation rates. Association between mutations and clinicopathological features was evaluated by Fisher or chi-squared tests. Association with overall survival was assessed by Kaplan-Meier and the Cox-Proportional Hazards Model. Gene Set Enrichment Analysis was used to interrogate pathways. Immunohistochemistry and *in situ* hybridization tested expression of COL7A1 in stomach tumors.

**Results:** In stomach adenocarcinomas, we identified individual collagen genes and sets of collagen genes harboring somatic mutations at a high frequency compared to background in both microsatellite stable, and microsatellite instable tumors in TCGA. Many of the missense mutations resemble the same types of loss of function mutations in collagenopathies that disrupt tissue formation and destabilize cells providing guidance to interpret the somatic mutations. We identified combinations of somatic mutations in collagens associated with overall survival, with a distinctive tumor microenvironment marked by lower matrisome expression and immune cell signatures. Truncation mutations were strongly associated with improved outcomes suggesting that loss of expression of secreted collagens impact tumor progression and treatment response. Germline collagenopathy variants guided interpretation of impactful somatic mutations on tumors.

**Conclusions:** These observations highlight that many collagens, expressed in non-physiologically relevant conditions in tumors, harbor impactful somatic mutations in tumors, suggesting new approaches for classification and therapy development in stomach cancer. In sum, these findings demonstrate how classification of tumors by collagen mutations identified strong links between specific genotypes and the tumor environment.

**Research Highlights:** Collagen mutations are prevalent in stomach cancer

Collagen somatic missense mutations resemble collagenopathy mutations

Collagen mutations associate with overall survival in stomach cancer

Tumors with collagen mutations have distinct molecular pathways and tumor microenvironments

## Background

Collagens are the most abundant proteins in extracellular matrix and are critical components and regulators of the tumor microenvironment (1, 2). Increased collagen expression in many solid tumors has been associated with poor outcomes and resistance in multiple settings (3), likely through increased epithelial-to-mesenchymal transitions (EMT) and drug resistance (4). The 28 members of the collagen family are expressed by 43 genes and defined by the common triple helix motif. Collagens are classified into families including fibrillar collagens (i.e. Collagen type I, II, III, V, XI, XIV), network collagens (i.e. Collagen type IV), membrane (i.e. type XVII) and other (type VII, XXVIII) (5) (Table S1). Although most studies have focused on the most abundant collagen, collagen type I, in cancer there is increasing awareness of the role of many minor collagens in cancer such as types X and XI (6, 7). Minor collagens are defined by their low abundance compared to the major fibrillar collagen types such as type I, but nonetheless they have critical functions and large impacts on tissues. Collagen structures are very complex because of the tendency to form heterotrimers, interact with each other, post-translational modifications and regulation through crosslinking (5). The breadth of mechanisms by which collagens mediate tumor progression is not yet understood and collagens could have context dependent functions in tumors analogous to their normal tissue specific expression and functions (4). The cellular origin of collagens is not always clear as both cancer and stroma cells are known to secrete collagens as indicated both by *in situ* hybridization studies (8, 9) and recent proteomic studies also suggest tumor cells secrete collagens (10).

**Table 1.** Association of collagen mutation status with clinicopathological characteristics.

| Factor | MSI Status (MSI-H, MSI-L, MSS) | All Collagen Mutations | All Collagens Missense | All Collagens Truncation | All COL7A1 Mutations | Mutation Rate (High/Low) |
| --- | --- | --- | --- | --- | --- | --- |
| **Microsatellite** |  | **P<1e-3** | **P<1e-3** | **P<1e-3** | **P<1e-3** | **P<1e-3** |
| MSI-H |  | 100% (85/85) | 100% (85/85) | 81% (69/85) | 41% (35/85) | 100% (84/84) |
| MSI-L |  | 63% (40/63) | 59% (37/63) | 13% (8/63) | 0% (0/63) | 59% (37/63) |
| MSS |  | 48% (143/295) | 46% (135/295) | 8% (23/295) | 2% (6/295) | 33% (98/293) |
| **Grade** | **P=0.5** | **P=0.9** | **P=0.9** | **P=0.9** | **P=0.7** | **P=0.3** |
| G1 | 33% (4/12) MSIH<br>8% (1/12) MSIL<br>58% (7/12) MSS | 67% (8/12) | 67% (8/12) | 33% (4/12) | 8% (1/12) | 67% (8/12) |
| G2 | 17% (26/156) MSIH<br>15% (23/156) MSIL<br>69% (107/156) MSS | 63% (98/156) | 61% (95/156) | 22% (35/156) | 7% (11/156) | 54% (82/153) |
| G3 | 21% (54/259) MSIH<br>15% (39/259) MSIL<br>64% (166/259) MSS | 61% (157/259) | 58% (149/259) | 23% (60/259) | 11% (28/259) | 47% (121/259) |
| GX | 11% (1/9) MSIH<br>0% (0/9) MSIL<br>89% (8/9) MSS | 56% (5/9) | 56% (5/9) | 11% (1/9) | 11% (1/9) | 67% (6/9) |
| **TNM Stage** | **P=0.5** | **P=0.4** | **P=0.4** | **P=0.3** | **P=0.4** | **P=0.10** |
| I | 26% (15/57) MSIH<br>18% (10/57) MSIL<br>56% (32/57) MSIH | 70% (40/57) | 68% (39/57) | 28% (16/57) | 7% (4/57) | 57% (32/56) |
| II | 21% (27/131) MSIH<br>15% (20/131) MSIL<br>64% (84/131) MSS | 61% (80/131) | 56% (74/131) | 24% (31/131) | 9% (12/131) | 56% (73/131) |
| III | 16% (29/186) MSIH<br>13% (24/186) MSIL<br>72% (133/186) MSS | 58% (108/186) | 56% (105/186) | 18% (34/186) | 9% (16/186) | 45% (83/186) |
| IV | 16% (7/44) MSIH<br>16% (7/44) MSIL<br>68% (30/44) MSS | 59% (26/44) | 57% (25/44) | 27% (12/44) | 16% (7/44) | 40% (17/42) |
| **Gender** | **P=2e-3** | **P=0.5** | **P=0.4** | **P=0.02** | **P=0.11** | **P=0.05** |
| Male | 14% (40/279) MSIH<br>15% (43/279) MSIL<br>70% (196/279) MSS | 63% (175/279) | 61% (169/279) | 19% (54/279) | 8% (21/279) | 46% (129/278) |
| Female | 29% (45/157) MSIH<br>13% (23/157) MSIL<br>59% (92/157) MSS | 59% (93/157) | 56% (88/157) | 30% (46/157) | 13% (20/157) | 57% (88/155) |
| **Age** | **P=2e-3** | **P=1e-3** | **P<1e-3** | **P=0.2** | **P=0.5** | **P<1e-3** |
| ≤65 | 12% (23/192) MSIH<br>16% (30/192) MSIL<br>72% (139/192) MSS | 101/192 (53%) | 95/192 (38%) | 38/192 (20%) | 7% (16/192) | 40% (76/192) |
| >65 | 26% (61/239) MSIH<br>13% (32/239) MSIL<br>61% (146/239) MSS | 163/239 (68%) | 158/239 (66%) | 61/239 (26%) | 11% (26/239) | 59% (139/237) |
| **Immune Subtype** | **P<1e-3** | **P=1e-3** | **P=4e-3** | **P=0.07** | **P=0.05** | **P<1e-3** |
| C1 | 14% (18/129) MSIH<br>19% (24/129) MSIL<br>67% (87/129) MSS | 66% (85/129) | 61% (79/129) | 21% (27/129) | 7% (9/129) | 50% (65/129) |
| C2 | 27% (57/210) MSIH<br>12% (25/210) MSIL<br>61% (128/210) MSS | 65% (135/210) | 62% (131/210) | 28% (58/210) | 13% (28/210) | 59% (123/210) |
| C3 | 0% (0/36) MSIH<br>6% (2/36) MSIL<br>94% (34/36) MSS | 31%<br>(11/36) | 31%<br>(11/36) | 6% (2/33) | 0% (0/33) | 8% (3/36) |
| C4 | 11% (1/9) MSIH<br>22% (2/9) MSIL<br>67% (6/9) MSS | 78% (7/9) | 78% (7/9) | 22% (2/9) | 0% (0/9) | 44% (4/9) |
| C5 | 0 | 0% (0/0) | 0% (0/0) | 0% (0/0) | 0% (0/0) | 0% (0/0) |
| C6 | 0% MSIH<br>0% MSIL<br>100% (7/7) MSS | 57% (4/7) | 43% (3/7) | 14% (1/7) | 0% (0/7) | 0% (0/7) |
| NA | 16% (8/49) MSIH<br>20% (10/49) MSIL<br>63% (31/49) MSS | 49%<br>(24/49) | 49%<br>(24/49) | 18% (9/49) | 6% (3/49) | 49% (24/49) |
| **TCGA Subtype** | **P<1e-3** | **P<1e-3** | **P<1e-3** | **P<1e-3** | **P<1e-3** | **P<1e-3** |
| EBV | 0 MSIH<br>20% (4/30) MSIL<br>87% (26/30) MSS | 57%<br>(17/30) | 53%<br>(16/30) | 7% (2/30) | 0% (0/30) | 43% (13/30) |
| HM | 93% (74/80) MSIH<br>3% (2/80) MSIL<br>5% (4/80) MSS | 99%<br>(79/80) | 99%<br>(79/80) | 79% (63/80) | 41% (33/80) | 100%<br>(80/80) |
| GS | 0 MSIH<br>4% (2/50) MSIL<br>96% (48/50) MSS | 36%<br>(18/50) | 32%<br>(16/50) | 6% (2/50) | 0% (0/50) | 14% (7/50) |
| CIN | 0.4% (1/223) MSIH<br>20% (44/223) MSIL<br>80% (178/223) MSS | 56%<br>(124/223) | 52%<br>(116/223) | 9% (20/223) | 1% (3/223) | 41%<br>(91/223) |

Worldwide, gastric cancer remains one of the top deadliest malignancies (11). Advanced gastric tumors are treated with surgery and chemotherapy with 5-year survival rates above 50% if the disease has not spread, and <10% if metastasis has occurred (12). The connections between therapy outcomes, the stroma, and collagens remains uncertain in stomach cancer. Physical properties of collagen fibers have been associated with outcomes in gastric cancer (GC) and these observations are likely driven by the most abundant collagen, type I (13). Collagen type I expression has been associated with metastasis in early onset gastric cancer (14).

To gain new insights into the function of collagens in cancer, we hypothesized that collagens are significantly mutated in tumors and that these mutations impact disease progression, therapy response, and patient outcomes. We further hypothesized that somatic mutations in collagens would resemble mutations observed in many collagenopathies, providing insights to function of collagens expressed and secreted from cancer cells. (15, 16). Patients with collagenopathies have both missense and truncation mutations that can be either dominant or recessive and demonstrate a range of penetrance depending on the mutation and collagen (17–20). Notably, the lessons from collagenopathies highlight how collagen mutations impact tissue even in the presence of wild-type collagen and even when collagens are expressed at low levels in the tissue.

There has been limited analysis of somatic mutations in the matrisome. COL2A1 has been reported to harbor recurring mutations in chondrosarcoma (21). Screening by MutSig2CV across 27 cancers identified COL11A1, COL13A1, COL19A1, COL1A2, and COL4A4 as borderline significantly mutated (22), and 2 collagens were significant in the TCGA stomach adenocarcinoma (STAD) (**Table S1**). Functional studies of these variants were not pursued in larger -omic screens in part due to “technical limitations”, as stated by the authors (22). COL14A1 was reported to have a nonsynonymous mutation rate of 4.4% in Microsatellite Stable (MSS) gastric tumors, (23). Grouping genes either by network methods (24) or by careful examination of specific gene families and mutation has provided insights into splicing regulators (25), TGF-β signaling (26), and complement genes (27), for example. Because there is some redundancy and overlap in function of collagens, we applied this approach to consider the collagens as a group, and to identify sets of collagens that may be significantly mutated and impactful in stomach cancer. We focused on collagens, as opposed to the whole matrisome to ease interpretation of the mutations, evaluate combinations, and leverage insights from collagenopathies to interpret the impact of mutations and propose specific hypotheses for future testing. A recent study by Izzi et al. also suggested that matrisome genes, including collagens, are significantly mutated across many cancer types (28). This study focused on reporting the existence of somatic mutations, enrichment of mutations in some protein domains, and identified individual genes associated with patient survival. However, Izzi et al. did not consider gene combinations, gene families, or using germline mutations to interpret the potential impacts of the somatic mutations. We also present a new approach to consider significant association with overall survival by defining a background set of genes considering mutation rate to identify collagen genes likely not associated with overall survival by chance. By focusing on collagens and leveraging the insights from collagenopathies, we also highlight how association with protein domains does not tell the whole story of the impact of somatic mutations of ECM factors.

In this article, we use bioinformatics to elucidate how collagen mutations affect gastric tumor outcomes. First, we find that many expressed collagens harbor somatic missense and truncation mutations, at a higher rate than expected compared to the background mutation rate. Next, we show that collagen genes and combinations of these genes associate with differential patient outcomes. We further investigate how collagen mutations correlate with tumor hallmarks, extracellular matrix components, and immune infiltration. Together, these findings suggest that collagen mutations impact stomach tumors via distinct tumor microenvironments, and that many collagens have unexpected novel functions in stomach tumors.

## Results

### Collagen mutations are prevalent in STAD

We evaluated the frequency of somatic mutations in the 43 human collagen genes. We observed a clear bias in the distribution of the frequency of mutations in collagens compared to other genes (p < 1e-16, Wilcoxon Rank test) (Figure 1A). Five individual collagen genes are mutated at frequencies larger than 8% (Figure 1B). Frequently mutated genes include COL12A1, COL11A1, COL6A2, and COL7A1, representing a range of collagen families and functions (see Table S1 for collagen family information). To account for the range of mutation rates in stomach tumors, we evaluated the MSS and MSIH types separately and found frequent somatic mutations in collagens in both MSIH and MSS tumors (Figure 1C). Some collagens such as COL12A1 and COL4A1 showed high mutation rates in both MSIH and MSS tumors, while others such as COL7A1 were frequently mutated in MSIH, but not MSS tumors. Every MSIH tumor has at least one mutation in a collagen gene. In MSS tumors, COL12A1 was the most frequently mutated at 8% with only 20 tumors harboring any collagen truncation mutation.

**Figure 1.**
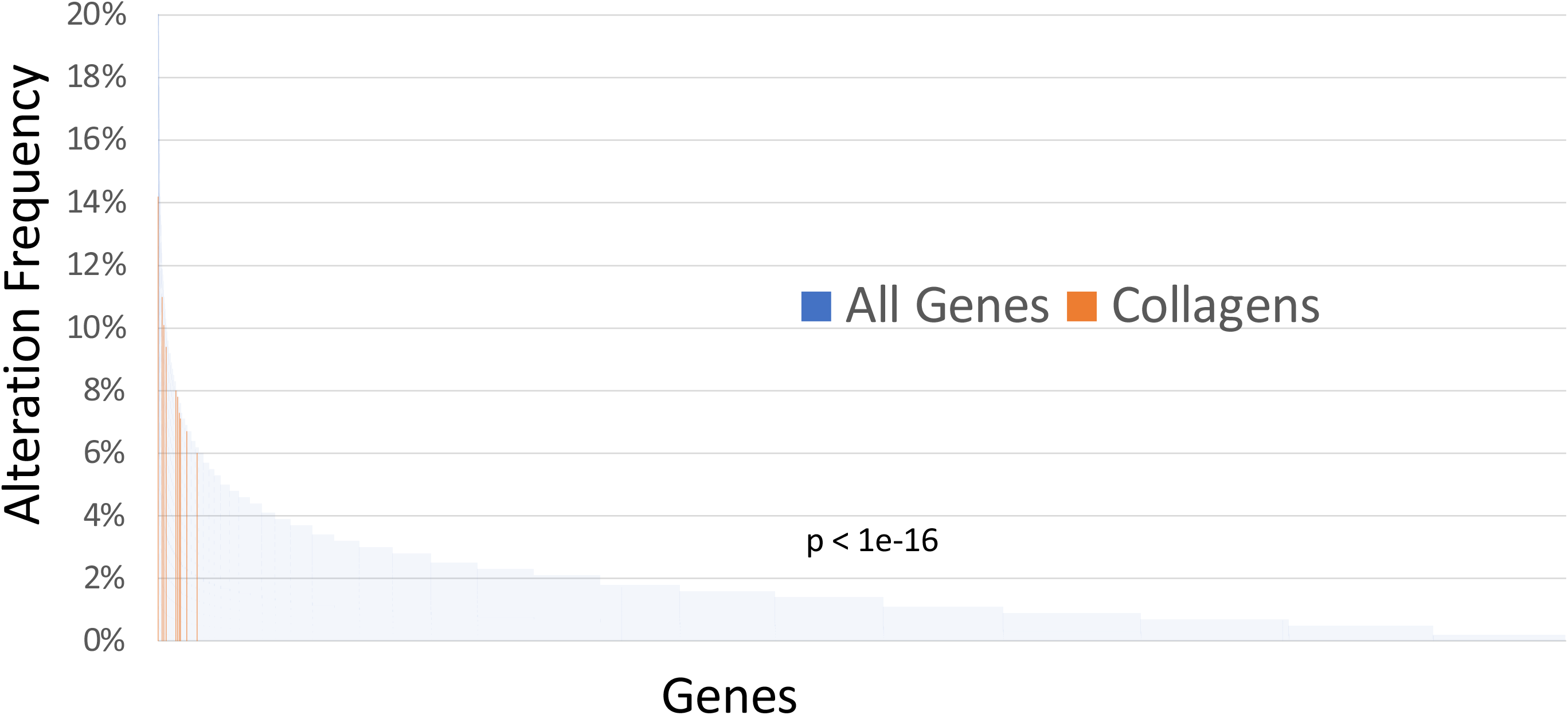
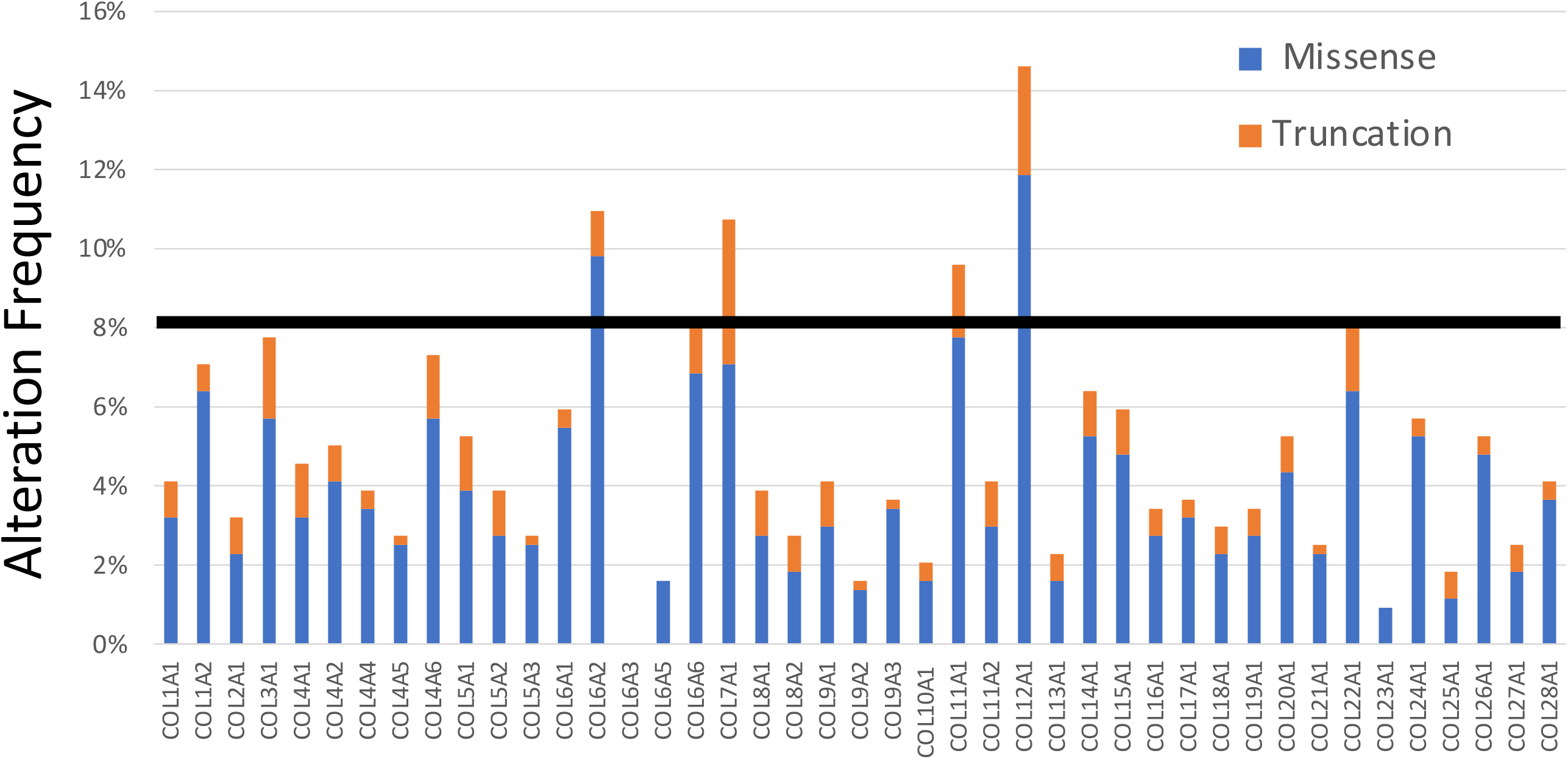
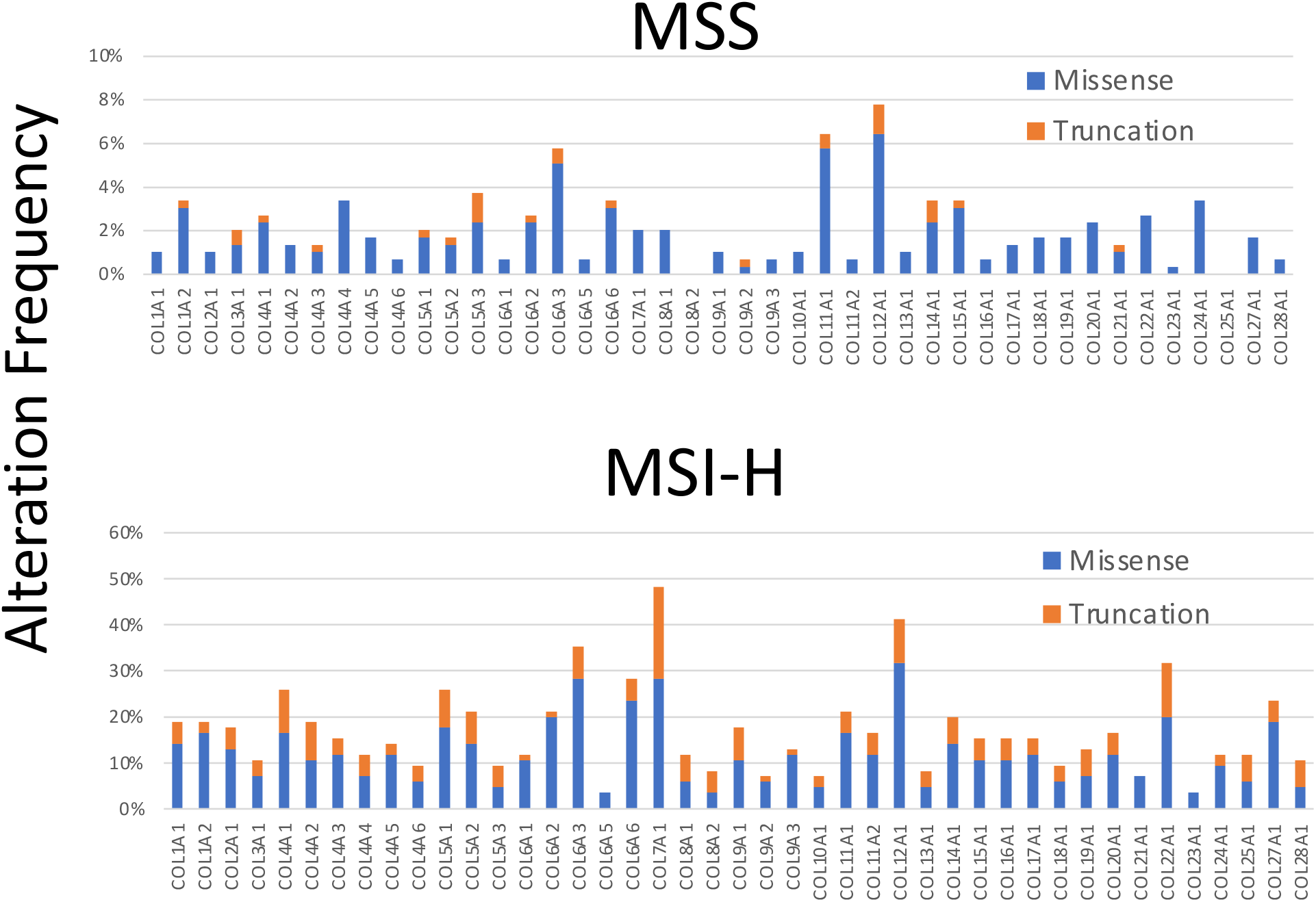
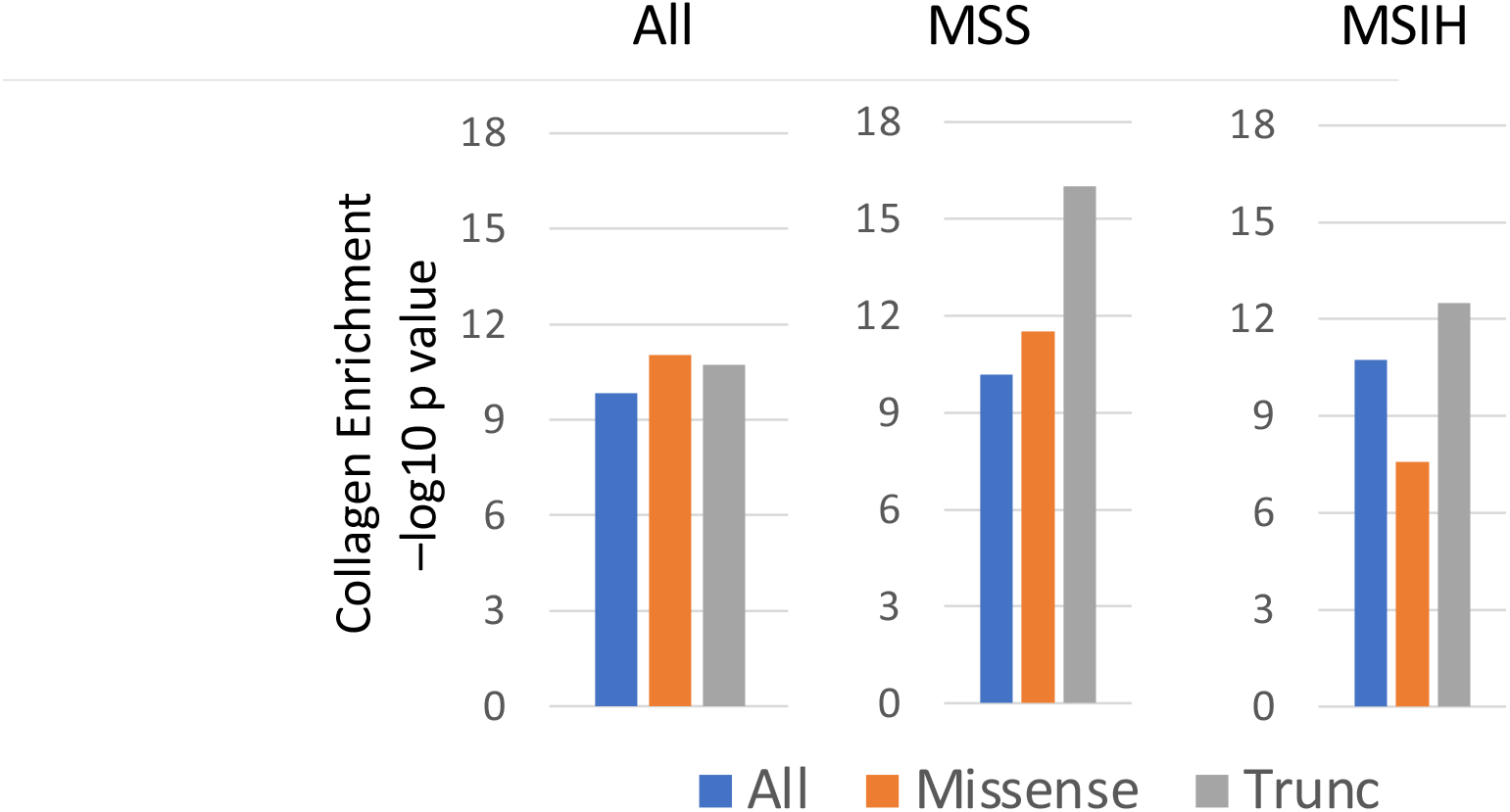
Collagens are significantly mutated in stomach adenocarcinoma in the TCGA dataset. **A.** Distribution of alteration frequencies for collagen genes (orange) compared to all other genes (blue) in the TCGA STAD cohort. P-value determined by Wilcoxon rank test comparing the distribution of collagen genes relative to all other genes. **B.** Alteration frequencies for each collagen gene in all TCGA STAD cases. **C.** and in MSS, MSIH, and MSIL STAD cases. **D.** Kolmogorov-Smirnov moderated tests suggest that collagen genes as a group are significantly mutated compared to gene sets of similar size and length in the whole TCGA STAD cohort and in both MSS and MSIH tumors.

Because collagen somatic variants are relatively rare and have not previously been identified as significantly mutated by standard algorithms, we evaluated if collagen genes were significantly mutated relative to the background mutation rate using multiple approaches. By MutSigCV2, only 2 collagen genes were significantly mutated in the TCGA gastric cancer cohort while 2 other collagens had borderline q-values (**Table S1**). To determine the significance of somatic mutations in collagens relative to other genes, accounting for mutation rate, we applied a modified Kolmogorov-Smirnov (KS) test (27). KS test analysis revealed that as a group, mutation rate of collagens, accounting for gene size, occurred significantly above background (Figure 1D). Because some GC tumors are MSIH with high mutation rates compared to MSS tumors, we determined that collagens and subsets of collagen genes had significantly higher mutation rates in these more specific cohorts as well (Figure 1D).

### Collagens are mutated at similar rates in independent datasets

We examined independent datasets to assess if other GC cohorts harbor similar mutations rates of collagens. 52% of tumors have at least one somatic mutation in a collagen in the Pfizer/Hong Kong whole genome sequencing dataset in 100 cases collected in Hong Kong, including a 19% rate of truncation mutations (29), but patient survival data is not available. The Asian Cancer Research Group (ACRG) performed targeted sequencing of 251 gastric tumors including a selection of collagens including COL11A1, COL12A1, COL21A1, COL22A1, COL4A1, COL5A1, COL5A3, COL6A3, and COL6A5 (30). Recurrent variants of the collagen genes tested were reported at frequencies slightly lower than observed in TCGA (**Figure S1**). Tumors harboring at least one mutation in COL11A1, COL5A1, COL5A3, COL6A3, COL6A5, or COL4A1 were moderately associated with improved outcomes in both TCGA and ACRG (**Figure S1B**). Major differences in the studies included that ACRG only sequenced patients of Asian ethnicity compared to TCGA including mostly Caucasians. Patients in the ACRG cohort had a longer overall survival (OS) than the TCGA patients (<50% survival vs. >60% survival at 5 years), and the MSI cases in the ACRG cohort showed a stronger association with improved outcomes compared to the TCGA cohort (**Figure S5A**).

### Collagen mutations associated with clinicopathological characteristics

Collagen mutations classify STAD tumors independently of clinicopathological characteristics including stage, grade, MSI status, and mutation rate (**Tables 1 and 2**). Age at diagnosis was associated with missense mutations and the overall mutation rate, consistent with MSIH tumors’ known association with older patients (31). Previous STAD classification identifies 4 major groups: Epstein-Barr Virus (EBV), High Mutation (HM), Genomically Stable (GS), and chromosome instability (CIN). Almost every tumor in HM, characterized by high mutation rates, has at least one mutation in a collagen gene, but also, 56% of the CIN and 36% of the GS groups have collagen mutations even though the mutation rates are much lower in these groups. Neither EBV nor H. Pylori status was associated with collagen mutations, or COL7A1 mutations (**Table 1**). These TCGA defined classifications were not associated with patient outcomes.

**Table 2.** Univariate and multivariate analysis by cox proportional hazards analysis. Multivariate survival analysis of all variables with p<0.05 in univariate analysis by cox proportional hazards analysis

| **Group** | **OS HR** | **OS P** |
| --- | --- | --- |
| All Collagens; all mutations | 0.8 (0.6-1.1) | 0.2 |
| All Collagens; missense only | 0.9 (0.7-1.2) | 0.6 |
| All Collagens; truncation only | 0.5 (0.4-0.8) | **0.002** |
| COL5A2; all mutations | 0.3 (0.1-0.8) | **0.02** |
| Grade | 1.3 (1.0-1.7) | 0.07 |
| Stage | **1.7 (1.4-2.0)** | **<1e-3** |
| MSI status (MSS vs. MSIH) | 0.7 (0.4-1.1) | 0.09 |
| Age at diagnosis | 1.0 (1.0-1.0) | **0.03** |
| Nonsilent mutation rate | 1.0 (1.0-1.0) | 0.06 |
| Stromal Fraction | **2.4 (1.1-5.3)** | **0.04** |
| Leukocyte Fraction | 1.5 (0.5-4.6) | 0.5 |
| TGF-beta Response | **1.1 (1.0-1.2)** | **0.004** |
| **MSIH only (N=68)** |  |  |
| COL14A1/COL5A3; all mutations | **3.2 (1.4-7.3)** | **0.006** |
| COL7A1/COL11A1; all mutations | **0.4 (0.2-0.9)** | **0.02** |
| Stage | 1.7 (1.0-2.9) | 0.06 |
| Stroma | 1.8 (0.2-20) | 0.6 |
| TGF-beta Response | **1.3 (1.0-1.5)** | **0.02** |
| Age at diagnosis | 1.1 (1.0-1.1) | 0.007 |
| Nonsilent mutation rate | 1.0((1.0-1.0) | 0.4 |
| MULTIVARIATE |  |  |
| **All cases (N=388)** |  |  |
| Stage | **2.0 (1.5-2.5)** | **<1e-3** |
| All Collagens; truncation only | **0.5 (0.3-0.9)** | **0.02** |
| Stromal Fraction | 1.8 (0.7 – 4.3) | 0.2 |
| **MSIH only (N=68)** |  |  |
| TGF-beta Response | **10.1 (2.8-37.5)** | **0.007** |
| Missense COL5A3 | **1.3 (1.1-1.6)** | **<1e-3** |
| **MSS Only (N=289)** |  |  |
| Stage | **1.7 (1.3-2.1)** | **<1e-3** |
| COL11A1, COL16A1, COL4A1, COL6A2 missense | **0.37 (0.2-0.8)** | **0.02** |

### Collagen mutations associated with patient survival

We evaluated the association of tumors harboring a somatic nonsynonymous missense or truncation mutation in any collagen with OS by Kaplan-Meier analysis in the TCGA STAD dataset (Figure 2A). Tumors with at least one mutation in any collagen were not associated with OS, but tumors with at least one truncation mutation in any collagen were significantly associated with longer OS (Figure 2A). Only COL5A2, COL11A1, and COL19A1 were significantly associated with longer OS when considered as individual genes (**Figure S1)**. COL23A1 was associated with shorter survival but is mutated in only 4 cases and is not significantly expressed in STAD (**Table S2**).

**Figure 2.**
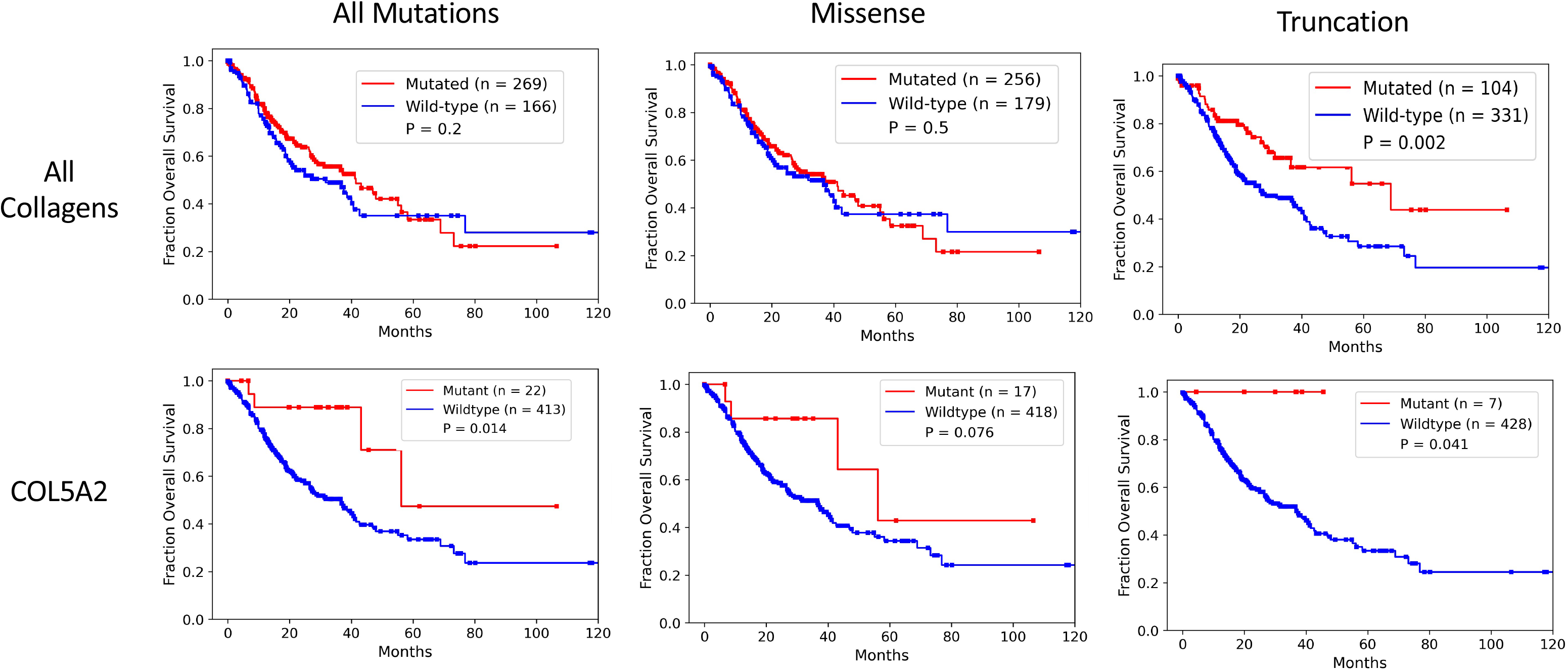
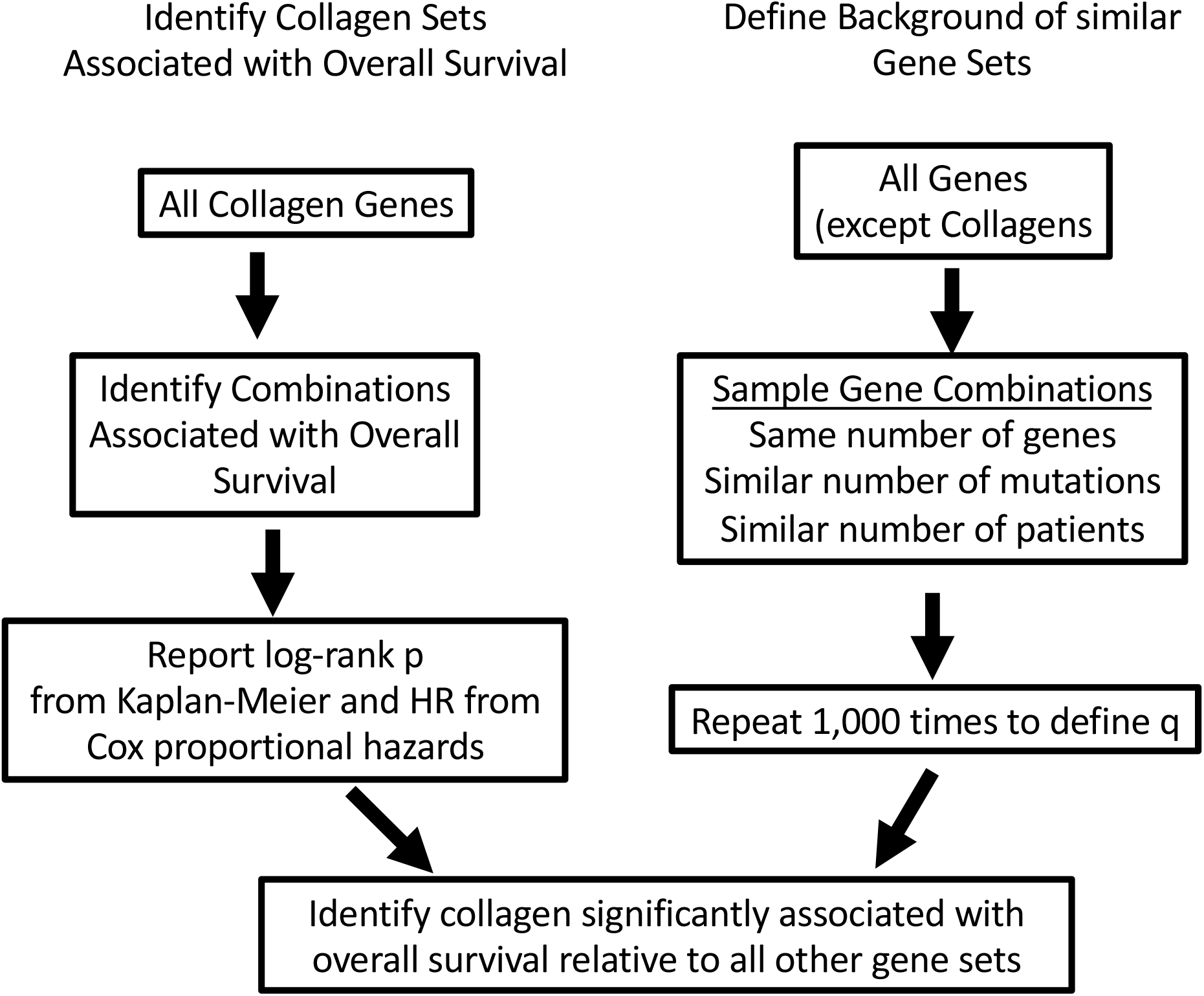
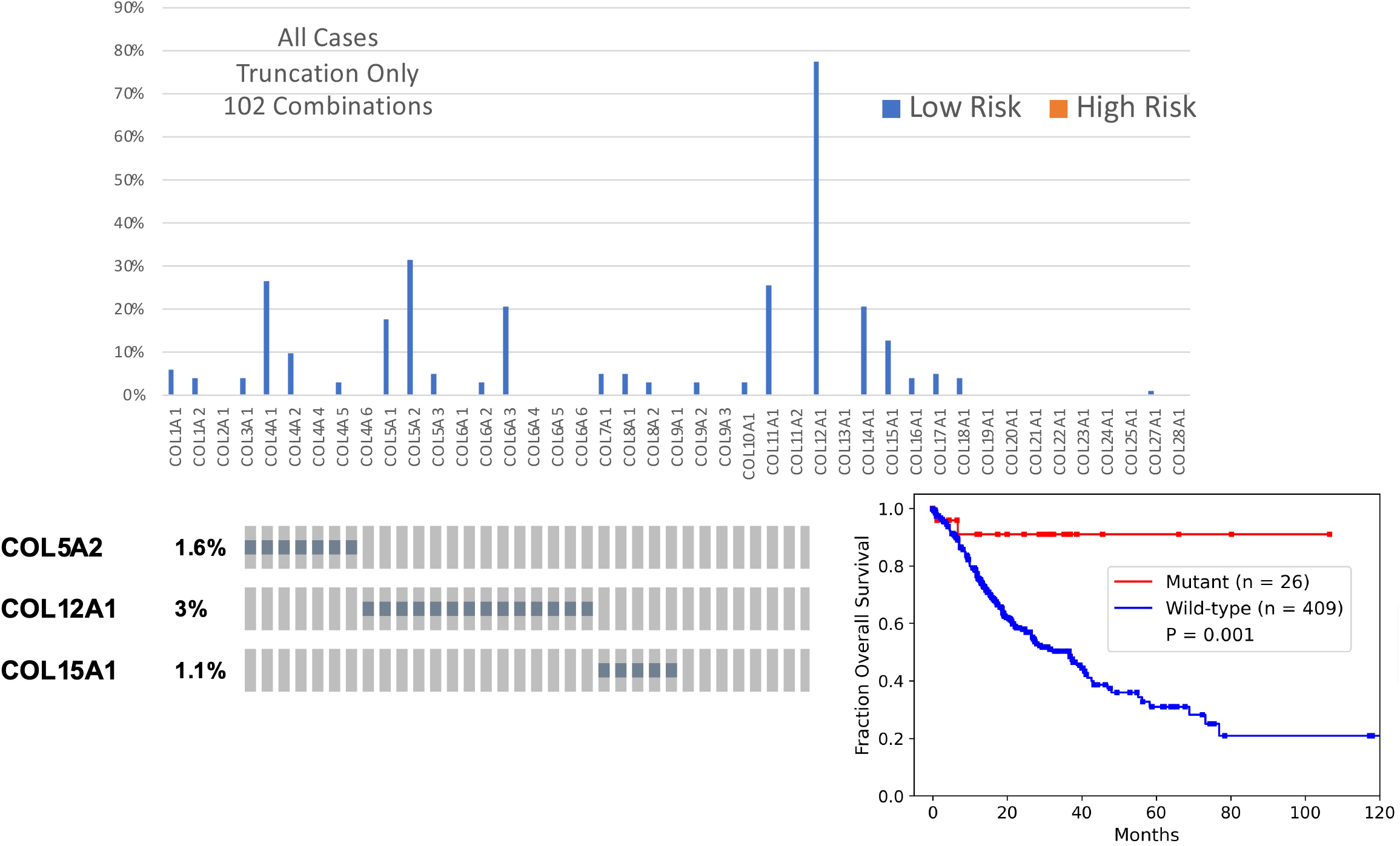
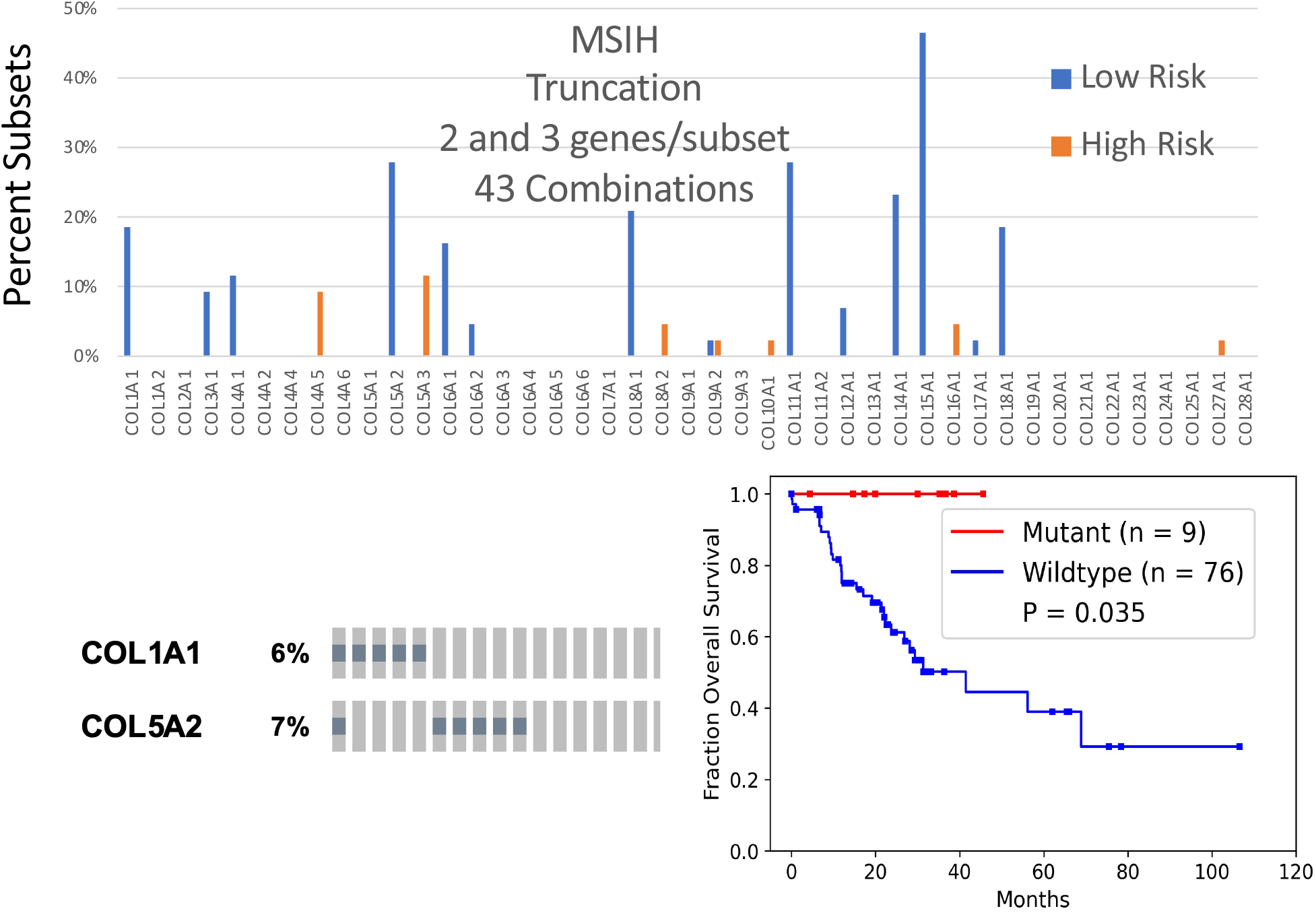
Identification of collagen genes mutations associated with overall survival. **A.** Patients with tumors that harbor at least one mutation in a collagen gene, have significantly better outcomes in the STAD TCGA cohort. Patients with tumors with at least one collagen mutation of the type indicated in red. Wild-type tumors in blue. Log-rank test p-values shown. Truncation mutations in any collagen gene were associated with better outcomes while nonsynonymous missense mutations were not associated with overall survival. Both missense and truncation mutations in COL5A2 were associated with longer overall survival. **B.** Schematic of approach to identify tumors with combinations of mutated collagens associated survival more significantly relative to background accounting for mutation rate, gene size and number of patients. **C.** Frequency of the inclusion of each collagen gene with a truncation mutation in a combination significantly associated with overall survival. A representative combination of collagen genes strongly associated with overall survival curve and the oncoprint. **D.** Identification of collagen genes with truncation mutations in MSIH tumors most strongly associated with overall survival. Frequency of the inclusion of each collagen in subsets consisting of 2 and 3 collagen genes with truncation only mutations in MSIH tumors.

To address the potential redundancy of collagen functions and since many collagens were not mutated in sufficient number of cases for survival analysis, we undertook a combinatorics approach to identify sets of tumors with mutated collagen genes associated with OS more significantly than a bootstrap defined background accounting for number of patients, mutation rate and number of genes (Figures 2B). We evaluated all combinations of tumors with at least one mutation in 2 or 3 expressed collagen genes (**Table S3**) and tested their association with OS. The combinatorics approach identified collagen gene sets associated with OS at q≤0.05 (Figure 2C). COL5A2, COL4A1, COL11A1, COL15A1, and COL16A1 were the most frequently included collagen genes in the combinations (**Figure S3A**), and each at least trended with longer OS on their own (**Figure S2A**). Truncation mutations were particularly strongly associated with OS with no combinations identified associated with shorter OS at these thresholds (Figure 2A). Of the 104 tumors with at least one truncation mutation, 65% were in MSIH tumors (**Table 1**). We identified combinations of 2 or 3 collagen genes with truncation mutations strongly associated with OS and COL12A1 was the most collagen gene most frequently included in these combinations (Figure 2C, **Table S3**).

### Collagen Genes Classify MSIH Tumors by Overall Survival

Because the majority of collagen mutations were in MSIH cases, and because of the differences in MSIH and MSS stomach tumors in treatment response, we evaluated each of these groups separately. For simplicity and to observe differences between MSIH and MSS differences more clearly, we removed the MSIL annotated tumors to avoid complications from these tumors with moderate mutation burden that are clinically treated as MSS tumors. Even in just MSIH cases, most truncation mutations in collagens were associated with longer survival including in COL1A1, COL5A2, COL11A1, and COL15A1 (Figure 2D). A few collagens were associated with shorter survival; including most notably, COL5A3. All patients with either a COL1A1 or COL5A2 truncation in MSIH tumors were associated with longer OS (Figure 2D). In MSS tumors, COL5A3, COL6A2, COL11A1, and COL24A1 were associated with longer OS (**Figure S3C**).

Combining the top truncation variants identified by combinatorics defined a group of tumors strongly associated with longer OS in MSS and MSIH tumors (Figures S3E and S4). These observations suggest that loss of expression of many collagens, especially those involved in collagen type I expression and formation (see Table S1), such as COL12A1, was associated with increased OS in both MSS and MSIH tumors. On the other hand, the loss of function of some collagen type I regulating collagens, such as COL5A3 and COL14A1, is detrimental to patients with MSIH tumors. These observations can be explained by considering that COL5A3 and COL14A1 are negative regulators of collagen type I fiber size. LOF mutations in these two collagens lead to gain of function of collagen type I. COL14A1 is a Fibril Associated Collagens with Interrupted Triple helices (FACIT) collagen that regulates collagen fibrillogenesis such that absence of COL14A1 leads to larger fibers in mice (32–34). Increasing collagen type I fiber width has been associated with poor outcomes in stomach cancer (13). Analogous to the observations in stomach tumors, mutations of collagen α3(V) chains have phenotypes distinct from collagen α1(V) and α2(V) chains in mice (35). Germline mutations in COL5A1 and COL5A2 cause Ehlers-Danlos-like phenotypes, while mutations in COL5A3 affect adiposity (35). On the other hand, LOF mutations in other regulators of fiber formation such as COL11A1 (36) were associated with longer OS (**Figure S2**). These observations suggest that regulation of COL1A1 and fibrillogenesis, when unchecked, leads to even worse survival in MSIH cases. These examples demonstrate how we can leverage observations of collagen biochemistry and collagenopathies to interpret the impact of mutations and generate novel hypotheses to begin to explain the range of treatment responses and patient survival.

Many collagen gene combinations’ associations with OS were specific for MSIH or MSS tumors (**Table S3**, **Figure S4**, Figure 3). Representative sets were associated with OS in either MSIH or MSS tumor, but not both (Figure 3). In particular, even though COL5A3 and COL14A1 were mutated at similar levels, these collagens showed MSI status dependent associations with OS.

**Figure 3.**
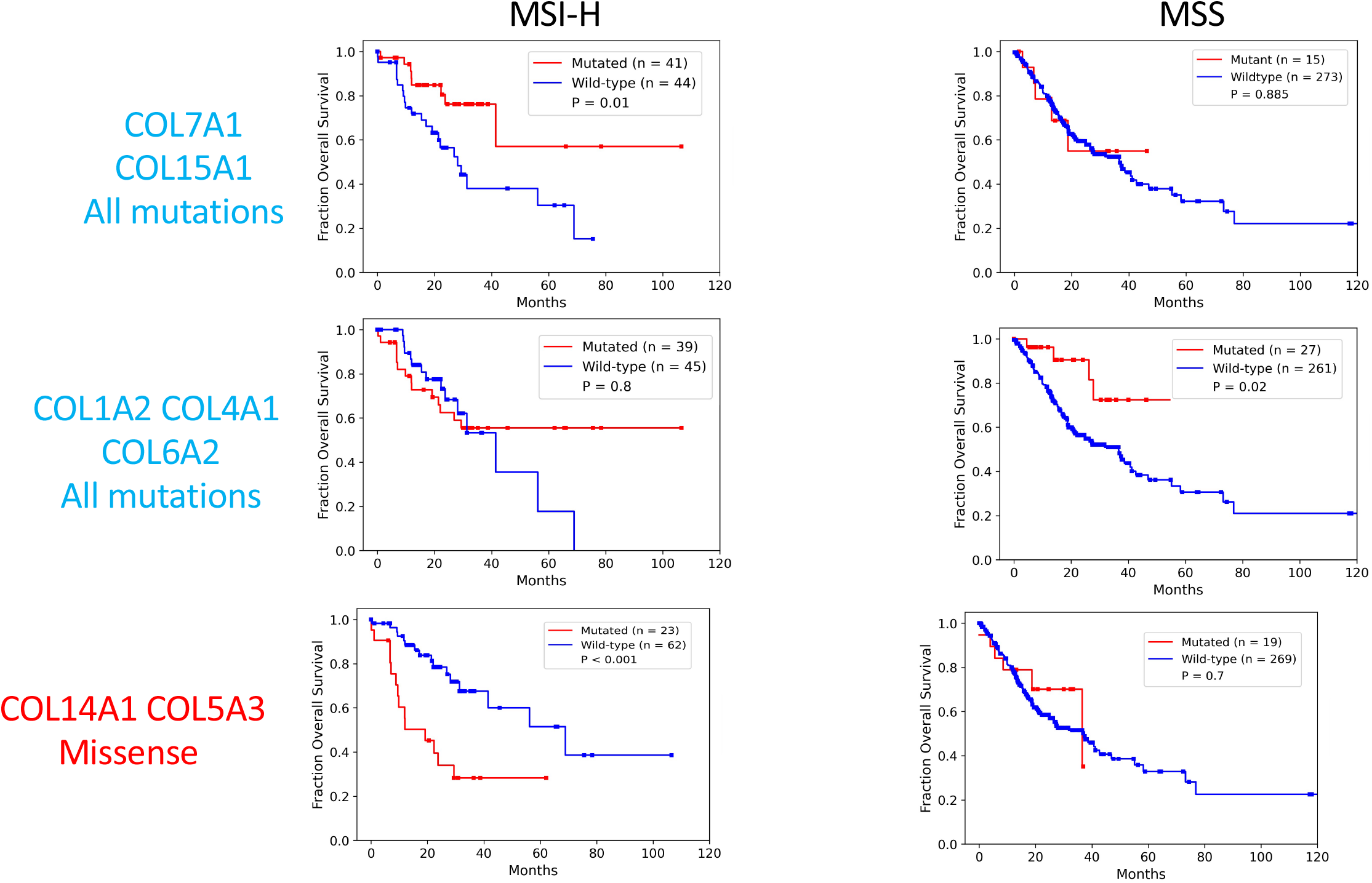
Specific collagen mutation combinations have context dependent association with overall survival in MSIH and MSS tumors. Kaplan-Meier survival analysis of representative collagen mutation combinations with differing patterns of association with overall survival in MSIH and MSS tumors. P-values determined by a log-rank test.

We applied the cox proportional hazards model to test the relationship of collagen mutations with other common survival associated characteristics including age and stage. Multivariate analysis showed that collagen mutations were independent predictors of OS compared to other clinicopathological characteristics including stage (**Table 2**). Neither mutation rate nor MSI status were associated with OS, despite including many collagen mutations. These findings suggest that tumors with collagen mutations specifically define a class of tumors with distinct properties and treatment responses.

### Collagen mutations impact on stomach tumors

To gain insight into how collagens could be affecting STAD tumors, we used pre-ranked Gene Set Enrichment Analysis (GSEA) of TCGA normalized RSEM scores to identify biological processes associated with tumors that harbor a collagen mutation compared to tumors without collagen mutations. We first evaluated the 50 MSigDB hallmark gene sets (37) (Figure 4). There was high similarity of the impact of collagen mutations on expression of cancer hallmarks highlighted by the higher expression of cell cycle drivers including E2F targets and MYC gene sets (Figure 4A, B). On the other hand, EMT, KRAS and myogenesis gene sets were expressed higher in wild-type compared to collagen mutant tumors (Figure 4A, B). Lower expression of the EMT hallmark in tumors with collagen mutations is consistent with reduced collagen function and an altered ECM that leads to more epithelial features in these tumors (38).

**Figure 4.**
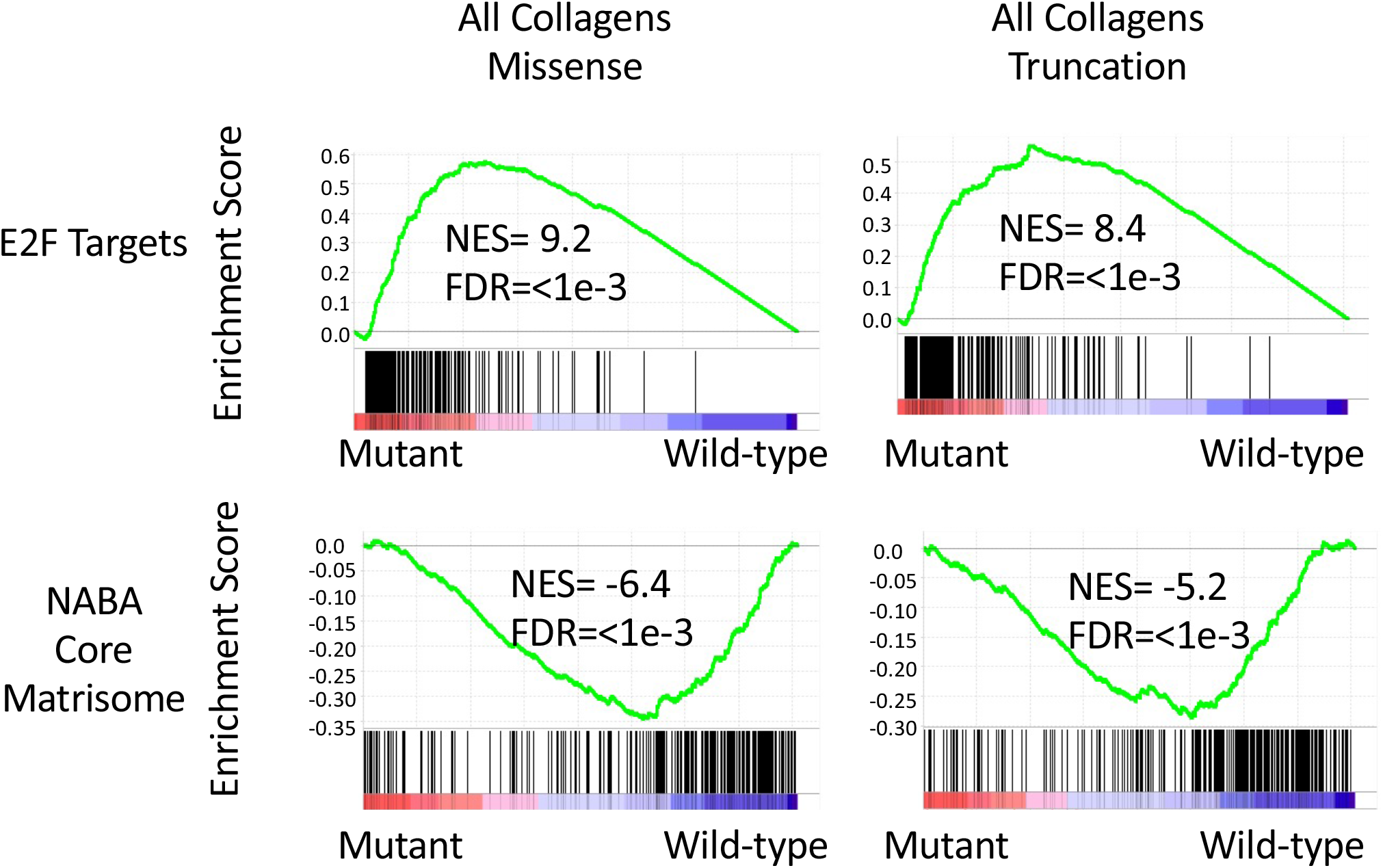
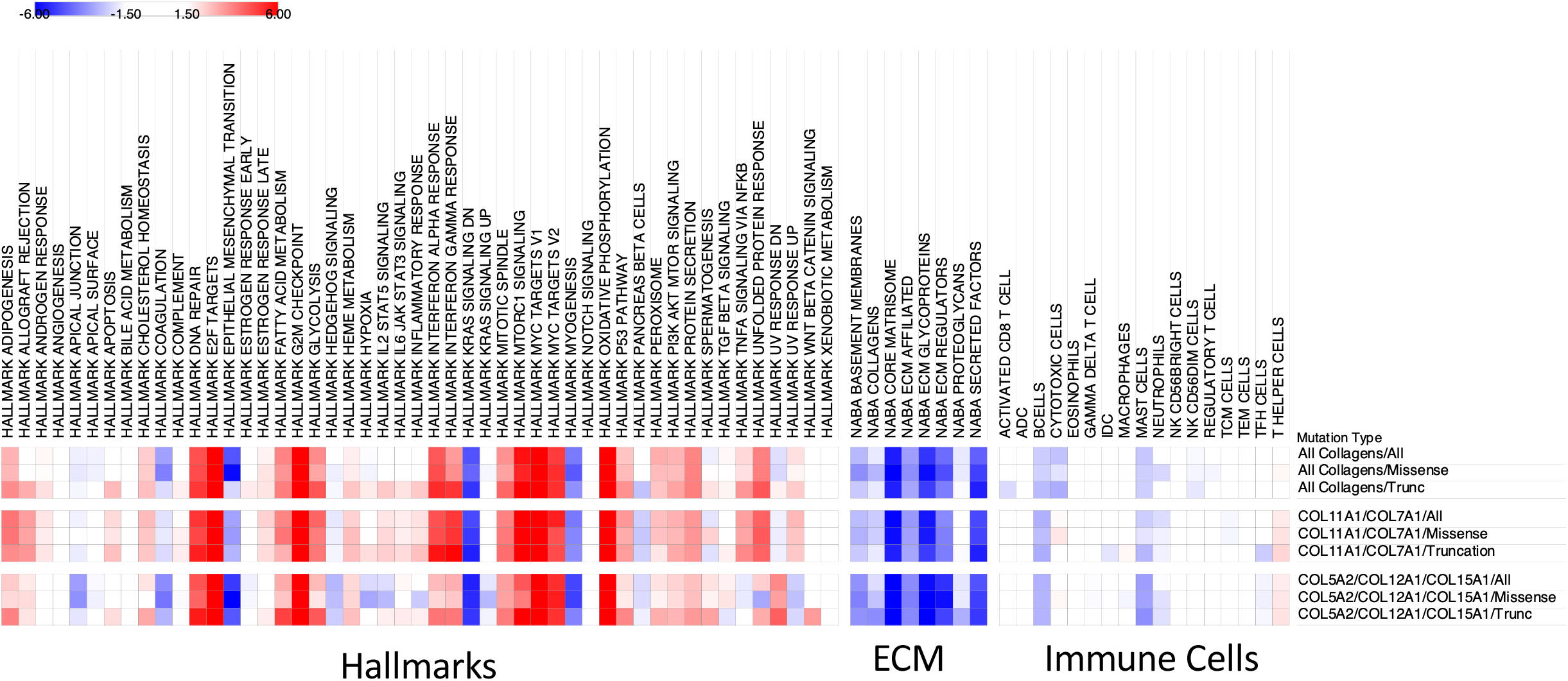
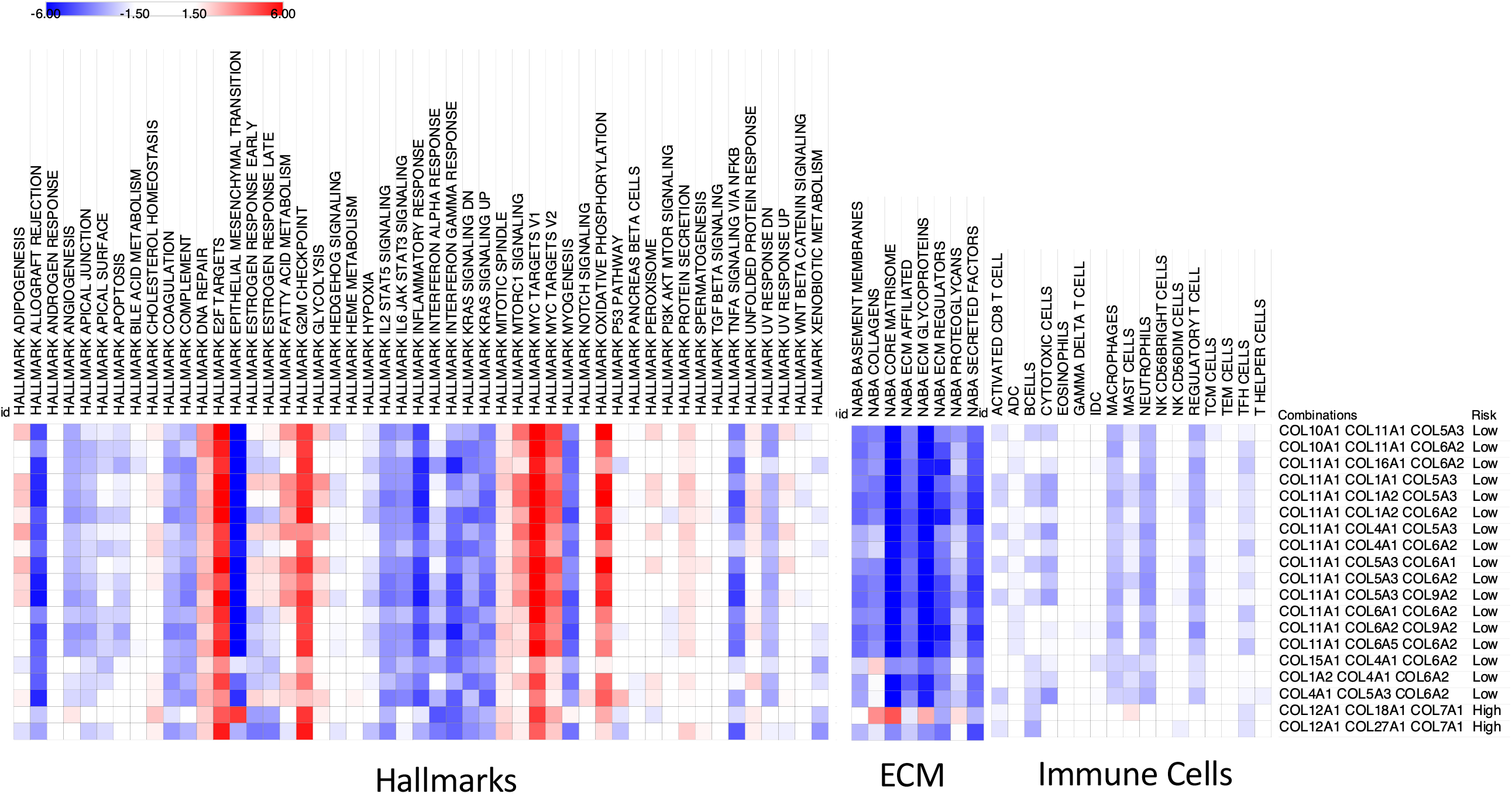
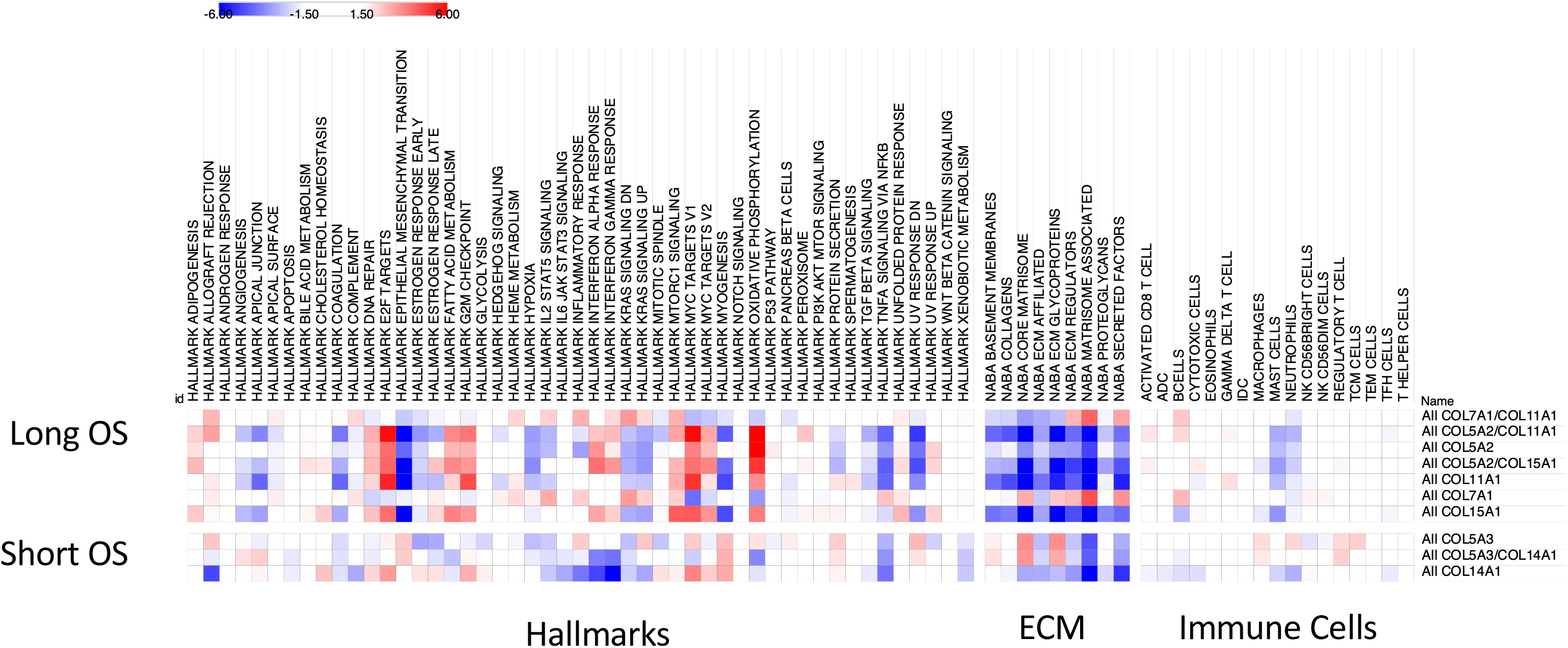
Tumors with collagen mutations have distinct expression of cancer hallmarks and tumor environments. **A.** Representative enrichment plots from pre-rank GSEA suggest upregulation of E2F regulated transcripts and down-regulation of the expression of the matrisome in tumors with collagen mutations. TCGA stomach tumors were classified by collagen mutation status and pre-ranked GSEA revealed associations with the indicated gene set. **B.** Heat map of normalized enrichment scores of the cancer hallmark, immune cell, and NABA ECM gene sets. Red indicates higher expression in mutant tumors and blue indicates higher expression in wild-type tumors. Nonsignificant and modest enrichment scores between −1.5 and 1.5 are in white. In the full STAD TCGA cohort, tumors with any collagen mutation or with only mutations in COL7A1 or COL11A1 showed similar patterns. **C.** Heatmap of gene sets in MSS only tumors. **D.** Heatmaps of gene sets in MSIH only tumors reveal a more diverse pattern of enrichment. All heatmaps generated in Morpheus (61).

To test if the ECM was different in tumors with collagen mutations, we evaluated the NABA ECM gene sets. All the NABA gene sets were expressed lower in tumors with mutations compared to wild-type tumors when considering the full TCGA STAD cohort, consistent with a disrupted ECM in the collagen mutated tumors relative to the wild-type tumors (Figure 4B). Total collagen expression has been associated with patient outcomes in many cancers (39). Although collagens as a group were expressed lower in mutant tumors, only rarely was the expression directly associated with mutation within that collagen gene (**Table S4**). This may be because many of the collagens are expressed from both cancer and stroma cells, obscuring the relationship between collagen mutation and expression in bulk RNAseq data.

### Collagen mutations associated with distinct tumor microenvironments

Collagens can mediate the migration and infiltration of immune cells in tumors (40). To evaluate association between collagen mutations and immune cell infiltration, we evaluated immune cell signatures (41) with pre-ranked GSEA. MSIH tumors have higher expression of most of the immune cell expression signatures compared to MSS tumors, consistent with more immune cell infiltration in MSIH tumors (**Figure S5**). Together with lower expression of the NABA ECM gene sets in MSIH tumors (**Figure S5**), these observations suggest that the MSIH and MSS tumors differ in their tumor microenvironments. Immune cell gene expression signatures were expressed higher in MSIH tumors compared to MSS tumors, consistent with a more inflammatory environment and higher tumor mutation burden in MSIH compared to MSS tumors. Because of these large differences in the tumor environments of MSS and MSIH tumors, considering all the stomach tumors together may be obfuscating impacts. We therefore evaluated the impact of collagen mutations in the whole cohort as well in MSIH and MSS tumors separately.

Combinations of collagen mutations had a more consistent impact on the expression of ECM and immune cell gene signature in MSS tumors, compared to more variable associations in MSIH tumors (Figure 4 **and Figures S6-S10**). Figure 4 shows representative combinations and pre-ranked GSEA of all combinations listed in Table S3 are shown in Figures S6-S10. The consistent nature of the tumors with collagen mutations suggests that changes in EMT, expression in basement membranes, and many immune cells, are common features of tumors with collagen mutations in both MSS and MSIH tumors. This consistency could be because we selected for tumors with collagen mutations associated with overall survival and these tumors have similar mechanisms of impact in mediating EMT and expression of the basement membrane.

In MSS tumors, tumors with collagen mutations had consistently lower expression of extracellular matrix gene sets and the majority of the immune cell gene sets (Figure 4C). While in MSIH tumors, the ECM NABA and immune cell signature gene sets were split into 2 groups (Figure 4D). Notably, tumors with COL14A1 and COL5A3 mutations were associated with higher expression of NABA gene sets in MSIH tumors and shorter OS (Figure 4D). On the other hand, other fibril associated collagens such as COL11A1 and COL5A2 were associated with longer OS (**Figure S3**, **Table S4**). COL11A1, COL5A1, and COL5A2 promote fibril formation and loss of function mutations of these collagens have been associated with smaller collagen type I fibers (35). Mutations in COL1A1, COL11A1, COL5A1, and COL5A2 were all associated with lower expression of the EMT hallmark gene set compared to wild type in MSIH tumors, while mutations in COL14A1 and COL5A3 were associated with higher expression of EMT expression signature in MSIH tumors (Figure 4D). These observations predict that tumors with mutations in collagen types XIV and Vα3 would have thicker fibers, promoting cell migration, and subsequent tumor cell escape. On the other hand, tumors with in COL1A1, COL11A1, COL5A1, and COL5A2 are predicted to have thinner or fewer collagen type I fibers leading to less migration, lower mesenchymal properties, less metastasis, and higher sensitivity to treatments.

### Immunoenvironment

We used the immune cell gene sets reported by Tamborero et al. to evaluate the immunoenvironment by pre-rank GSEA (41). Across the whole cohort, B and mast cells were lower in tumors with a variety of mutant collagens, while T helper cells were modestly increased in some mutant collagen tumors (Figure 4A). In MSS tumors, Mast cells, macrophages, neutrophils were lower in mutant tumors associated with longer survival, and higher in wildtype tumors (Figure 4B). Mast cells and neutrophils have been associated with short OS in STAD (42, 43). When considering just the MSIH tumors, neutrophils and mast cells were lower in mutant tumors, but no other clear patterns emerged across the cohort. Immunosuppressive cell types including macrophages and regulatory T cells are higher in many of the shorter OS mutation combinations in both MSS and MSIH tumors. Tumors with COL14A1 mutations for example had higher levels of macrophages, neutrophils and mast cells Mast cells have been associated with shorter OS in stomach cancer (42). These observations, based on molecular signatures, suggest changes to the immunoenvironment in tumors with collagen mutations.

### COL7A1 mutations

We hypothesized that insight into the functional impact of mutations can be gained by comparing the pattern and type of mutation to those observed in collagenopathies. As an example, we focused on COL7A1 which is the mutational cause of Dystrophic Epidermolysis Bullosa (DEB) and had significant associations with patient outcomes in MSIH tumors (Figure 2B). COL7A1 germline mutations were downloaded from a DEB mutation database (44). The distribution of germline and stomach somatic mutations were very similar (Figure 5B). A Kruskal-Wallis test (*P*=0.3) suggested that the two distributions were not significantly different. COL7A1 mutations in STAD were slightly more biased towards the N-terminus of the protein compared to DEB. The largest exon, exon 73, was most frequently mutated in both the germline and cancer mutations. The recurring nonsense mutation at position 2029 is the same hotspot observed in DEB patients (Figure 5A and B). The distribution of somatic mutations resembled the distribution of DEB germline mutations suggesting that no unusual tumor specific mutation pattern is prevalent in stomach tumors. Moreover, because the type of mutation is similar as observed in DEB, we can infer the function of the somatic mutations in tumors.

**Figure 5.**
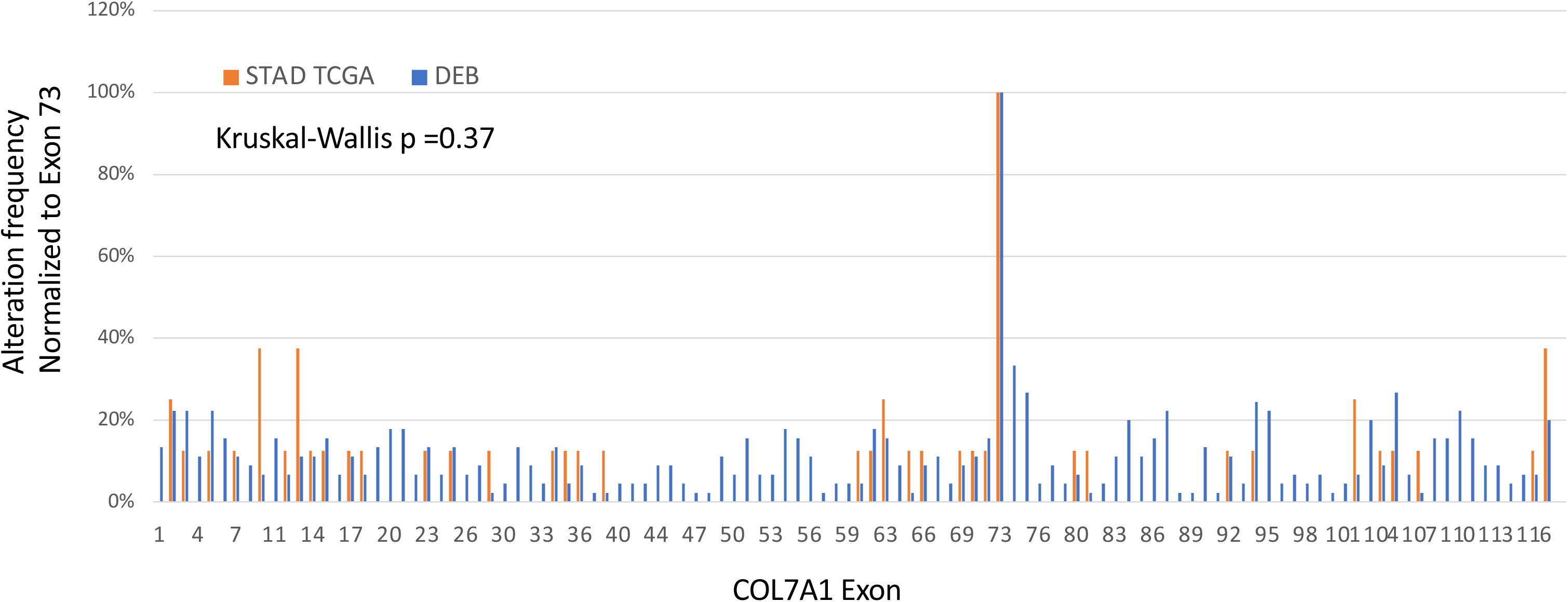
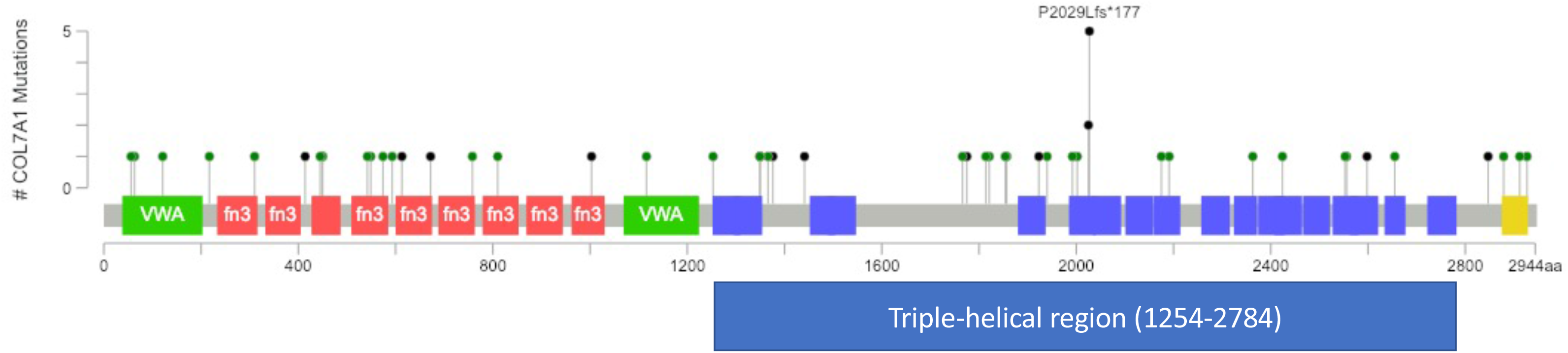
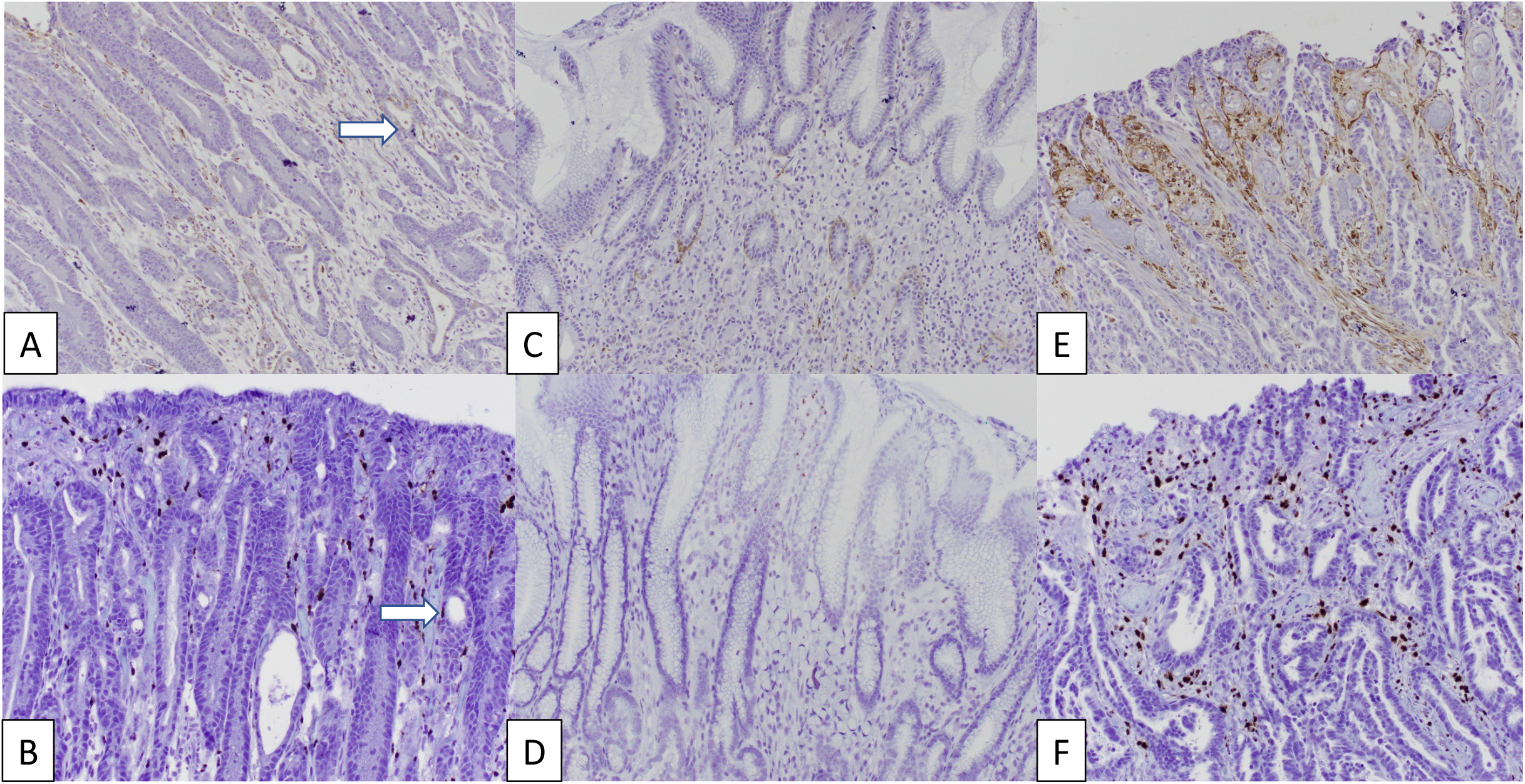
COL7A1 somatic mutations resemble inherited germline mutations found in collagenopathies. **A.** Distribution of somatic variants in TCGA STAD is similar to the germline variants observed in DEB as determined by Kruskal-Wallis test. Mutations in the N-terminal domain often reduce COL7A1 expression in skin (63, 64). **B.** Lollipop plot showing the distribution of variants on the COL7A1 protein domain map. A recurring truncation variant is found in the collagen domain in exon 73. Other variants only were observed once or twice, but have redundant impacts in each domain.

### Expression of COL7A1 in stomach tumors

Many minor collagens have a high tissue specificity including COL7A1 (45). Because COL7A1 is not known to be expressed in normal stomach, or anywhere in the gastrointestinal tract, we wanted to confirm that COL7A1 was actually expressed in stomach tumors, and importantly determine if COL7A1 was expressed in cancer cells and not just in the stroma. We evaluated COL7A1 protein expression by immunohistochemistry in a set of 10 stomach tumors from patients treated at Rhode Island Hospital (Figure 5C **and S11**). COL7A1 was expressed in the stroma (4/10), in the epithelium (3/10) or was not detectable (3/10) (**Table S5**). In the epithelium, COL7A1 was expressed in the cytoplasm, similar to skin cells highly expressing COL7A1 (Figure S11C), suggesting that these tumor cells are also expressing and secreting high levels of COL7A1. Expression from tumor cells was confirmed by *in situ* hybridization using RNAscope, controlling for non-specific antibody staining (Figure 5C). Other collagens have been shown to be expressed in tumor cells by *in situ* hybridization such as collagen type IV (9). Hynes and colleagues have suggested that matrisome components secreted from tumors cells in pancreatic tumors are more impactful than those originating from the stroma (46). Larger studies are needed to evaluate any connection between COL7A1 expression patterns and mutation status.

## Discussion

This work demonstrates that collagen mutations are significant relative to background, stratify patients in meaningful ways, and are likely impactful in STAD. Although this is an association study, we believe that this report will inspire additional investigation into the function of collagens in tumors and specifically in stomach cancer.

In collagenopathies, many causative collagen mutations are heterozygotes and genetically dominant because the missense mutation forms destabilized triple helices. A second class of missense mutations in non-collagen domains and truncation mutations both reduce or eliminate expression of the collagen. Compared to tumors secreting wild-type versions of these collagens, stomach tumors with reduced levels of these collagens respond better to treatment and have distinct expression of multiple cancer hallmarks.

Izzi and co-workers recently reported on somatic mutations in the matrisome including collagens PanCancer (28). Their PanCancer approach focused on identifying common features across multiple cancer types while this study focused on the presence and potential impacts of mutations in stomach cancer. They also identified COL5A2 and COL15A1 associated with longer OS in STAD but did not consider combinations or accounting for MSI status. This study reports the presence of collagen mutations in both high and low mutation burden tumors and their differential impacts. Unlike Izzi et al., in the case of collagens in STAD, we did not observe a correlation between RNA expression levels and mutations. This is likely because of the contribution to collagen expression from myriad cell types in tumors. We also leverage the vast knowledge from germline mutations to interpret the tumor somatic mutations. Together, the report from Izzi et al. and this report, using different methods to account for mutation burden, provide data emphasizing the importance of somatic mutations in ECM components.

We hypothesized that comparing somatic mutations in tumors to germline mutations in collagenopathies would be informative and aide interpretation of the somatic mutations including the recessive or dominant nature of the mutation and the mechanisms that the mutation impacts collagen structure, subsequent tumor hallmarks and response to treatment. For example, COL12A1 was the collagen gene with the highest truncation mutation frequency and most often strongly associated with shorter OS in the combinatorics analysis (Figure 2C). COL12A1 is a FACIT homotrimer collagen that mediates interactions between collagen type I and the rest of the ECM. COL12A1 is reported to be expressed by both stroma and tumor cells in STAD (47–50) and is expressed in gastric cancer cell lines (51). Germline mutations, including rare splicing mutation truncation variants, in COL12A1 cause a Ehlers-Danlos/Bethlem-like myopathy syndrome (52, 53). Together, these observations suggest that COL12A1 is a critical determinant of STAD disease progression and therapy response and exemplifies how comparison of somatic and germline mutations aids interpretation and provides new insights into tumors.

There are two general patterns of collagenopathy mutations also observed in STAD: mutations that disrupt the collagen triple helix such as Glycine mutations and mutations outside the protein domains known to cause reduced expression of the protein including both missense and truncation variants. These are the major mechanisms that cause multiple collagenopathies. As highlighted, these types of mutations are observed in COL7A1 and COL12A1. Similar variants are observed in other collagens such as COL1A1 and collagen type V (Figure S12).

Collagens form two of the major structures in the ECM: the basement membrane and the interstitial ECM. Both of these ECM components are impacted by collagen mutations. The loss of integrity of the basement membrane in tumors suggests a disorganized, more porous structure that could cause increased inflammation, analogous to COL7A1 mutations in DEB, or with collagen type IV and type VI variants (16). Collagen type I, expressed by the COL1A1 and COL1A2 genes, plays a critical role in forming the ECM and organizing cell-cell interactions and mechanical properties in tumors (54). Increased collagen type I has been associated with worse outcomes in many cancers including stomach (13, 55). Both COL1A1 and COL1A2 had modest mutation frequencies with only weak association with OS in STAD (Figure 1**; Figure S2**). However, truncation mutations of COL1A1 were associated with longer survival and most truncation mutations of COL1A2 were also associated with longer survival except for 1 case, TCGA-HU-A4GQ-01, which was reported to have deceased at 0 months, and therefore may be reflecting other causes of death. Collagen type I missense variants were not associated with OS, perhaps because the majority of collagen type I originates from the stroma and therefore any impact of a mutated collagen type I originating from tumor cells may be diluted. Collagens that interact with collagen type I and regulate fiber size and structure including all 3 collagen type V genes, COL11A1, COL12A1, and COL14A1 have significant mutation rates and association with OS. These observations further support the concept that loss of collagen type I from the tumor cells, or dysregulation of the network that forms collagen type I dependent structures affects patient outcomes.

Altogether, these observations suggest the regulation of collagen type I and the basement membrane by a panoply of cancer cell secreted collagens are critical for tumor fate. Collagens, as one of the dominant structural proteins in the ECM play myriad functions in regulating cancer hallmarks (56). Minor collagens such as COL7A1 and COL12A1 form structural links between the collagen type I fiber network and/or the basement membrane zone. These data support a model where a local ECM derived from components secreted from the cancer cells, reshape the local ECM and are critical for tumor phenotypes, including EMT, drug response, the immunoenvironment and overall disease progression (Figure 6). In tumors with wild-type collagens, EMT is higher, collagen type I fibers are wider, and higher expression of the matrisome including the basement membrane compared to tumors with mutant collagens. On the other hand, some mutant MSIH tumors, associated with shorter OS, exemplified by missense COL5A3 and COL14A1 which regulate COL1A1, have higher expression of mesenchymal genes, wider collagen type I fibers, and a different immune cell infiltration pattern (Figure 4D). Linking hallmarks and pathways to dysregulated ECM caused by collagen mutations may lead to new opportunities to refine drug targeting and development.

**Figure 6.**
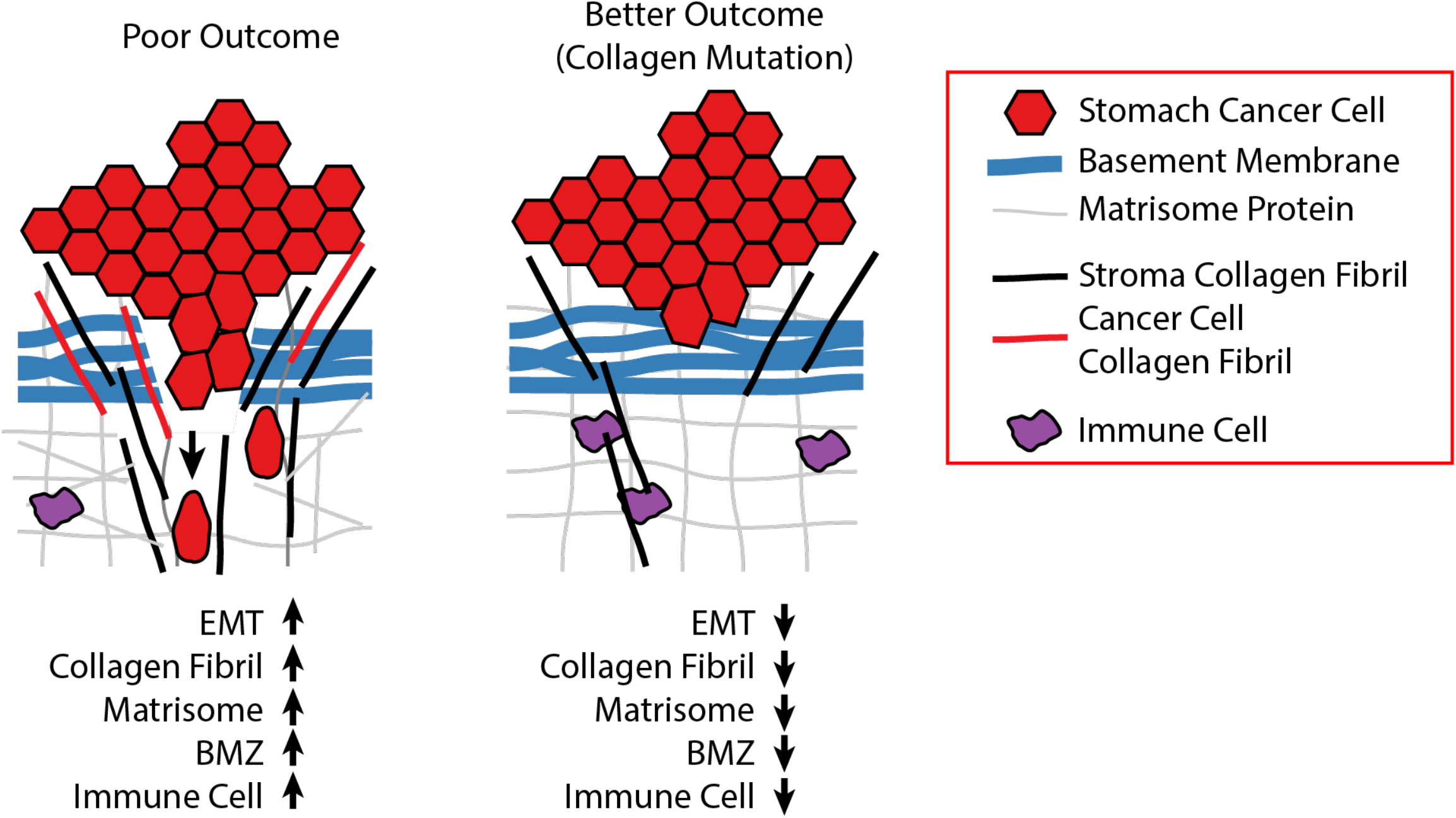
Model of impact of cancer cell secreted collagens on tumors. Collagens originate from either the cancer or stroma cells. Truncation and missense collagen mutants reorganize the tumor microenvironment decreasing multiple processes that increase drug sensitivity and reduce metastasis risk including reduced EMT, less local collagen around the cancer cells, a more disorganized collagen structure, and increased infiltration of cytotoxic immune cells and drugs.

## Conclusions

In conclusion, we find a high frequency of individual collagen genes and sets of collagen genes harboring somatic mutations compared to background in both microsatellite stable (MSS), and microsatellite instable (MSIH) stomach adenocarcinomas in TCGA and comparable datasets. Overall, combinations of somatic mutations are predictive of patient survival, and truncation mutations associate with improved survival. We further associate these combinations with distinctive tumor microenvironments based on lower matrisome expression, cell cycle and EMT, as well as immune cell infiltration. Interestingly, stomach cells express COL7A1, normally associated with skin ECM, and somatic mutations in COL7A1 predict improved overall survival. It should be noted that this study is limited by the dependence on genotype-phenotype correlations in patients so that there is risk in oversimplifying rare mutations. Some of this risk is ameliorated by the combinatorial approach and the interpretation of the mutations based on similar variants observed in collagenopathies. Nevertheless, many of these missense mutations resemble the loss of function (LOF) mutations in collagenopathies, which give some insight into their potential role in tumor progression. Overall, this study suggests the further testing of collagen mutations in stomach cancer is promising and collagen mutations could be incorporated into strategies to classify cancer patients.

## Abbreviations

ACRG: Asian Cancer Research Group
CIN: Chromosome Instability
DEB: Dystrophic Epidermolysis Bullosa
EBV: Epstein-Barr Virus
ECM: Extracellular Matrix
EMT: Epithelial mesenchymal transition
FACIT: Fibril Associated Collagens with Interrupted Triple helices
GDC: Genomic Data Commons
GSEA: Gene Set Enrichment Analysis
GS: Genomically Stable
HM: High Mutation
MSIH: Microsatellite Instable
MSS: Microsatellite stable
OS: Overall Survival
RSEM: RNA-seq by Expectation Maximum
STAD: Stomach Adenocarcinoma
TCGA: The Cancer Genome Atlas

## Methods

### Data sources

The Cancer Genome Atlas (TCGA) Pan-Cancer RNA-seq V2 normalized gene expression and clinical data was downloaded from Firebrowse in April 2018. TCGA somatic mutation data file, mc3.v0.2.8.PUBLIC.maf, was downloaded from the Genomic Data Commons (GDC) (57). Microsatellite data was downloaded from Firebrowse (STAD.merged_only_auxillary_clin_format.txt). Immune gene sets input into GSEA were defined by Tamborero et al. (41). Stroma scores, and overall mutation rates were derived from Table S1 by Thorsson et al (58). Hallmark (59) and NABA (60) gene sets were downloaded from MsigDB v7.0 (37). For the analysis of collagen mutations in the ACRG, collagen mutation data was obtained from the supplemental data published by Cristescu et al. (30). Truncation mutations include all variants predicted to cause a shorter protein or degrade the mRNA including nonsense mutations, frameshift mutations and mutations that affect splicing. Germline collagen mutations were obtained from the Leiden Open Variation Database, except for COL7A1. COL7A1 pathological mutations were obtained from a DEB mutation database (44).

### Software and statistical tests

Analyses were performed using R and python custom scripts. GSEA version 2.4 was run on either a Unix or MacOS system. Statistical tests were performed using the Lifelines v0.25.1 and SciPy v1.5.2 libraries in Python. Moderated Kolmogorov-Smirnov test was adopted from Olcina et al. to assess significance of collagen somatic mutations relative to other genes (27). Morpheus was used to generate the heatmaps (61). Survival curves were generated using cBioPortal’s oncoprinter web app and matplotlib v3.3.1. Lollipop plots were generated via the MutationMapper tool in cBioPortal.

### Identifying collagen gene combinations

We aimed to identify sets of collagen genes significantly associated with overall survival, accounting for gene size and mutation rate. To correct for multiple combinations occurring by chance, we calculated a q value for a given subset of collagen genes. To determine background, genes were randomly chosen until the expected number of mutations were within 5 of the number of observed mutations in collagen genes. A survival analysis was performed on the subset of patients used in the collagen subset analysis where the indicator variable was based on whether a patient has a mutation in the randomly chosen subset in at least 5% of cases of the designated cohort. We considered subsets with collagen genes significantly expressed with an average RSEM > 200. If a combination of 2 collagens was identified, this combination was not considered in combinations of 3 collagens. We then counted the frequency of each collagen included in the subsets as an indication of the contribution of each collagen to overall survival risk and exclusivity with the other collagens.

### Case selection for immunohistochemistry

With institutional review board approval, IRB #1070389-9, 10 cases of gastric adenocarcinoma diagnosed from 2010 to 2019 were retrieved from the archives of the Department of Pathology and Laboratory Medicine at Lifespan Academic Medical Center (Providence, RI).

### Immunohistochemistry

Immunohistochemistry staining for COL7A1 was performed on 4-μm paraffin sections. After incubation at 60°C for 30 minutes, the sections were deparaffinized and rehydrated with xylene and graded alcohols. Antigen retrieval was performed with Ready-to-Use Proteinase K (Agilent, Santa Clara, CA) incubating at 37°C for 10 minutes. The slides were then incubated with anti-COL7A1 antibody (1:5000) for overnight at 4°C. The immunoreactivity was detected by using the DAKO Envision + Dual Link System and the DAKO Liquid 3,3’-diaminobenzidine (DAB+) Substrate Chromagen System (Agilent, Santa Clara, CA). Immunohistochemistry was assessed by 2 pathologists (MR and EW).

### In situ hybridization

mRNA expression was determined using ISH with the RNAscope Assay (Advanced Cell Diagnostics, Hayward, CA). The ISH staining for COL7A1 was performed on 4-μm paraffin sections. After baking slides at 60°C for 1 hour and deparaffinizing FFPE sections with xylene, RNAscope® 2.5 HD Reagent Kit was used for the ISH assay. All the steps were done according to the kit protocol. After pre-treating the sample with hydrogen peroxide solution, heat target retrieval and protease plus, COL7A1 probe was added for 2hr at 40°C, sequentially hybridize with AMP 1, AMP 2, AMP 3, AMP 4, AMP 5, and AMP 6 reagents, for 30, 15, 30, 15, 60, 15 min, respectively. ISH signal was detected by the application of a chromogenic substrate. Tissue was counter-stained with haematoxylin. Scrambled negative control probes showed no signal.

### Antibody sources

Rabbit polyclonal anti-COL7A1 targeting the human LH7.2 domain was a kind gift from Alexander Nystrom, University of Freiburg (62).

## Declarations

### Ethics approval and consent to participate

Use of patient material was approved by the Lifespan institutional review board approval, IRB #1070389-9.

### Consent to publish

Not applicable

### Availability of Data and Materials

All genomic data used in this study is publicly available.

### Author Contributions

Conceptualization, ASB; Manuscript writing: ASB, IW; Methodology: ASB, IW, JK, EDG Coding: ASB, JK, KSG. IHC and pathology: DY, EW, ASS, MJR. Data interpretation, ASB, JK, KSG, MJR, EDG, EW, ASS, IW. All authors have read and approved the manuscript.

## Acknowledgements

We thank Dr. Alexander Nystrom for providing the anti-COL7A1 antibody. We are very grateful for our funders for providing support these last years. We thank the patients and their families for their participation in the individual TCGA projects.

## List of Additional Files

**Additional File 1.** Table S1

**Additional File 2.** Table S2

**Additional File 3.** Table S3

**Additional File 4.** Table S4

**Additional File 5.** Table S5

**Additional File 6.** All Supplemental Figures

## Supplemental Tables

### Additional File 1

**Table S1**. MutSig 2CV v3.1 analysis of significantly mutated collagen genes in STAD TCGA cohort. Data downloaded from Firebrowse.

### Additional File 2

**Table S2**. Average expression level of each collagen gene in the STAD TCGA cohort. Values are RSEM.

### Additional File 3

**Table S3**. Collagen gene combinations identified from combinatorics approach.

### Additional File 4

**Table S4**. Summary of COL7A1 protein expression in stomach tumors from Rhode Island Hospital as assessed by immunohistochemical staining.

### Additional File 5

## Supplemental Figure Legends

**Figure S1. Alteration frequencies of collagens in ACRG and HK/Pfizer datasets. A.** Alteration frequencies of sequenced collagens in other stomach cancer cohorts. **B.** Kaplan-Meier analysis of COl11A1, COL5, COL4, COL6 mutations in the ACRG targeted sequencing dataset compared to the same set of collagen genes in the TCGA cohort.

**Figure S2. Survival analysis of somatic mutations in each collagen gene. A.** Kaplan Maier analysis of tumors with any type of mutation in each collagen gene across the whole STAD TCGA cohort. Tumors with the designated collagen mutation are in red. Wild-type tumors are in blue. P-values determined by log-rank test. **B.** Kaplan Maier analysis of tumors with truncation mutations in each collagen gene across the whole STAD TCGA cohort. **C.** Kaplan Maier analysis of tumors with any type of mutation in each collagen gene in MSIH cases. **D.** Kaplan Maier analysis of tumors with truncation mutation in each collagen gene in MSIH cases.

**Figure S3. Identification of combinations of collagens genes associated with overall survival relative to background. A.** All mutations across the whole TCGA cohort. **B.** Representative examples of combinations of 2 collagens associated with overall survival. **C.** Combinations of all mutations in MSS tumors only. **D.** Combinations of all mutations in MSIH tumors only. **E.** Example of collagen genes with truncation mutations most frequently associated with overall survival when combined, classify MSIH tumors into high and low overall survival risk.

**Figure S4. Collagen mutations have MSIH and MSS context dependent differences in overall survival.** Mutations in COL5A3 and COL14A1 have different associations in MSIH and MSS tumors even though the total number of mutations is similar.

**Figure S5. MSIH and MSS tumors have distinct microenvironments in TCGA. A.** MSI status was associated with outcome in ACRG but not in TCGA. **B.** Comparison of MSIH and MSS stomach tumors by pre-ranked GSEA reveals differences in expression. Each heatmap plots the Normalized Enrichment Scores (NES) from the GSEA. NABA ECM gene sets were expressed higher in MSS tumors compared to MSIH tumors. Many immune cell expression signatures including cytotoxic cells were expressed higher in MSIH tumors compared to MSS tumors. B cells were expressed higher in MSS tumors. The majority of cancer hallmark expression signatures were expressed significantly higher in MSIH tumors compared to MSS tumors.

**Figure S6. Pre-ranked GSEA of collagen mutation combinations in Table S3 for the whole TCGA STAD cohort shows consistent impact for each mutation combination. A.** Hallmarks for combinations with both missense and truncation mutations. **B.** The NABA and immune signature genes sets for combinations with both missense and truncation mutations. **C.** Hallmark, NABA, and immune signature gene sets for combinations with just truncation mutations. **D.** Clustering of hallmark gene sets for tumors with missense mutations only in the whole TCGA cohort showed significant difference for the EMT hallmark relative to overall survival. P-value calculated by Kolmogorov-Smirnov.

**Figure S7. In MSS cases, pre-ranked GSEA of tumors with either a missense or truncation mutation combination as listed in Table S3 show impact of collagen mutations on pathways some of which are correlated with overall survival. A.** For all mutations in MSS cases, some hallmarks such as EMT were associated with overall survival as shown in the heat map and box plot. P-value calculated by Kolmogorov-Smirnov. **B.** NABA ECM and immune signature gene sets in MSS tumors. Basement membrane and macrophage signature gene sets were among the gene sets most associated with overall survival, showing consistent downregulation in tumors with mutant collagens and higher expression in wild-type tumors. P-value calculated by Kolmogorov-Smirnov.

**Figure S8. In MSIH cases, pre-ranked GSEA of tumors with either a missense or truncation mutation combination as listed in Table S3 show impact of collagen mutations on pathways some of which are correlated with overall survival. A.** Clustering of hallmark gene sets partitions tumors with collagen combinations by overall survival. Box plot shows the significant difference in the EMT hallmark as defined by combinations associated with high or low risk of overall survival. **B.** NABA gene sets showing large differences in Basement Membrane and ECM Affiliated gene sets relative to overall survival. **C.** Immune cell signature gene sets showing large difference in Tregs and Macrophage expression signatures. P-value calculated by Kolmogorov-Smirnov.

**Figure S9. In MSIH cases, pre-ranked GSEA of tumors with only missense mutation combinations as listed in Table S3 show impact of collagen mutations on pathways some of which are correlated with overall survival. A.** Hallmark gene sets. **B.** NABA ECM sets. **C.** Immune cell gene signatures. P-value calculated by Kolmogorov-Smirnov.

**Figure S10. In MSIH cases, pre-ranked GSEA of tumors with only truncation mutation combinations as listed in Table S3 show impact of collagen mutations on pathways some of which are correlated with overall survival. A.** Hallmark gene sets. **B.** NABA ECM and immune cell signature gene sets. P-value calculated by Kolmogorov-Smirnov.

**Figure S11. COL7A1 is expressed in some tumor cells in STAD. Representative images of COL7A1 protein and RNA expression in stomach adenocarcinoma. A.** Immunohistochemistry (A, C, E) and in situ hybridization (B, D, F) for COL7. Stromal localization in C, E, D, and F, and mixed stromal and carcinoma localization (at white arrows; A, B). **B.** Higher magnification of panels A and B from S7A showing expression by IHC in panel A and ISH in panel B of COL7A1 in epithelial regions. The arrow shows ISH signal in tumor cells. **C.** Representative images at higher power of COL7A1 protein expression by IHC in the epithelium and stroma. **D.** Representative image of COL7A1 protein expression in normal human skin. Note the line of expression in the ECM between the dermal and epidermal layers (red arrow). Cells expressing COL7A1 show cytoplasmic signal as they are overexpressing COL7 to be secreted to form the “anchorage” line.

Figure S12. Comparison of pathological germline and somatic mutations in STAD in three collagens. Comparison of the distribution of mutations across each gene and a lollipop plot mapping the somatic mutations to the protein domains. A. COL1A1. B. COL4A1. C. COL5A1.

