## Supplemental File 1 for "Somatic Mutations in Collagens are Associated with a Distinct Tumor Environment and Overall Survival in Gastric Cancer"

**Table S1. Collagen family for collagens of interest in stomach cancer**

| **Structural Family** | **Gene** | **Putative Role** | **Genetic Disease** | **References** |
| --- | --- | --- | --- | --- |
| **Fibril forming** | COL1A1 | Fiber collagen | Osteogenesis imperfecta; Ehlers-Danlos | (1) |
|  | COL1A2 | Fiber collagen | Osteogenesis imperfecta; Ehlers-Danlos | (1) |
|  | COL2A1 | Fiber collagen | spondyloepiphyseal dysplasia congenita | (2) |
|  | COL3A1 | Fiber collagen | Vascular Ehlers-Danlos | (3) |
|  | COL5A1, COL5A2 | Promotes Type I fibers | Classic Ehlers-Danlos | (4) |
|  | COL5A3 | Negative regulator of Type I fibers | N/A |  |
|  | COL11A1 | Promotes Type I fibers | Sickler Syndrome;  Marshall Syndrome; skeletal dysplasia | (5, 6) |
|  | COL11A2 | Promotes Type I fibers | Sickler Syndrome  Marshall Syndrome  Otospondylomegaepiphyseal dysplasia | (7) |
|  | COL14A1 | Fibril surface; Negative regulator of Type I fibers | Keratoderma | (8) |
|  | COL24A1 | Type I fibrillogenesis regulator |  |  |
|  | COL27A1 |  | Steel syndrome | (9) |
| **FACIT** | COL9A1 | Crosslinked to fibrils | Multiple epiphyseal dysplasia; osteoarthritis; Stickler syndrome | (7) |
|  | COL9A2 | Crosslinked to fibrils | multiple epiphyseal dysplasia | (7) |
|  | COL9A3 | Crosslinked to fibrils | Multiple epiphyseal dysplasia Autosomal recessive Stickler syndrome | (7) |
|  | COL12A1 | Basement membrane | myopathy | (10, 11) |
|  | COL15A1 | Banded Fibril linker | glaucoma | (12) |
|  | COL19A1 | Basement membrane zones |  |  |
|  | COL20A1 |  | N/A |  |
|  | COL21A1 |  | N/A |  |
|  | COL22A1 | Basement membrane zones | N/A |  |
| **Network** |  |  |  |  |
|  | COL4A1 | Basement | Familial porencephaly Hereditary angiopathy with nephropathy, aneurysms and muscle cramps syndrome | (13) |
|  | COL4A2 | Basement | Familial porencephaly Hereditary angiopathy with nephropathy, aneurysms and muscle cramps syndrome | (13) |
|  | COL4A3 | Basement | Alport Syndrome | (14) |
|  | COL4A4 | Basement | Alport Syndrome | (14) |
|  | COL4A5 | Basement | Alport Syndrome | (14) |
|  | COL4A6 | Basement | N/A |  |
|  | COL8A1 | Basement | N/A |  |
|  | COL10A1 | Chondrocyte matrix deposition | Schmid type metaphyseal chondrodysplasia; spondylometaphyseal dysplasia | (15) |
| **COL6** |  |  |  |  |
|  | COL6A1 | Basement membrane/interstitial matrix | Bethlem myopathy, Ullrich congenital muscular dystrophy | (16) |
|  | COL6A2, COL6A3 | Basement membrane/interstitial matrix | Ullrich congenital muscular dystrophy | (16) |
|  | COL7A1 | Dermoepidermal Anchoring fibril | Dystrophic epidermolysis bullosa (DEB) | (17) |
|  | COL26A1 |  | N/A |  |
|  | COL28A1 |  | N/A |  |
| **Membrane** | COL13A1 | Not known function | Congenital myasthenic | (18) |
|  | COL17A1 | Dermoepidermal anchoring complex | Junctional epidermolysis bullosa-other | (19) |
|  | COL23A1 | Not known function | N/A |  |
|  | COL25A1 | Linked with amyloid formation | congenital cranial dysinnervation disorder | (20) |
| **Multiplexins** | COL18A1 |  | Knobloch syndrome; glaucoma | (12) |
