## Supplemental File 2 for "Somatic Mutations in Collagens are Associated with a Distinct Tumor Environment and Overall Survival in Gastric Cancer"

**Supplemental Table S2. Collagen genes significantly mutated in the TCGA STAD cohort as assessed by MutSigCV2.**

| **Gene** | **p-value** | **q-value** |
| --- | --- | --- |
| **COL12A1** | **1.1e-4** | **0.006** |
| **COL11A1** | **0.001** | **0.04** |
| **COL20A1** | **0.004** | **0.09** |
| **COL8A1** | **0.014** | **0.2** |
