## Supplemental File 5 for "Somatic Mutations in Collagens are Associated with a Distinct Tumor Environment and Overall Survival in Gastric Cancer"

**Table S5. Immunohistochemical staining for COL7 in gastric adenocarcinoma tumor cells and tumor-associated stroma.**

| **Case number** | **COL7 staining in stroma** | **COL7 staining in carcinoma** | **Histologic grade and features** |
| --- | --- | --- | --- |
| 1 | Weak positive staining | Negative | Moderately to poorly differentiated, mixed intestinal and diffuse types |
| 2 | Negative | Negative | Poorly differentiated, intestinal type |
| 3 | Negative | Multifocal, weak staining in carcinoma | Well to moderately differentiated, intestinal type |
| 4 | Negative | Multifocal, weak staining in carcinoma | Moderately to poorly differentiated, intestinal type |
| 5 | Moderate to intense staining (<10% of tumor-associated stroma) | Focal, weak staining in carcinoma | Poorly differentiated, intestinal type |
| 6 | Negative | Negative | Poorly differentiated, mixed intestinal and diffuse types |
| 7 | Moderate to intense staining (<10% of tumor-associated stroma) | Negative | Poorly differentiated, diffuse type |
| 8 | Negative | Negative | Moderately to poorly differentiated, intestinal type |
| 9 | Moderate to intense staining (<10% of tumor-associated stroma) | Negative | Moderately to poorly differentiated, intestinal and diffuse types |
| 10 | Negative | Negative | Poorly differentiated, mixed intestinal and diffuse types |
