## Supplemental File 6 for "Somatic Mutations in Collagens are Associated with a Distinct Tumor Environment and Overall Survival in Gastric Cancer"

Figure S1A

ACRG
Targeted Sequencing

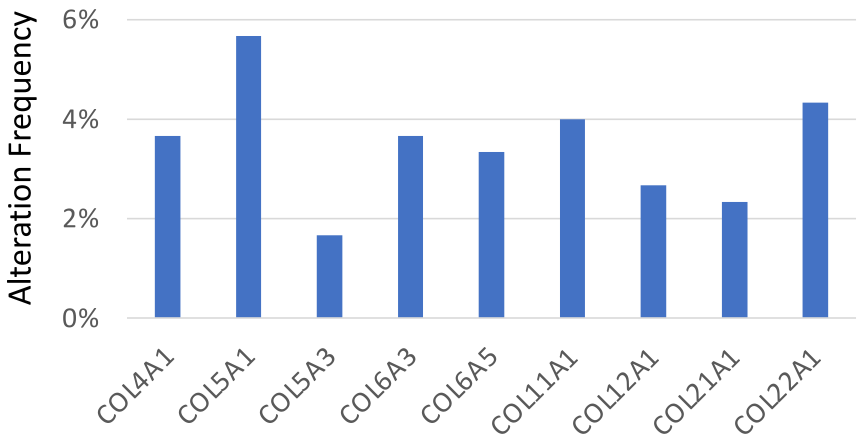


Pfizer/Hong Kong
Whole Genome
Sequencing

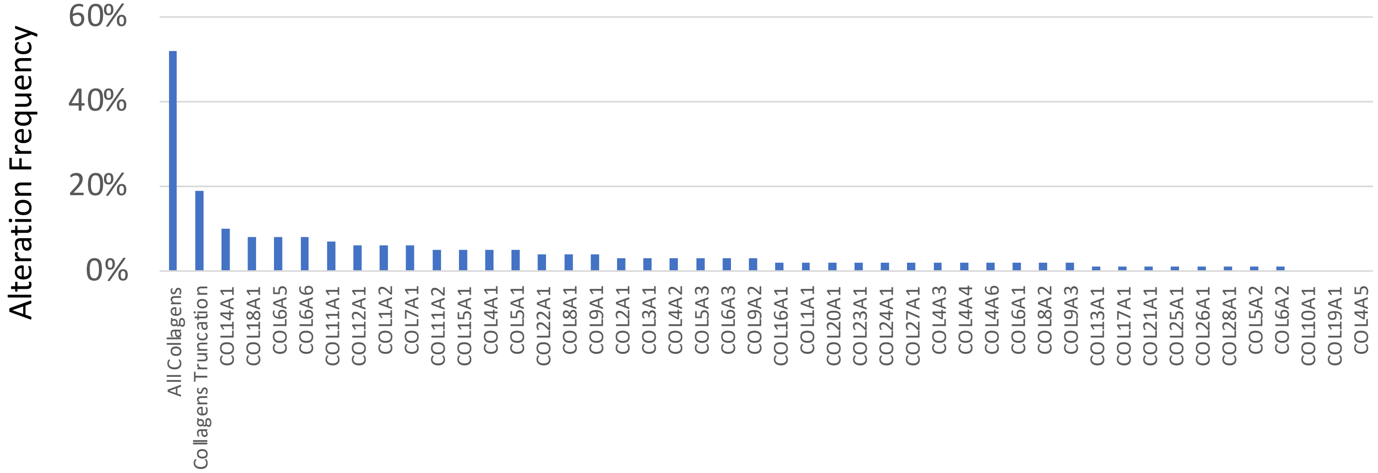

Figure S1B

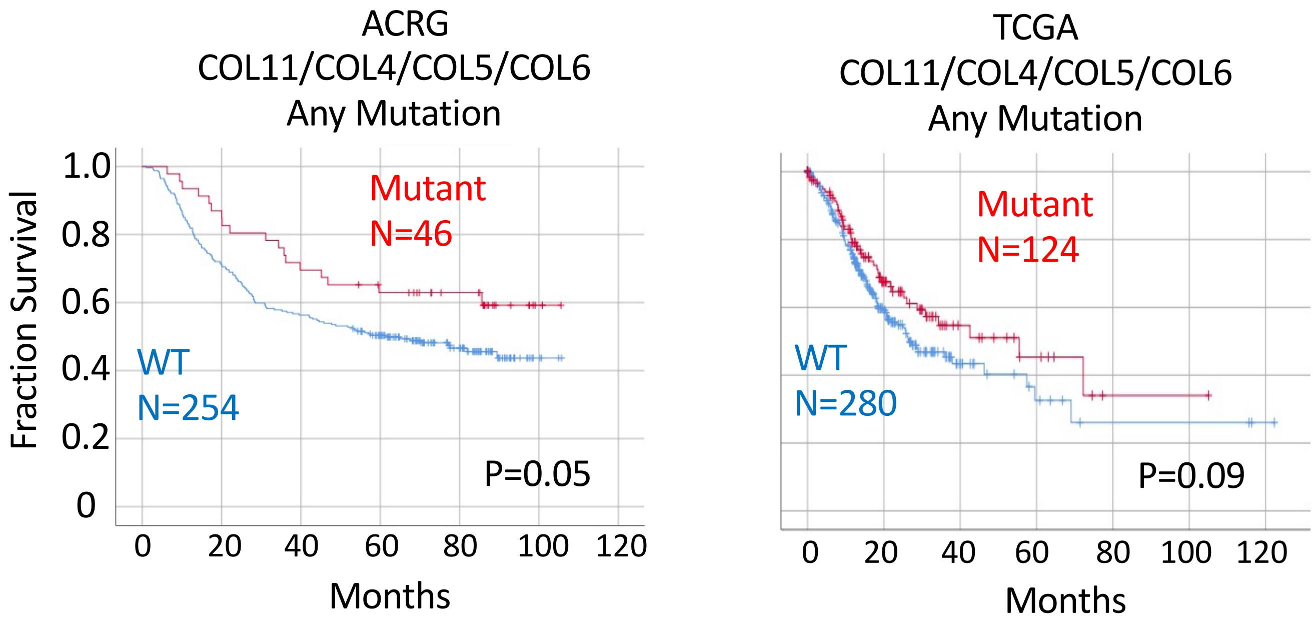
ACRG
COL11/COL4/COL5/COL6
Any Mutation
Fraction Survival
1.0
0.8
0.6
0.4
0.2
0
Mutant
N=46
WT
N=254
P=0.05
0
20
40
60
80
100
120
Months
TCGA
COL11/COL4/COL5/COL6
Any Mutation
Mutant
N=124
WT
N=280
P=0.09
0
20
40
60
80
100
120
Months

### Figure S2A

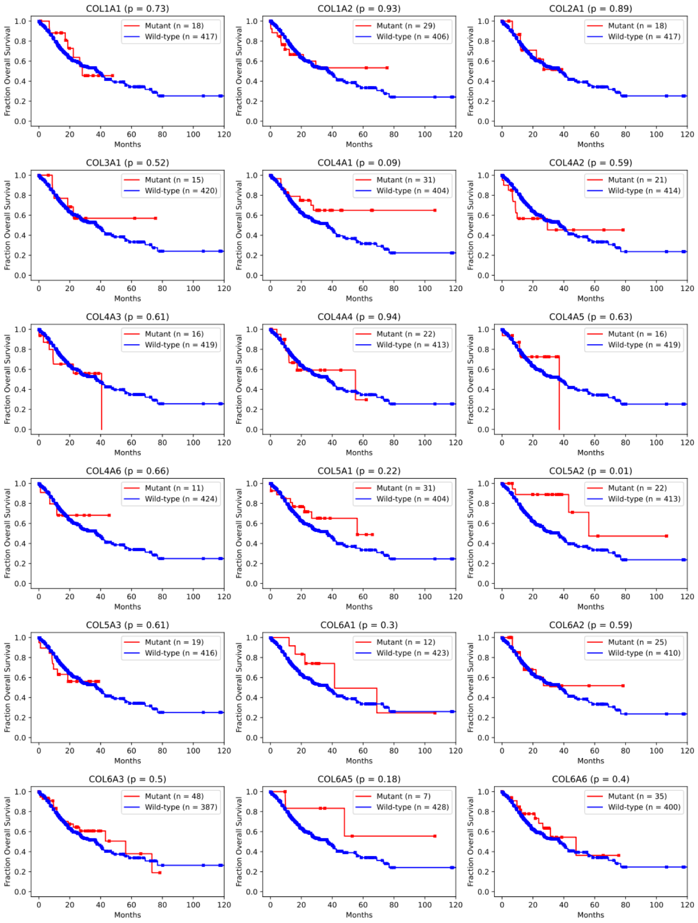
COL1A1 (p = 0.73)
Mutant (n = 18)
Wild-type (n = 417)
COL1A2 (p = 0.93)
Mutant (n = 29)
Wild-type (n = 406)
COL2A1 (p = 0.89)
Mutant (n = 18)
Wild-type (n = 417)
COL3A1 (p = 0.52)
Mutant (n = 15)
Wild-type (n = 420)
COL4A1 (p = 0.09)
Mutant (n = 31)
Wild-type (n = 404)
COL4A2 (p = 0.59)
Mutant (n = 21)
Wild-type (n = 414)
COL4A3 (p = 0.61)
Mutant (n = 16)
Wild-type (n = 419)
COL4A4 (p = 0.94)
Mutant (n = 22)
Wild-type (n = 413)
COL4A5 (p = 0.63)
Mutant (n = 16)
Wild-type (n = 419)
COL4A6 (p = 0.66)
Mutant (n = 11)
Wild-type (n = 424)
COL5A1 (p = 0.22)
Mutant (n = 31)
Wild-type (n = 404)
COL5A2 (p = 0.01)
Mutant (n = 22)
Wild-type (n = 413)
COL5A3 (p = 0.61)
Mutant (n = 19)
Wild-type (n = 416)
COL6A1 (p = 0.3)
Mutant (n = 12)
Wild-type (n = 423)
COL6A2 (p = 0.59)
Mutant (n = 25)
Wild-type (n = 410)
COL6A3 (p = 0.5)
Mutant (n = 48)
Wild-type (n = 387)
COL6A5 (p = 0.18)
Mutant (n = 7)
Wild-type (n = 428)
COL6A6 (p = 0.4)
Mutant (n = 35)
Wild-type (n = 400)
Fraction Overall Survival
Months

### Figure S2A

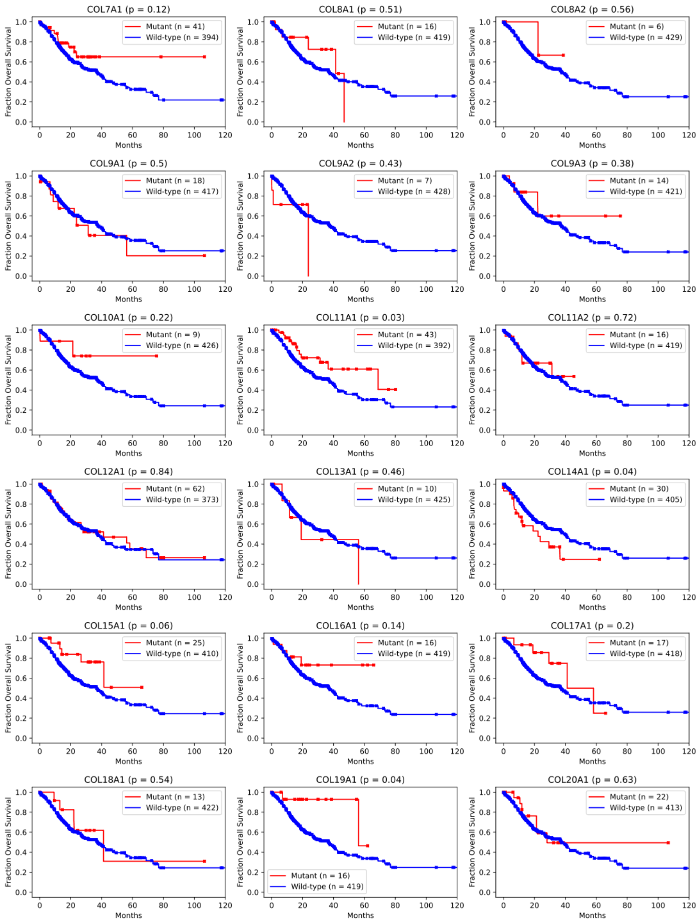
COL7A1 (p = 0.12)
Mutant (n = 41)
Wild-type (n = 394)
COL8A1 (p = 0.51)
Mutant (n = 16)
Wild-type (n = 419)
COL8A2 (p = 0.56)
Mutant (n = 6)
Wild-type (n = 429)
COL9A1 (p = 0.5)
Mutant (n = 18)
Wild-type (n = 417)
COL9A2 (p = 0.43)
Mutant (n = 7)
Wild-type (n = 428)
COL9A3 (p = 0.38)
Mutant (n = 14)
Wild-type (n = 421)
COL10A1 (p = 0.22)
Mutant (n = 9)
Wild-type (n = 426)
COL11A1 (p = 0.03)
Mutant (n = 43)
Wild-type (n = 392)
COL11A2 (p = 0.72)
Mutant (n = 16)
Wild-type (n = 419)
COL12A1 (p = 0.84)
Mutant (n = 62)
Wild-type (n = 373)
COL13A1 (p = 0.46)
Mutant (n = 10)
Wild-type (n = 425)
COL14A1 (p = 0.04)
Mutant (n = 30)
Wild-type (n = 405)
COL15A1 (p = 0.06)
Mutant (n = 25)
Wild-type (n = 410)
COL16A1 (p = 0.14)
Mutant (n = 16)
Wild-type (n = 419)
COL17A1 (p = 0.2)
Mutant (n = 17)
Wild-type (n = 418)
COL18A1 (p = 0.54)
Mutant (n = 13)
Wild-type (n = 422)
COL19A1 (p = 0.04)
Mutant (n = 16)
Wild-type (n = 419)
COL20A1 (p = 0.63)
Mutant (n = 22)
Wild-type (n = 413)
Fraction Overall Survival
Months

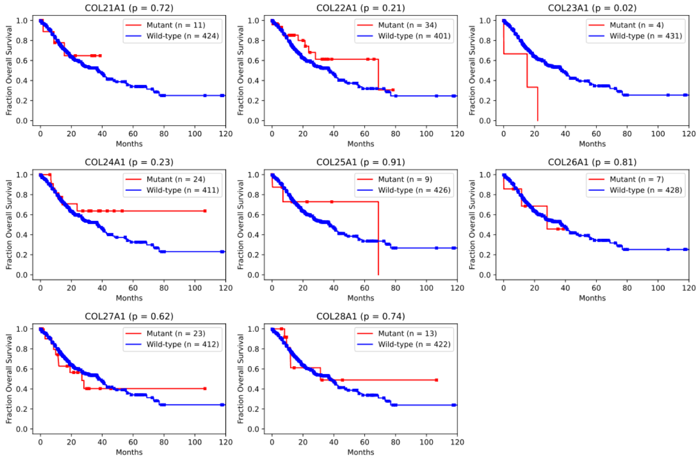
COL21A1 (p = 0.72)
Mutant (n = 11)
Wild-type (n = 424)
COL22A1 (p = 0.21)
Mutant (n = 34)
Wild-type (n = 401)
COL23A1 (p = 0.02)
Mutant (n = 4)
Wild-type (n = 431)
COL24A1 (p = 0.23)
Mutant (n = 24)
Wild-type (n = 411)
COL25A1 (p = 0.91)
Mutant (n = 9)
Wild-type (n = 426)
COL26A1 (p = 0.81)
Mutant (n = 7)
Wild-type (n = 428)
COL27A1 (p = 0.62)
Mutant (n = 23)
Wild-type (n = 412)
COL28A1 (p = 0.74)
Mutant (n = 13)
Wild-type (n = 422)
Fraction Overall Survival
Months

### Figure S2B

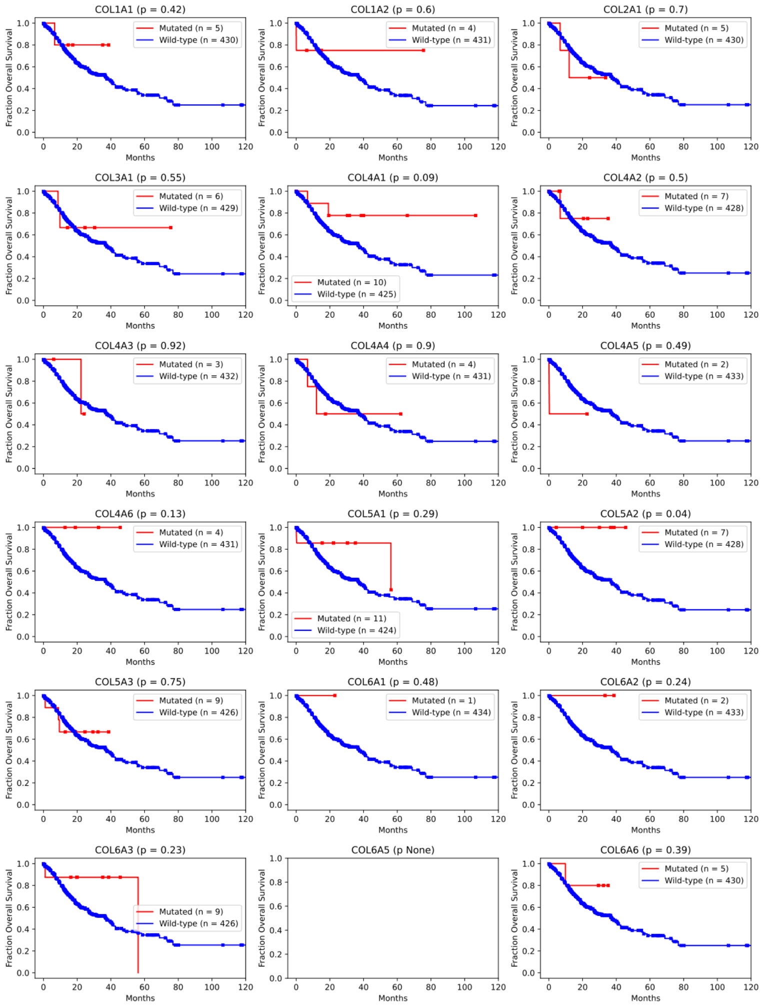

COL1A1 (p = 0.42)
Mutated (n = 5)
Wild-type (n = 430)
COL1A2 (p = 0.6)
Mutated (n = 4)
Wild-type (n = 431)
COL2A1 (p = 0.7)
Mutated (n = 5)
Wild-type (n = 430)
COL3A1 (p = 0.55)
Mutated (n = 6)
Wild-type (n = 429)
COL4A1 (p = 0.09)
Mutated (n = 10)
Wild-type (n = 425)
COL4A2 (p = 0.5)
Mutated (n = 7)
Wild-type (n = 428)
COL4A3 (p = 0.92)
Mutated (n = 3)
Wild-type (n = 432)
COL4A4 (p = 0.9)
Mutated (n = 4)
Wild-type (n = 431)
COL4A5 (p = 0.49)
Mutated (n = 2)
Wild-type (n = 433)
COL4A6 (p = 0.13)
Mutated (n = 4)
Wild-type (n = 431)
COL5A1 (p = 0.29)
Mutated (n = 11)
Wild-type (n = 424)
COL5A2 (p = 0.04)
Mutated (n = 7)
Wild-type (n = 428)
COL5A3 (p = 0.75)
Mutated (n = 9)
Wild-type (n = 426)
COL6A1 (p = 0.48)
Mutated (n = 1)
Wild-type (n = 434)
COL6A2 (p = 0.24)
Mutated (n = 2)
Wild-type (n = 433)
COL6A3 (p = 0.23)
Mutated (n = 9)
Wild-type (n = 426)
COL6A5 (p None)
COL6A6 (p = 0.39)
Mutated (n = 5)
Wild-type (n = 430)
Fraction Overall Survival
Months

### Figure S2B

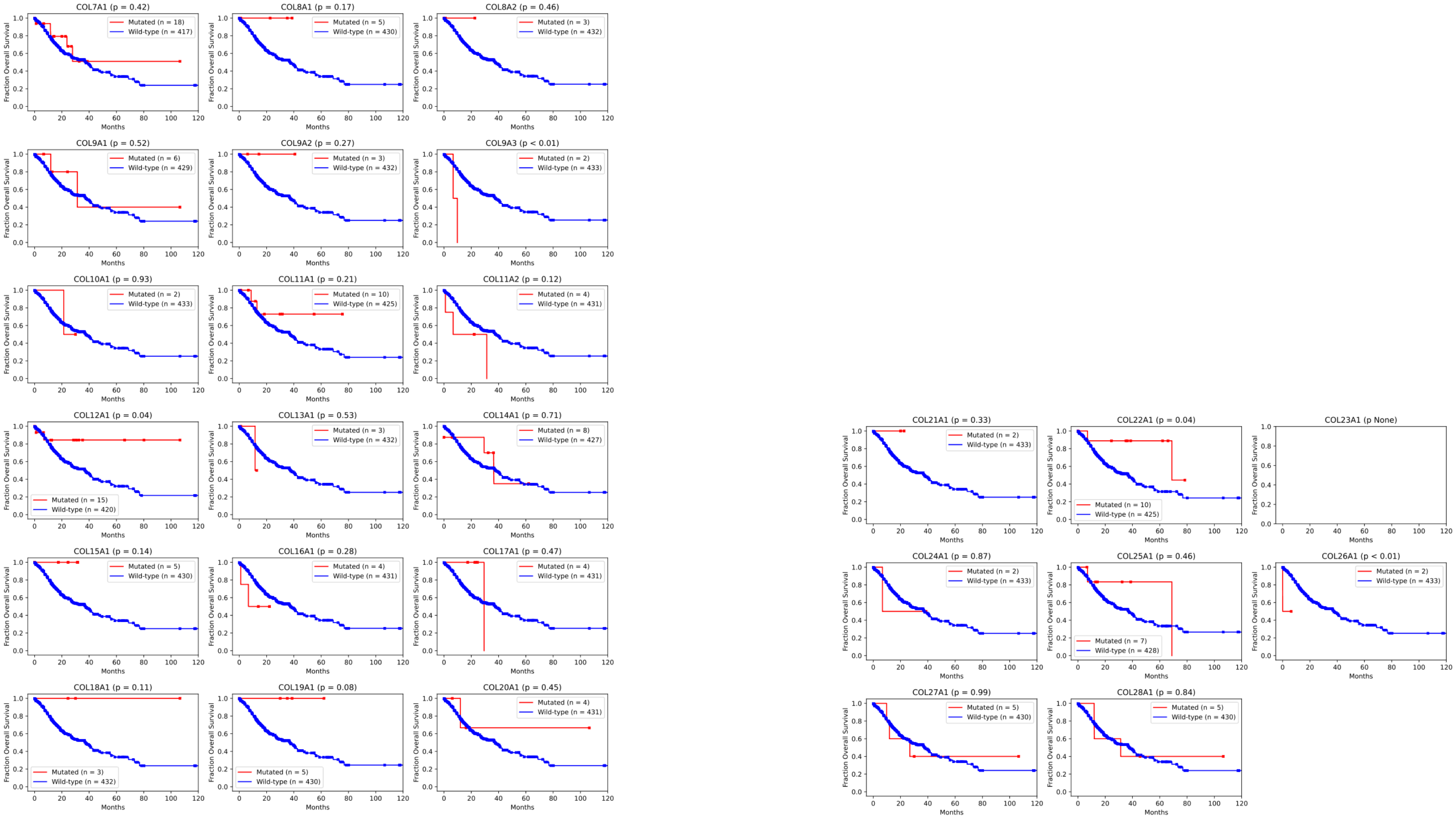

COL7A1 (p = 0.42)
Mutated (n = 18)
Wild-type (n = 417)
COL8A1 (p = 0.17)
Mutated (n = 5)
Wild-type (n = 430)
COL8A2 (p = 0.46)
Mutated (n = 3)
Wild-type (n = 432)
COL9A1 (p = 0.52)
Mutated (n = 6)
Wild-type (n = 429)
COL9A2 (p = 0.27)
Mutated (n = 3)
Wild-type (n = 432)
COL9A3 (p < 0.01)
Mutated (n = 2)
Wild-type (n = 433)
COL10A1 (p = 0.93)
Mutated (n = 2)
Wild-type (n = 433)
COL11A1 (p = 0.21)
Mutated (n = 10)
Wild-type (n = 425)
COL11A2 (p = 0.12)
Mutated (n = 4)
Wild-type (n = 431)
COL12A1 (p = 0.04)
Mutated (n = 15)
Wild-type (n = 420)
COL13A1 (p = 0.53)
Mutated (n = 3)
Wild-type (n = 432)
COL14A1 (p = 0.71)
Mutated (n = 8)
Wild-type (n = 427)
COL15A1 (p = 0.14)
Mutated (n = 5)
Wild-type (n = 430)
COL16A1 (p = 0.28)
Mutated (n = 4)
Wild-type (n = 431)
COL17A1 (p = 0.47)
Mutated (n = 4)
Wild-type (n = 431)
COL18A1 (p = 0.11)
Mutated (n = 3)
Wild-type (n = 432)
COL19A1 (p = 0.08)
Mutated (n = 5)
Wild-type (n = 430)
COL20A1 (p = 0.45)
Mutated (n = 4)
Wild-type (n = 431)
COL21A1 (p = 0.33)
Mutated (n = 2)
Wild-type (n = 433)
COL22A1 (p = 0.04)
Mutated (n = 10)
Wild-type (n = 425)
COL23A1 (p None)
COL24A1 (p = 0.87)
Mutated (n = 2)
Wild-type (n = 433)
COL25A1 (p = 0.46)
Mutated (n = 7)
Wild-type (n = 428)
COL26A1 (p < 0.01)
Mutated (n = 2)
Wild-type (n = 433)
COL27A1 (p = 0.99)
Mutated (n = 5)
Wild-type (n = 430)
COL28A1 (p = 0.84)
Mutated (n = 5)
Wild-type (n = 430)
Fraction Overall Survival
Months

### Figure S2C

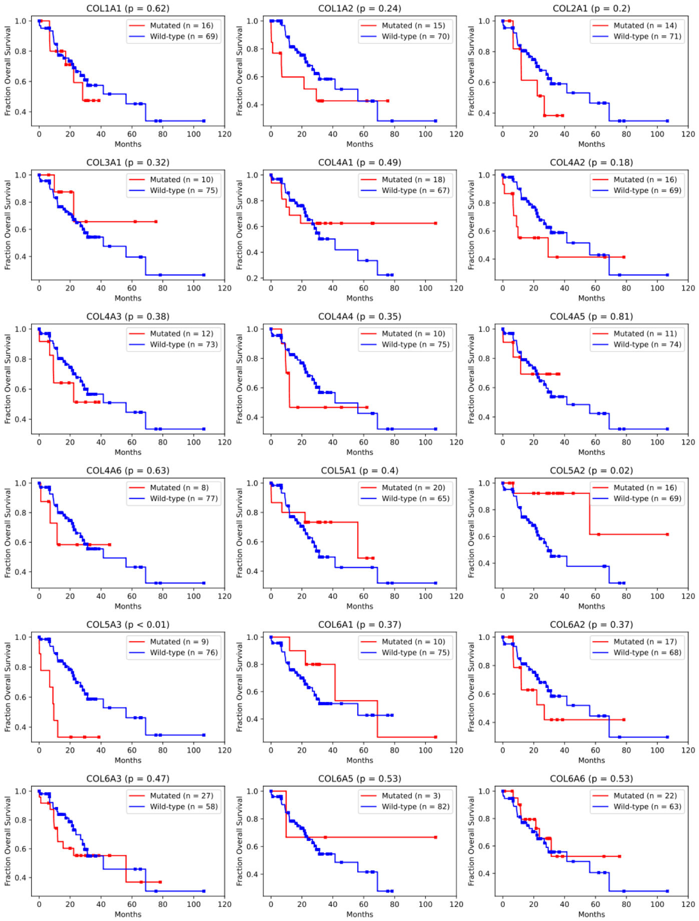

COL1A1 (p = 0.62)
Mutated (n = 16)
Wild-type (n = 69)
COL1A2 (p = 0.24)
Mutated (n = 15)
Wild-type (n = 70)
COL2A1 (p = 0.2)
Mutated (n = 14)
Wild-type (n = 71)
COL3A1 (p = 0.32)
Mutated (n = 10)
Wild-type (n = 75)
COL4A1 (p = 0.49)
Mutated (n = 18)
Wild-type (n = 67)
COL4A2 (p = 0.18)
Mutated (n = 16)
Wild-type (n = 69)
COL4A3 (p = 0.38)
Mutated (n = 12)
Wild-type (n = 73)
COL4A4 (p = 0.35)
Mutated (n = 10)
Wild-type (n = 75)
COL4A5 (p = 0.81)
Mutated (n = 11)
Wild-type (n = 74)
COL4A6 (p = 0.63)
Mutated (n = 8)
Wild-type (n = 77)
COL5A1 (p = 0.4)
Mutated (n = 20)
Wild-type (n = 65)
COL5A2 (p = 0.02)
Mutated (n = 16)
Wild-type (n = 69)
COL5A3 (p < 0.01)
Mutated (n = 9)
Wild-type (n = 76)
COL6A1 (p = 0.37)
Mutated (n = 10)
Wild-type (n = 75)
COL6A2 (p = 0.37)
Mutated (n = 17)
Wild-type (n = 68)
COL6A3 (p = 0.47)
Mutated (n = 27)
Wild-type (n = 58)
COL6A5 (p = 0.53)
Mutated (n = 3)
Wild-type (n = 82)
COL6A6 (p = 0.53)
Mutated (n = 22)
Wild-type (n = 63)
Fraction Overall Survival
Months

### Figure S2C

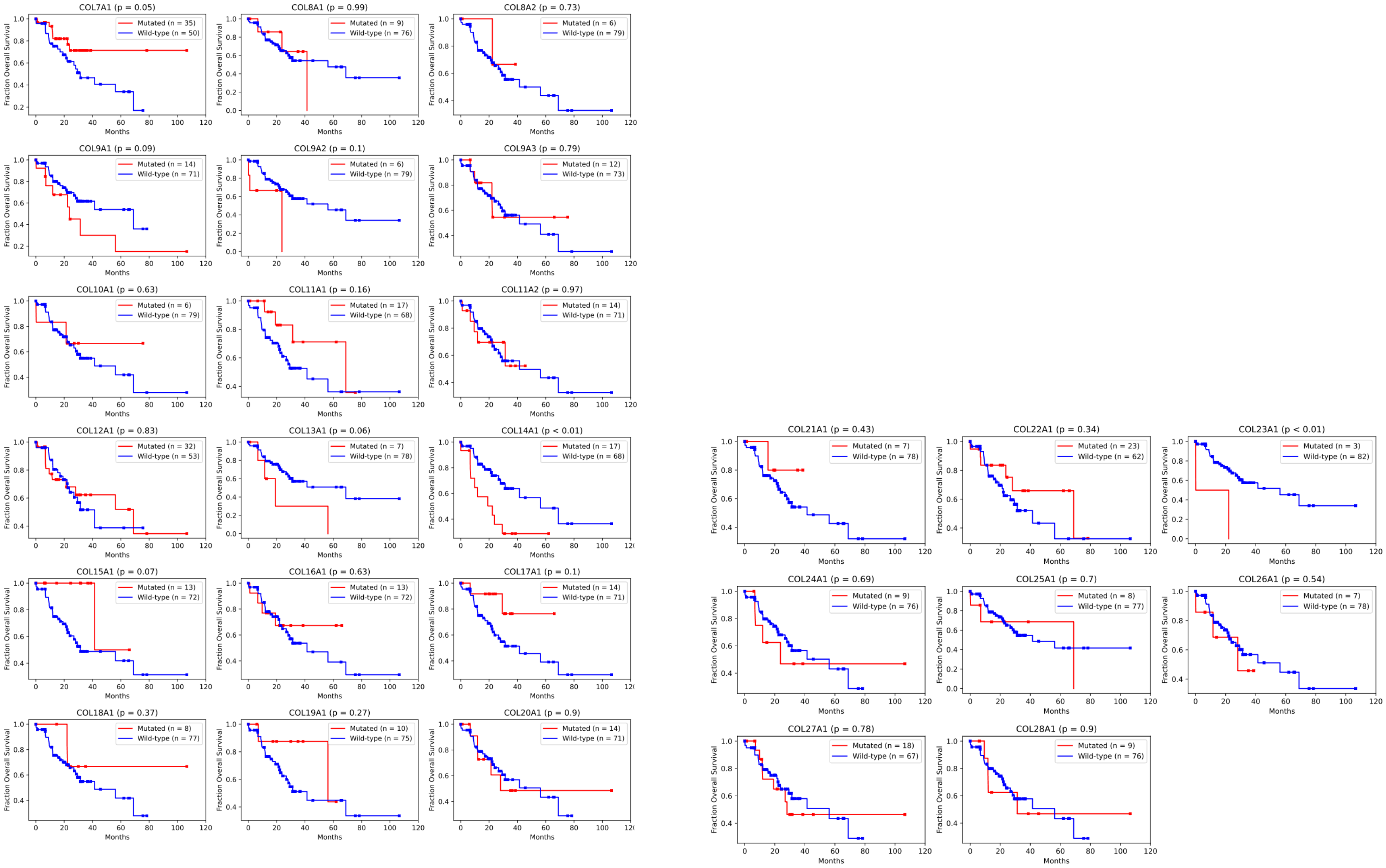
COL7A1 (p = 0.05)
Mutated (n = 35)
Wild-type (n = 50)
COL8A1 (p = 0.99)
Mutated (n = 9)
Wild-type (n = 76)
COL8A2 (p = 0.73)
Mutated (n = 6)
Wild-type (n = 79)
COL9A1 (p = 0.09)
Mutated (n = 14)
Wild-type (n = 71)
COL9A2 (p = 0.1)
Mutated (n = 6)
Wild-type (n = 79)
COL9A3 (p = 0.79)
Mutated (n = 12)
Wild-type (n = 73)
COL10A1 (p = 0.63)
Mutated (n = 6)
Wild-type (n = 79)
COL11A1 (p = 0.16)
Mutated (n = 17)
Wild-type (n = 68)
COL11A2 (p = 0.97)
Mutated (n = 14)
Wild-type (n = 71)
COL12A1 (p = 0.83)
Mutated (n = 32)
Wild-type (n = 53)
COL13A1 (p = 0.06)
Mutated (n = 7)
Wild-type (n = 78)
COL14A1 (p < 0.01)
Mutated (n = 17)
Wild-type (n = 68)
COL15A1 (p = 0.07)
Mutated (n = 13)
Wild-type (n = 72)
COL16A1 (p = 0.63)
Mutated (n = 13)
Wild-type (n = 72)
COL17A1 (p = 0.1)
Mutated (n = 14)
Wild-type (n = 71)
COL18A1 (p = 0.37)
Mutated (n = 8)
Wild-type (n = 77)
COL19A1 (p = 0.27)
Mutated (n = 10)
Wild-type (n = 75)
COL20A1 (p = 0.9)
Mutated (n = 14)
Wild-type (n = 71)
COL21A1 (p = 0.43)
Mutated (n = 7)
Wild-type (n = 78)
COL22A1 (p = 0.34)
Mutated (n = 23)
Wild-type (n = 62)
COL23A1 (p < 0.01)
Mutated (n = 3)
Wild-type (n = 82)
COL24A1 (p = 0.69)
Mutated (n = 9)
Wild-type (n = 76)
COL25A1 (p = 0.7)
Mutated (n = 8)
Wild-type (n = 77)
COL26A1 (p = 0.54)
Mutated (n = 7)
Wild-type (n = 78)
COL27A1 (p = 0.78)
Mutated (n = 18)
Wild-type (n = 67)
COL28A1 (p = 0.9)
Mutated (n = 9)
Wild-type (n = 76)
Fraction Overall Survival
Months

Figure S3A

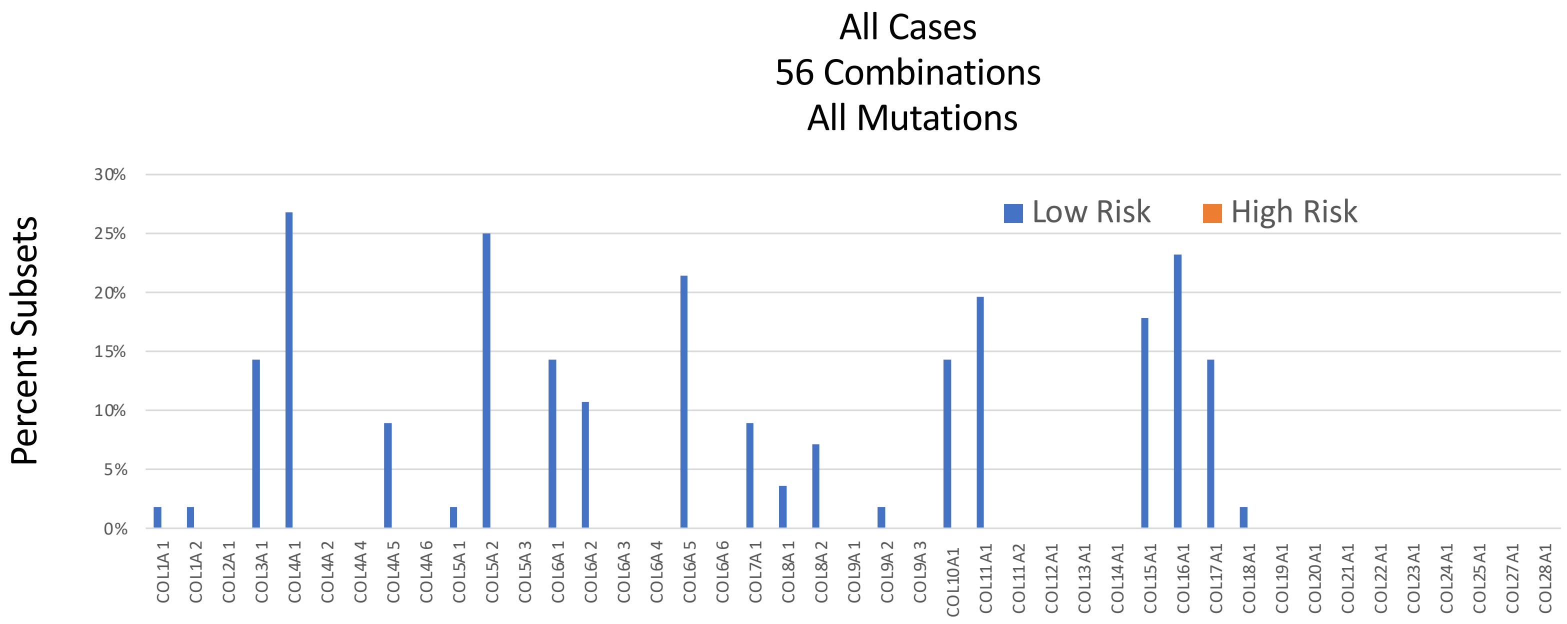
All Cases
56 Combinations
All Mutations
Percent Subsets
30%
25%
20%
15%
10%
5%
0%
Low Risk
High Risk
COL1A1
COL1A2
COL2A1
COL3A1
COL4A1
COL4A2
COL4A4
COL4A5
COL4A6
COL5A1
COL5A2
COL5A3
COL6A1
COL6A2
COL6A3
COL6A4
COL6A5
COL6A6
COL7A1
COL8A1
COL8A2
COL9A1
COL9A2
COL9A3
COL10A1
COL11A1
COL11A2
COL12A1
COL13A1
COL14A1
COL15A1
COL16A1
COL17A1
COL18A1
COL19A1
COL20A1
COL21A1
COL22A1
COL23A1
COL24A1
COL25A1
COL27A1
COL28A1

Figure S3B

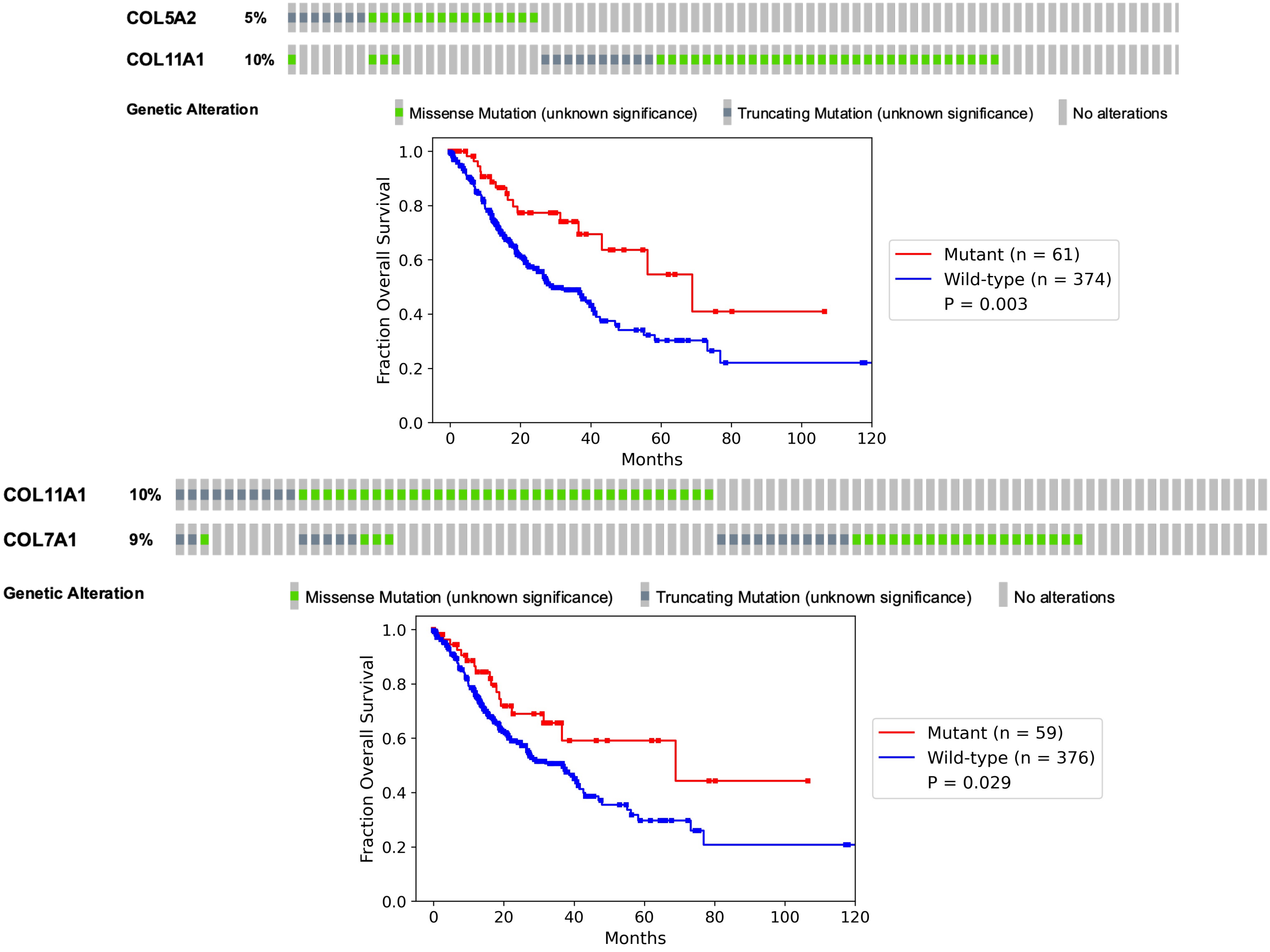

COL5A2 5%
COL11A1 10%
Genetic Alteration
Missense Mutation (unknown significance)
Truncating Mutation (unknown significance)
No alterations
Fraction Overall Survival
Months
Mutant (n = 61)
Wild-type (n = 374)
P = 0.003
COL11A1 10%
COL7A1 9%
Genetic Alteration
Missense Mutation (unknown significance)
Truncating Mutation (unknown significance)
No alterations
Fraction Overall Survival
Months
Mutant (n = 59)
Wild-type (n = 376)
P = 0.029

Figure S3C

MSS
66 Combinations

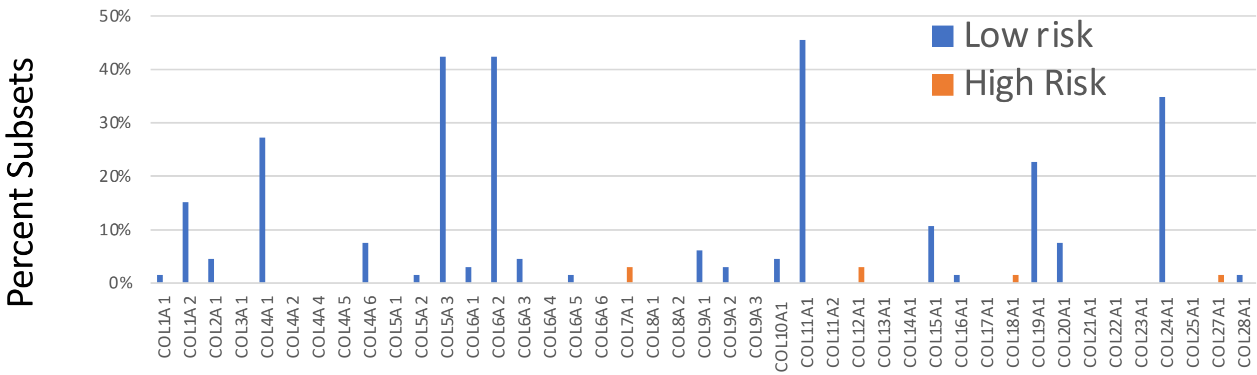


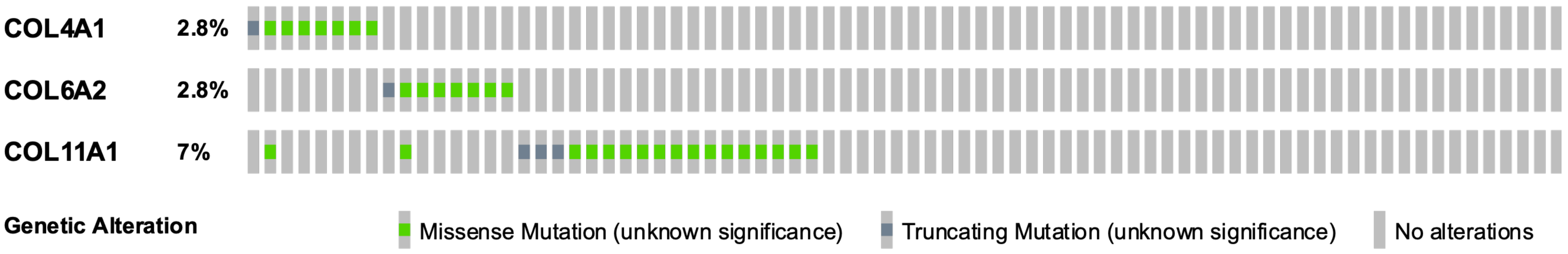


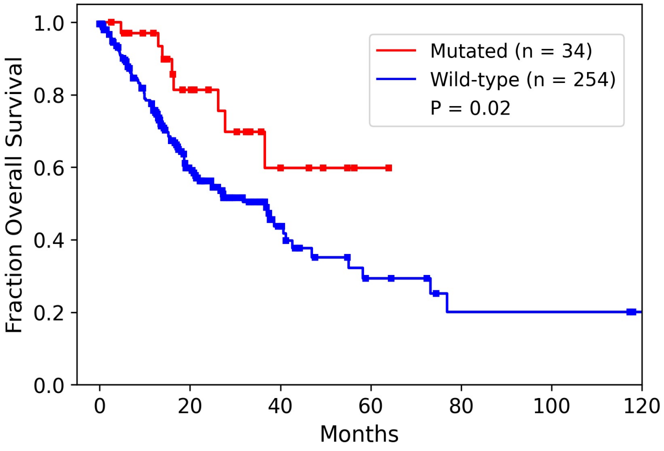

Figure S3D

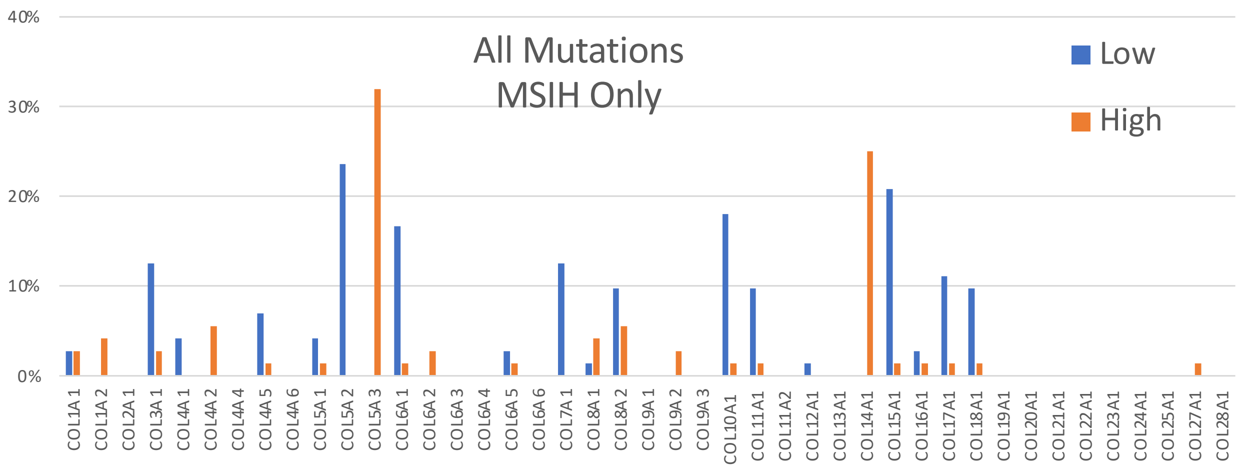

All Mutations
MSIH Only
Low
High
40%
30%
20%
10%
0%
COL1A1
COL1A2
COL2A1
COL3A1
COL4A1
COL4A2
COL4A4
COL4A5
COL4A6
COL5A1
COL5A2
COL5A3
COL6A1
COL6A2
COL6A3
COL6A4
COL6A5
COL6A6
COL7A1
COL8A1
COL8A2
COL9A1
COL9A2
COL9A3
COL10A1
COL11A1
COL11A2
COL12A1
COL13A1
COL14A1
COL15A1
COL16A1
COL17A1
COL18A1
COL19A1
COL20A1
COL21A1
COL22A1
COL23A1
COL24A1
COL25A1
COL27A1
COL28A1

Figure S3E

MSIH only; most frequent significant OS associated collagens

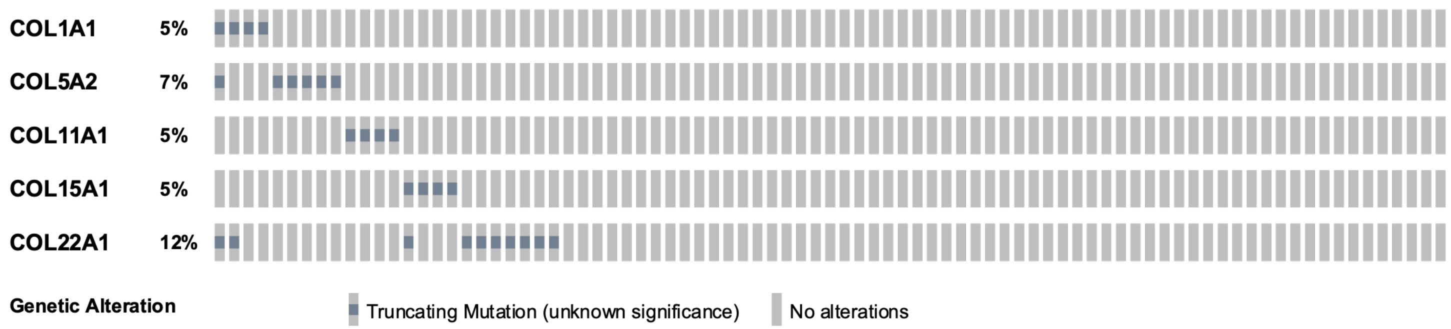


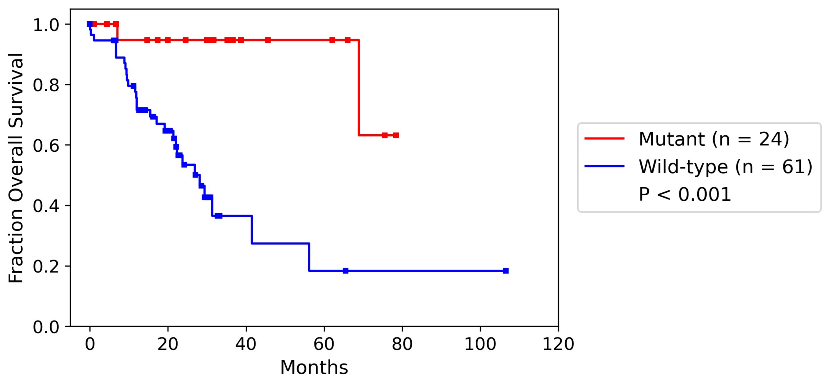

Figure S4

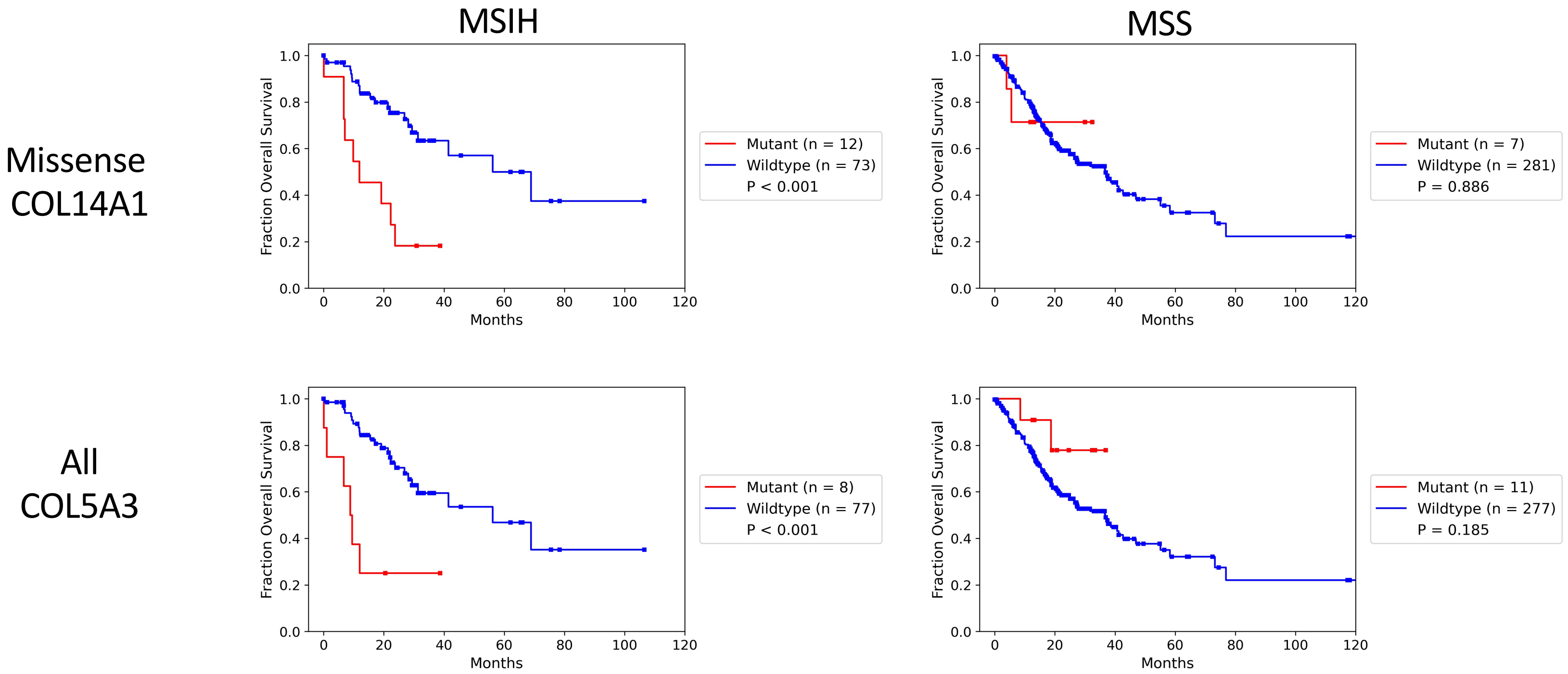
MSIH
MSS
Missense
COL14A1
All
COL5A3
Fraction Overall Survival
Months
Mutant (n = 12)
Wildtype (n = 73)
P < 0.001
Mutant (n = 7)
Wildtype (n = 281)
P = 0.886
Mutant (n = 8)
Wildtype (n = 77)
P < 0.001
Mutant (n = 11)
Wildtype (n = 277)
P = 0.185

Figure S5A

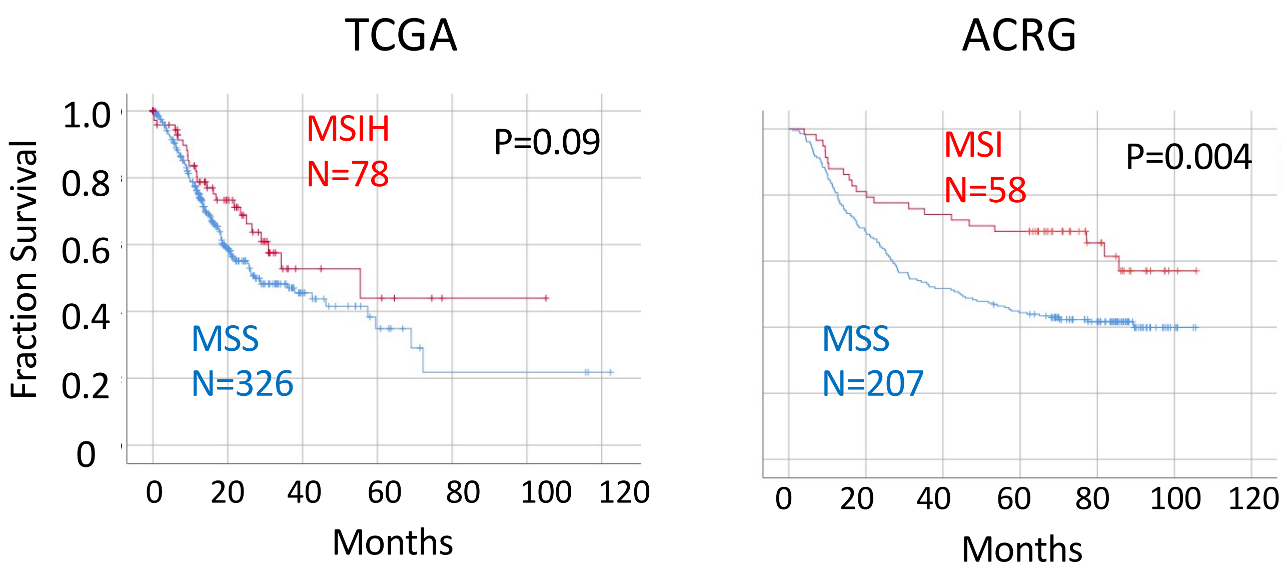

TCGA
ACRG
Fraction Survival
1.0
0.8
0.6
0.4
0.2
0
MSIH
N=78
P=0.09
MSS
N=326
MSI
N=58
P=0.004
MSS
N=207
0
20
40
60
80
100
120
Months
Months

Figure S5B

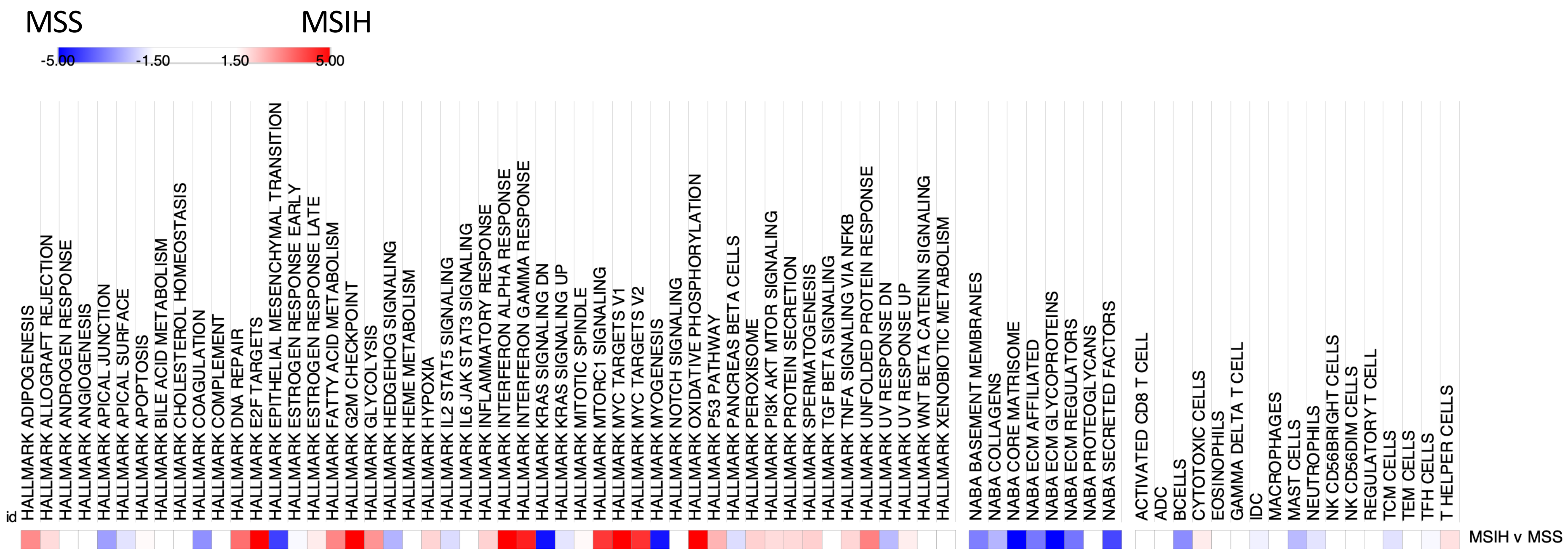
MSS
MSIH
-5.00
-1.50
1.50
5.00
id
HALLMARK ADIPOGENESIS
HALLMARK ALLOGRAFT REJECTION
HALLMARK ANDROGEN RESPONSE
HALLMARK ANGIOGENESIS
HALLMARK APICAL JUNCTION
HALLMARK APICAL SURFACE
HALLMARK APOPTOSIS
HALLMARK BILE ACID METABOLISM
HALLMARK CHOLESTEROL HOMEOSTASIS
HALLMARK COAGULATION
HALLMARK COMPLEMENT
HALLMARK DNA REPAIR
HALLMARK E2F TARGETS
HALLMARK EPITHELIAL MESENCHYMAL TRANSITION
HALLMARK ESTROGEN RESPONSE EARLY
HALLMARK ESTROGEN RESPONSE LATE
HALLMARK FATTY ACID METABOLISM
HALLMARK G2M CHECKPOINT
HALLMARK GLYCOLYSIS
HALLMARK HEDGEHOG SIGNALING
HALLMARK HEME METABOLISM
HALLMARK HYPOXIA
HALLMARK IL2 STAT5 SIGNALING
HALLMARK IL6 JAK STAT3 SIGNALING
HALLMARK INFLAMMATORY RESPONSE
HALLMARK INTERFERON ALPHA RESPONSE
HALLMARK INTERFERON GAMMA RESPONSE
HALLMARK KRAS SIGNALING DN
HALLMARK KRAS SIGNALING UP
HALLMARK MITOTIC SPINDLE
HALLMARK MTORC1 SIGNALING
HALLMARK MYC TARGETS V1
HALLMARK MYC TARGETS V2
HALLMARK MYOGENESIS
HALLMARK NOTCH SIGNALING
HALLMARK OXIDATIVE PHOSPHORYLATION
HALLMARK P53 PATHWAY
HALLMARK PANCREAS BETA CELLS
HALLMARK PEROXISOME
HALLMARK PI3K AKT MTOR SIGNALING
HALLMARK PROTEIN SECRETION
HALLMARK SPERMATOGENESIS
HALLMARK TGF BETA SIGNALING
HALLMARK TNFA SIGNALING VIA NFKB
HALLMARK UNFOLDED PROTEIN RESPONSE
HALLMARK UV RESPONSE DN
HALLMARK UV RESPONSE UP
HALLMARK WNT BETA CATENIN SIGNALING
HALLMARK XENOBIOTIC METABOLISM
NABA BASEMENT MEMBRANES
NABA COLLAGENS
NABA CORE MATRISOME
NABA ECM AFFILIATED
NABA ECM GLYCOPROTEINS
NABA ECM REGULATORS
NABA PROTEOGLYCANS
NABA SECRETED FACTORS
ACTIVATED CD8 T CELL
ADC
BCELLS
CYTOTOXIC CELLS
EOSINOPHILS
GAMMA DELTA T CELL
IDC
MACROPHAGES
MAST CELLS
NEUTROPHILS
NK CD56BRIGHT CELLS
NK CD56DIM CELLS
REGULATORY T CELL
TCM CELLS
TEM CELLS
TFH CELLS
T HELPER CELLS
MSIH v MSS

Figure S6A

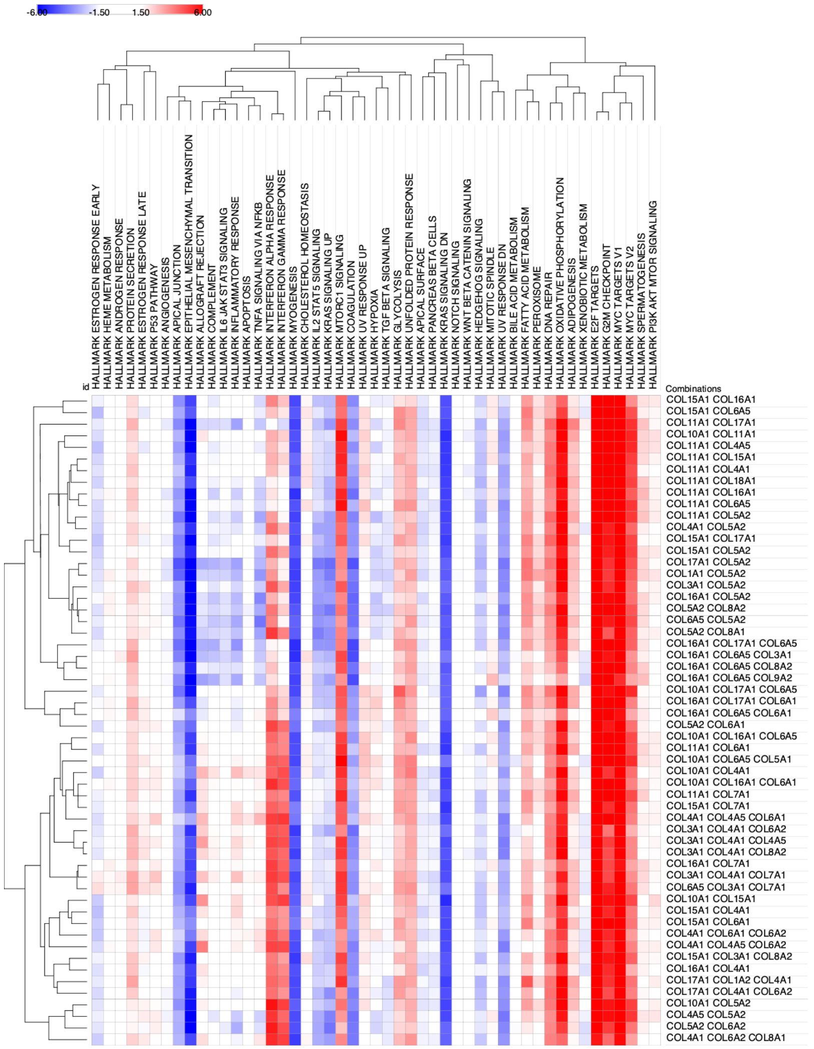

-6.00
-1.50
1.50
6.00
id
HALLMARK ESTROGEN RESPONSE EARLY
HALLMARK HEME METABOLISM
HALLMARK ANDROGEN RESPONSE
HALLMARK PROTEIN SECRETION
HALLMARK ESTROGEN RESPONSE LATE
HALLMARK P53 PATHWAY
HALLMARK ANGIOGENESIS
HALLMARK APICAL JUNCTION
HALLMARK EPITHELIAL MESENCHYMAL TRANSITION
HALLMARK ALLOGRAFT REJECTION
HALLMARK COMPLEMENT
HALLMARK IL6 JAK STAT3 SIGNALING
HALLMARK INFLAMMATORY RESPONSE
HALLMARK APOPTOSIS
HALLMARK TNFA SIGNALING VIA NFKB
HALLMARK INTERFERON ALPHA RESPONSE
HALLMARK INTERFERON GAMMA RESPONSE
HALLMARK MYOGENESIS
HALLMARK CHOLESTEROL HOMEOSTASIS
HALLMARK IL2 STAT5 SIGNALING
HALLMARK KRAS SIGNALING UP
HALLMARK MTORC1 SIGNALING
HALLMARK COAGULATION
HALLMARK UV RESPONSE UP
HALLMARK HYPOXIA
HALLMARK TGF BETA SIGNALING
HALLMARK GLYCOLYSIS
HALLMARK UNFOLDED PROTEIN RESPONSE
HALLMARK APICAL SURFACE
HALLMARK PANCREAS BETA CELLS
HALLMARK KRAS SIGNALING DN
HALLMARK NOTCH SIGNALING
HALLMARK WNT BETA CATENIN SIGNALING
HALLMARK HEDGEHOG SIGNALING
HALLMARK MITOTIC SPINDLE
HALLMARK UV RESPONSE DN
HALLMARK BILE ACID METABOLISM
HALLMARK FATTY ACID METABOLISM
HALLMARK PEROXISOME
HALLMARK DNA REPAIR
HALLMARK OXIDATIVE PHOSPHORYLATION
HALLMARK ADIPOGENESIS
HALLMARK XENOBIOTIC METABOLISM
HALLMARK E2F TARGETS
HALLMARK G2M CHECKPOINT
HALLMARK MYC TARGETS V1
HALLMARK MYC TARGETS V2
HALLMARK SPERMATOGENESIS
HALLMARK PI3K AKT MTOR SIGNALING
Combinations
COL15A1 COL16A1
COL15A1 COL6A5
COL11A1 COL17A1
COL10A1 COL11A1
COL11A1 COL4A5
COL11A1 COL15A1
COL11A1 COL4A1
COL11A1 COL18A1
COL11A1 COL16A1
COL11A1 COL6A5
COL11A1 COL5A2
COL4A1 COL5A2
COL15A1 COL17A1
COL15A1 COL5A2
COL17A1 COL5A2
COL1A1 COL5A2
COL3A1 COL5A2
COL16A1 COL5A2
COL5A2 COL8A2
COL6A5 COL5A2
COL5A2 COL8A1
COL16A1 COL17A1 COL6A5
COL16A1 COL6A5 COL3A1
COL16A1 COL6A5 COL8A2
COL16A1 COL6A5 COL9A2
COL10A1 COL17A1 COL6A5
COL16A1 COL17A1 COL6A1
COL16A1 COL6A5 COL6A1
COL5A2 COL6A1
COL10A1 COL16A1 COL6A5
COL11A1 COL6A1
COL10A1 COL6A5 COL5A1
COL10A1 COL4A1
COL10A1 COL16A1 COL6A1
COL11A1 COL7A1
COL15A1 COL7A1
COL4A1 COL4A5 COL6A1
COL3A1 COL4A1 COL6A2
COL3A1 COL4A1 COL4A5
COL3A1 COL4A1 COL8A2
COL16A1 COL7A1
COL3A1 COL4A1 COL7A1
COL6A5 COL3A1 COL7A1
COL10A1 COL15A1
COL15A1 COL4A1
COL15A1 COL6A1
COL4A1 COL6A1 COL6A2
COL4A1 COL4A5 COL6A2
COL15A1 COL3A1 COL8A2
COL16A1 COL4A1
COL17A1 COL1A2 COL4A1
COL17A1 COL4A1 COL6A2
COL10A1 COL5A2
COL4A5 COL5A2
COL5A2 COL6A2
COL4A1 COL6A2 COL8A1

### Figure S6B

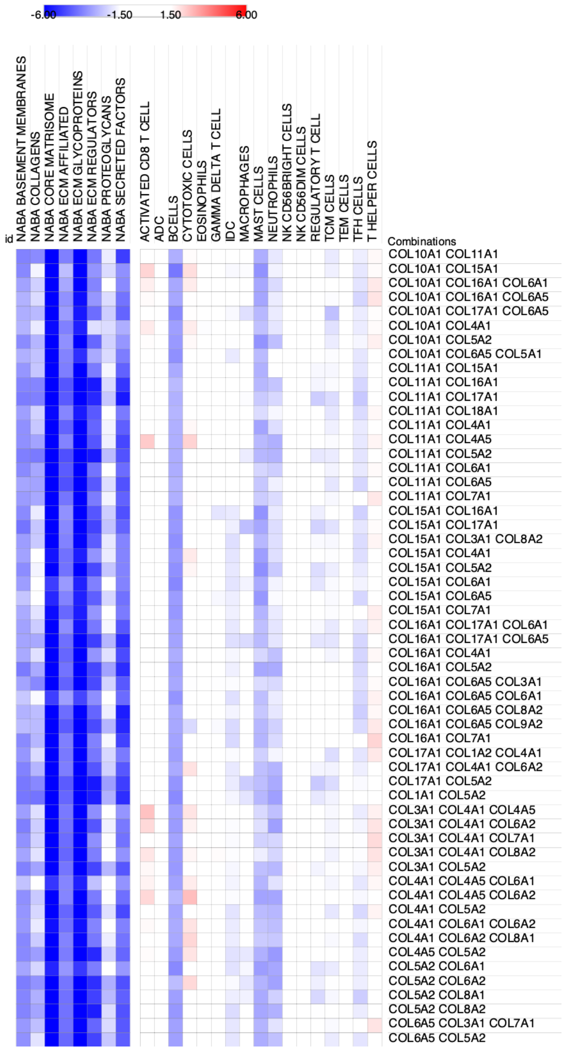

-6.00
-1.50
1.50
6.00
id
NABA BASEMENT MEMBRANES
NABA COLLAGENS
NABA CORE MATRISOME
NABA ECM AFFILIATED
NABA ECM GLYCOPROTEINS
NABA ECM REGULATORS
NABA PROTEOGLYCANS
NABA SECRETED FACTORS
ACTIVATED CD8 T CELL
ADC
BCELLS
CYTOTOXIC CELLS
EOSINOPHILS
GAMMA DELTA T CELL
IDC
MACROPHAGES
MAST CELLS
NEUTROPHILS
NK CD56BRIGHT CELLS
NK CD56DIM CELLS
REGULATORY T CELL
TCM CELLS
TEM CELLS
TFH CELLS
T HELPER CELLS
Combinations
COL10A1 COL11A1
COL10A1 COL15A1
COL10A1 COL16A1 COL6A1
COL10A1 COL16A1 COL6A5
COL10A1 COL17A1 COL6A5
COL10A1 COL4A1
COL10A1 COL5A2
COL10A1 COL6A5 COL5A1
COL11A1 COL15A1
COL11A1 COL16A1
COL11A1 COL17A1
COL11A1 COL18A1
COL11A1 COL4A1
COL11A1 COL4A5
COL11A1 COL5A2
COL11A1 COL6A1
COL11A1 COL6A5
COL11A1 COL7A1
COL15A1 COL16A1
COL15A1 COL17A1
COL15A1 COL3A1 COL8A2
COL15A1 COL4A1
COL15A1 COL5A2
COL15A1 COL6A1
COL15A1 COL6A5
COL15A1 COL7A1
COL16A1 COL17A1 COL6A1
COL16A1 COL17A1 COL6A5
COL16A1 COL4A1
COL16A1 COL5A2
COL16A1 COL6A5 COL3A1
COL16A1 COL6A5 COL6A1
COL16A1 COL6A5 COL8A2
COL16A1 COL6A5 COL9A2
COL16A1 COL7A1
COL17A1 COL1A2 COL4A1
COL17A1 COL4A1 COL6A2
COL17A1 COL5A2
COL1A1 COL5A2
COL3A1 COL4A1 COL4A5
COL3A1 COL4A1 COL6A2
COL3A1 COL4A1 COL7A1
COL3A1 COL4A1 COL8A2
COL3A1 COL5A2
COL4A1 COL4A5 COL6A1
COL4A1 COL4A5 COL6A2
COL4A1 COL5A2
COL4A1 COL6A1 COL6A2
COL4A1 COL6A2 COL8A1
COL4A5 COL5A2
COL5A2 COL6A1
COL5A2 COL6A2
COL5A2 COL8A1
COL5A2 COL8A2
COL6A5 COL3A1 COL7A1
COL6A5 COL5A2

Figure S6C

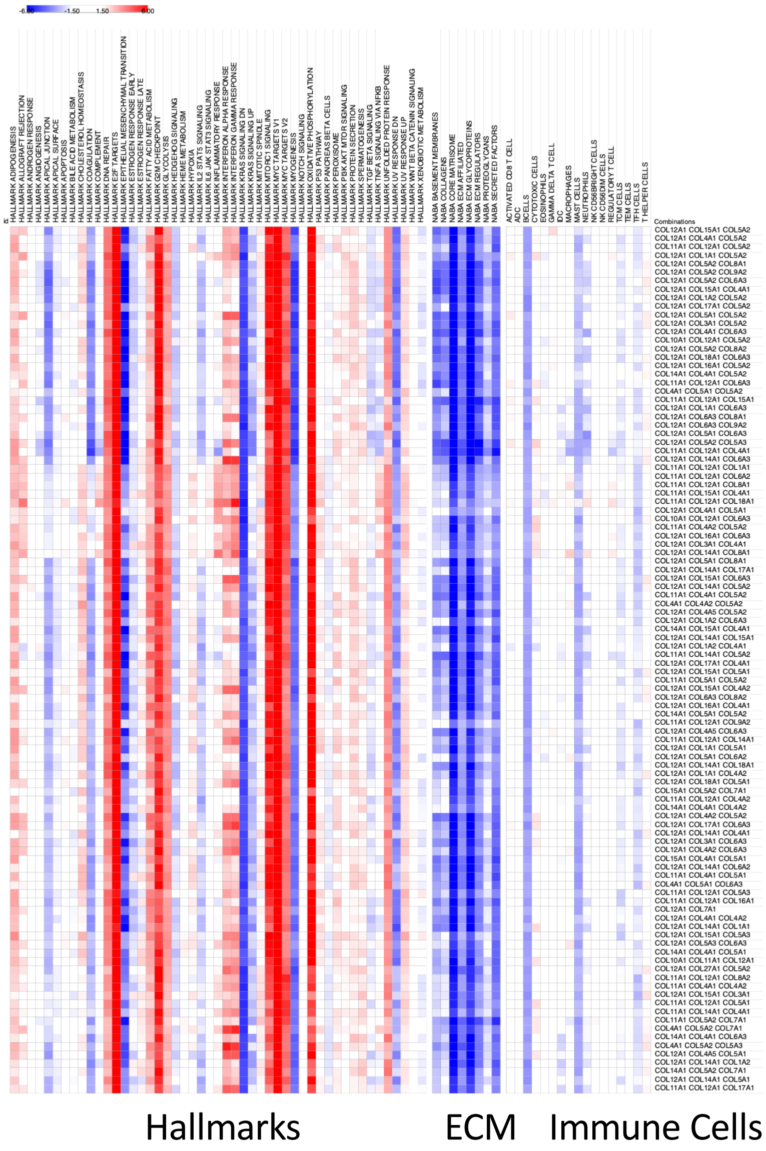

Figure S6D

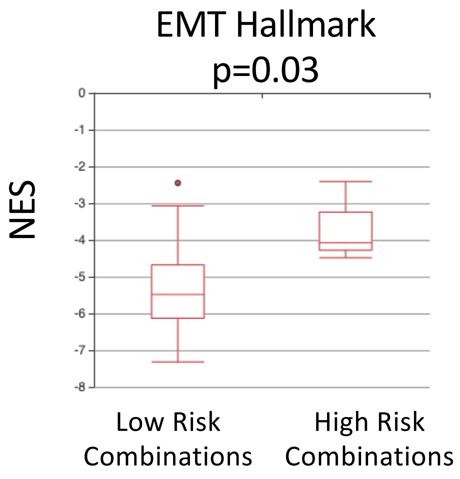


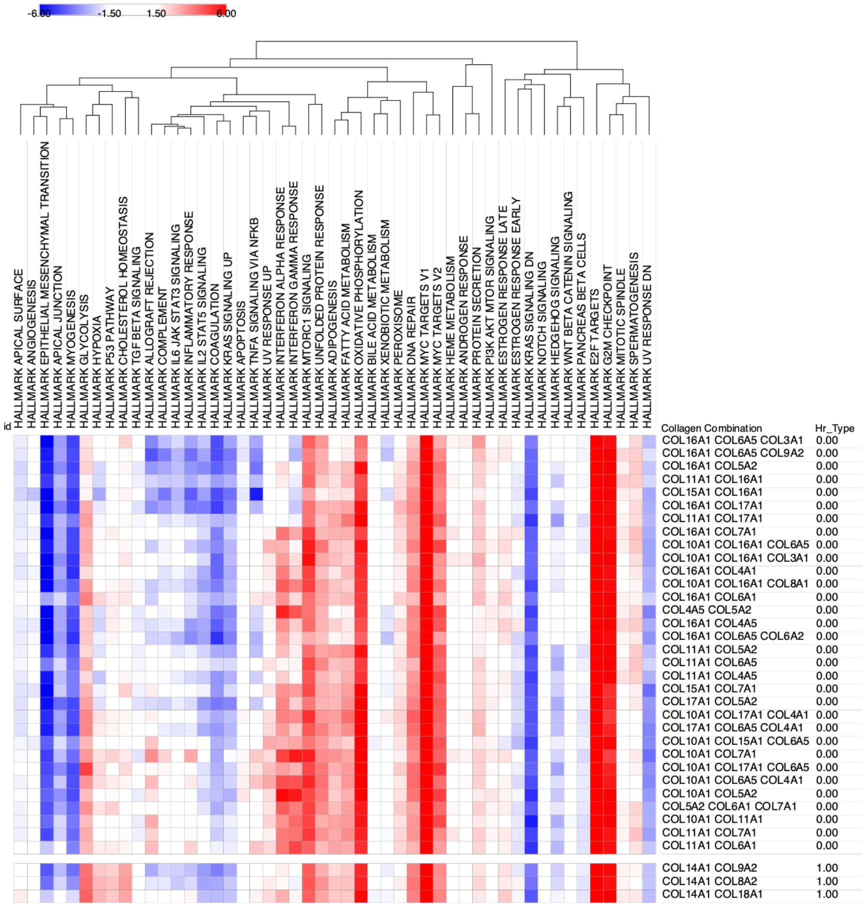


Low Risk
HR < 1

High Risk
HR > 1

Figure S7A

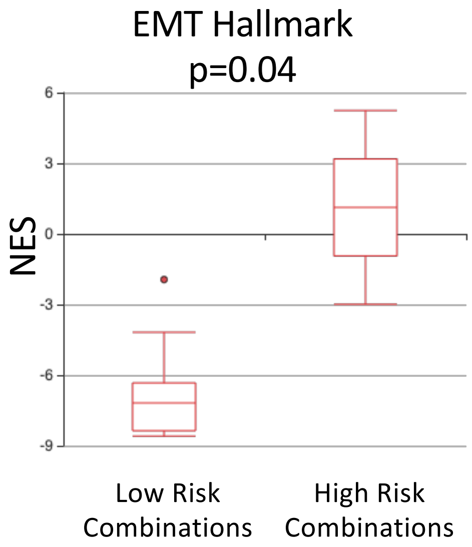
EMT Hallmark
p=0.04
NES
6
3
0
-3
-6
-9
Low Risk Combinations
High Risk Combinations

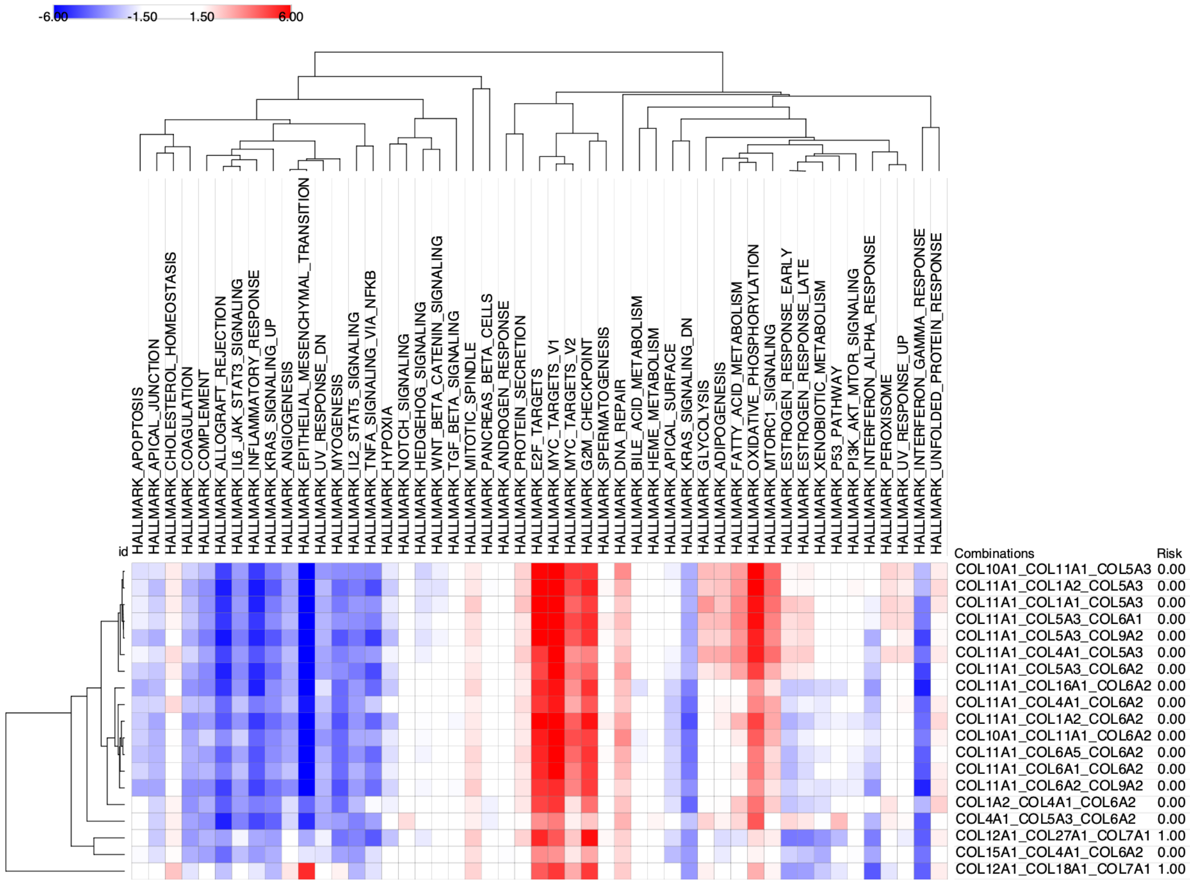
-6.00
-1.50
1.50
6.00
HALLMARK_APOPTOSIS
HALLMARK_APICAL_JUNCTION
HALLMARK_CHOLESTEROL_HOMEOSTASIS
HALLMARK_COAGULATION
HALLMARK_COMPLEMENT
HALLMARK_ALLOGRAFT_REJECTION
HALLMARK_IL6_JAK_STAT3_SIGNALING
HALLMARK_INFLAMMATORY_RESPONSE
HALLMARK_KRAS_SIGNALING_UP
HALLMARK_ANGIOGENESIS
HALLMARK_EPITHELIAL_MESENCHYMAL_TRANSITION
HALLMARK_UV_RESPONSE_DN
HALLMARK_MYOGENESIS
HALLMARK_IL2_STAT5_SIGNALING
HALLMARK_TNFA_SIGNALING_VIA_NFKB
HALLMARK_HYPOXIA
HALLMARK_NOTCH_SIGNALING
HALLMARK_HEDGEHOG_SIGNALING
HALLMARK_WNT_BETA_CATENIN_SIGNALING
HALLMARK_TGF_BETA_SIGNALING
HALLMARK_MITOTIC_SPINDLE
HALLMARK_PANCREAS_BETA_CELLS
HALLMARK_ANDROGEN_RESPONSE
HALLMARK_PROTEIN_SECRETION
HALLMARK_E2F_TARGETS
HALLMARK_MYC_TARGETS_V1
HALLMARK_MYC_TARGETS_V2
HALLMARK_G2M_CHECKPOINT
HALLMARK_SPERMATOGENESIS
HALLMARK_DNA_REPAIR
HALLMARK_BILE_ACID_METABOLISM
HALLMARK_HEME_METABOLISM
HALLMARK_APICAL_SURFACE
HALLMARK_KRAS_SIGNALING_DN
HALLMARK_GLYCOLYSIS
HALLMARK_ADIPOGENESIS
HALLMARK_FATTY_ACID_METABOLISM
HALLMARK_OXIDATIVE_PHOSPHORYLATION
HALLMARK_MTORC1_SIGNALING
HALLMARK_ESTROGEN_RESPONSE_EARLY
HALLMARK_ESTROGEN_RESPONSE_LATE
HALLMARK_XENOBIOTIC_METABOLISM
HALLMARK_P53_PATHWAY
HALLMARK_PI3K_AKT_MTOR_SIGNALING
HALLMARK_INTERFERON_ALPHA_RESPONSE
HALLMARK_PEROXISOME
HALLMARK_UV_RESPONSE_UP
HALLMARK_INTERFERON_GAMMA_RESPONSE
HALLMARK_UNFOLDED_PROTEIN_RESPONSE
id
Combinations
Risk
COL10A1_COL11A1_COL5A3 0.00
COL11A1_COL1A2_COL5A3 0.00
COL11A1_COL1A1_COL5A3 0.00
COL11A1_COL5A3_COL6A1 0.00
COL11A1_COL5A3_COL9A2 0.00
COL11A1_COL4A1_COL5A3 0.00
COL11A1_COL5A3_COL6A2 0.00
COL11A1_COL16A1_COL6A2 0.00
COL11A1_COL4A1_COL6A2 0.00
COL11A1_COL1A2_COL6A2 0.00
COL10A1_COL11A1_COL6A2 0.00
COL11A1_COL6A5_COL6A2 0.00
COL11A1_COL6A1_COL6A2 0.00
COL11A1_COL6A2_COL9A2 0.00
COL1A2_COL4A1_COL6A2 0.00
COL4A1_COL5A3_COL6A2 0.00
COL12A1_COL27A1_COL7A1 1.00
COL15A1_COL4A1_COL6A2 0.00
COL12A1_COL18A1_COL7A1 1.00

Figure S7B

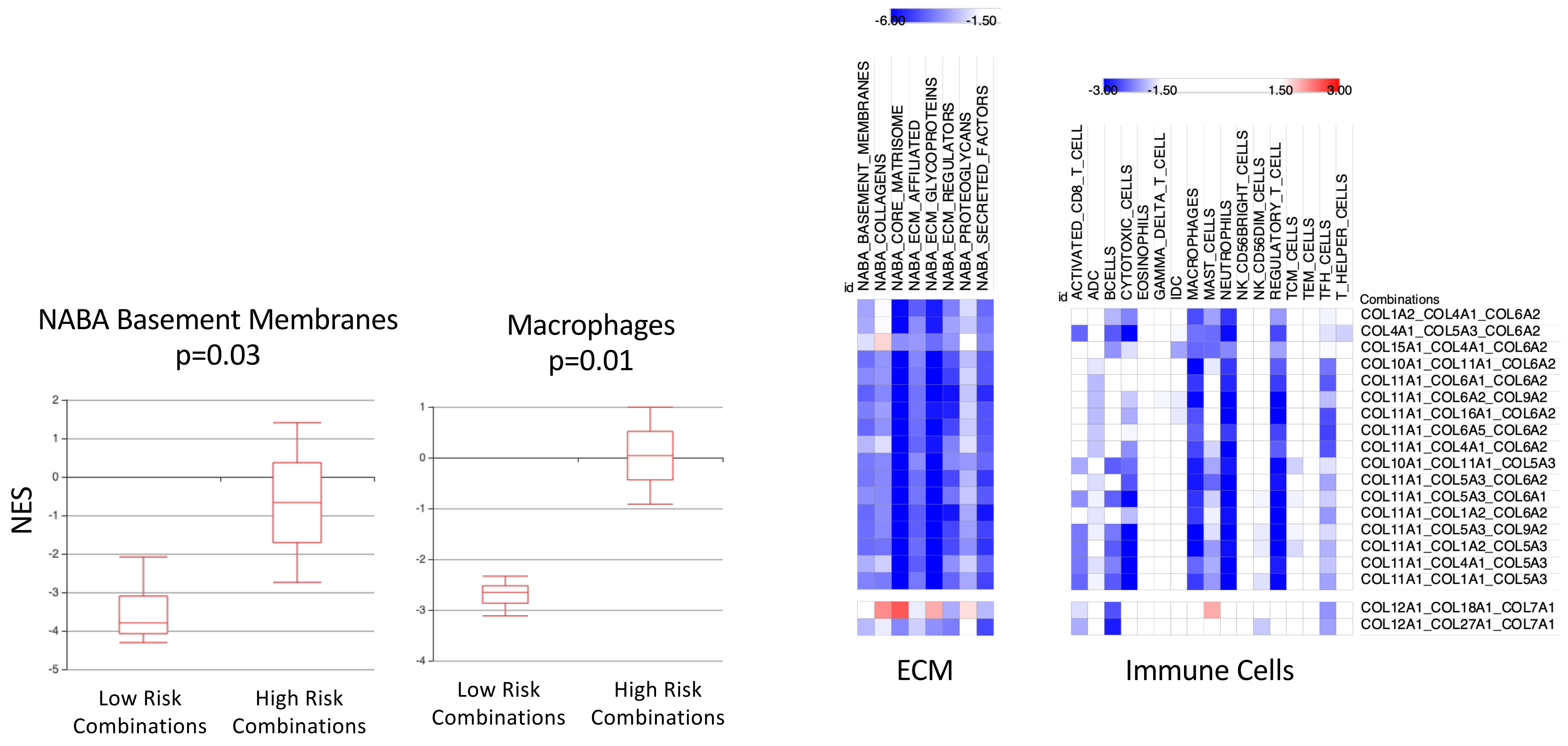

NABA Basement Membranes
p=0.03
NES
2
1
0
-1
-2
-3
-4
-5
Low Risk Combinations
High Risk Combinations
Macrophages
p=0.01
1
0
-1
-2
-3
-4
Low Risk Combinations
High Risk Combinations
-6.00
-1.50
id
NABA_BASEMENT_MEMBRANES
NABA_COLLAGENS
NABA_CORE_MATRISOME
NABA_ECM_AFFILIATED
NABA_ECM_GLYCOPROTEINS
NABA_ECM_REGULATORS
NABA_PROTEOGLYCANS
NABA_SECRETED_FACTORS
ECM
-3.00
-1.50
1.50
3.00
id
ACTIVATED_CD8_T_CELL
ADC
BCELLS
CYTOTOXIC_CELLS
EOSINOPHILS
GAMMA_DELTA_T_CELL
IDC
MACROPHAGES
MAST_CELLS
NEUTROPHILS
NK_CD56BRIGHT_CELLS
NK_CD56DIM_CELLS
REGULATORY_T_CELL
TCM_CELLS
TEM_CELLS
TFH_CELLS
T_HELPER_CELLS
Combinations
COL1A2_COL4A1_COL6A2
COL4A1_COL5A3_COL6A2
COL15A1_COL4A1_COL6A2
COL10A1_COL11A1_COL6A2
COL11A1_COL6A1_COL6A2
COL11A1_COL6A2_COL9A2
COL11A1_COL16A1_COL6A2
COL11A1_COL6A5_COL6A2
COL11A1_COL4A1_COL6A2
COL10A1_COL11A1_COL5A3
COL11A1_COL5A3_COL6A2
COL11A1_COL5A3_COL6A1
COL11A1_COL1A2_COL6A2
COL11A1_COL5A3_COL9A2
COL11A1_COL1A2_COL5A3
COL11A1_COL4A1_COL5A3
COL11A1_COL1A1_COL5A3
COL12A1_COL18A1_COL7A1
COL12A1_COL27A1_COL7A1
Immune Cells

Figure S8A

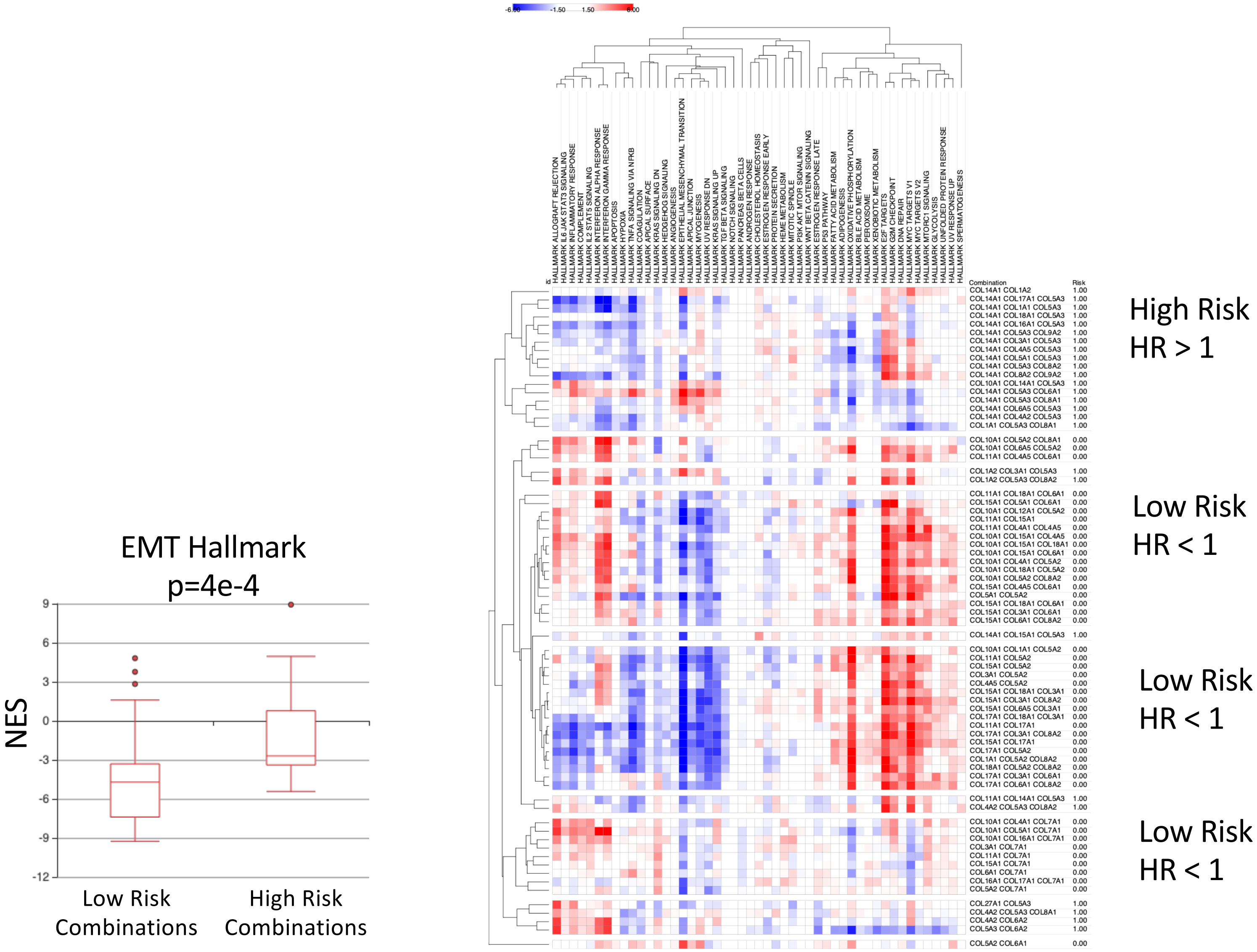

EMT Hallmark
p=4e-4
NES
9
6
3
0
-3
-6
-9
-12
Low Risk Combinations
High Risk Combinations
High Risk
HR > 1
Low Risk
HR < 1
Low Risk
HR < 1
Low Risk
HR < 1

Figure S8C

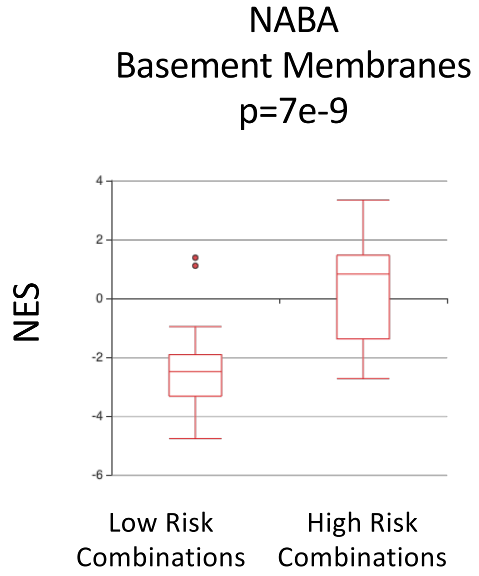
NABA
Basement Membranes
p=7e-9
NES
4
2
0
-2
-4
-6
Low Risk Combinations
High Risk Combinations

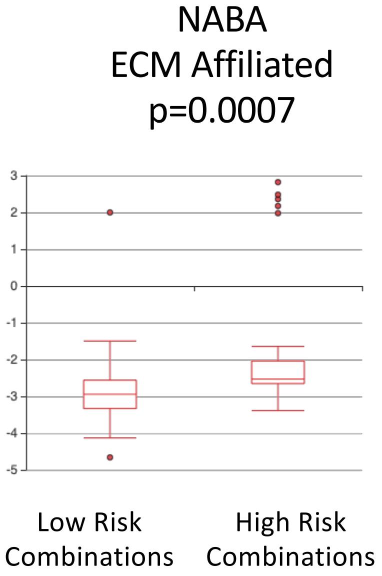
NABA
ECM Affiliated
p=0.0007
3
2
1
0
-1
-2
-3
-4
-5
Low Risk Combinations
High Risk Combinations

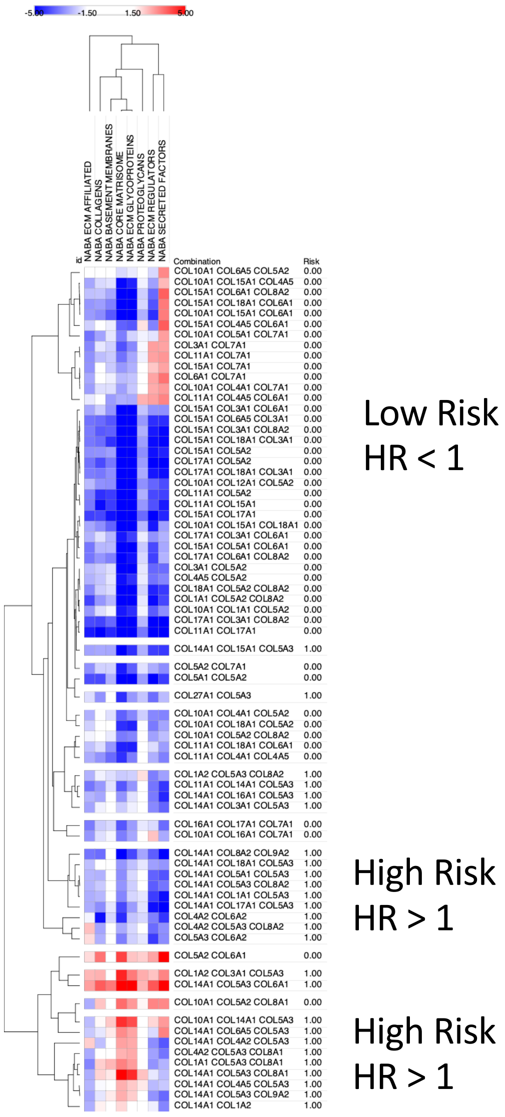
-5.00
-1.50
1.50
5.00
id
NABA ECM AFFILIATED
NABA COLLAGENS
NABA BASEMENT MEMBRANES
NABA CORE MATRISOME
NABA ECM GLYCOPROTEINS
NABA PROTEOGLYCANS
NABA ECM REGULATORS
NABA SECRETED FACTORS
Combination Risk
COL10A1 COL6A5 COL5A2 0.00
COL10A1 COL15A1 COL4A5 0.00
COL15A1 COL6A1 COL8A2 0.00
COL15A1 COL18A1 COL6A1 0.00
COL10A1 COL15A1 COL6A1 0.00
COL15A1 COL4A5 COL6A1 0.00
COL10A1 COL5A1 COL7A1 0.00
COL3A1 COL7A1 0.00
COL11A1 COL7A1 0.00
COL15A1 COL7A1 0.00
COL6A1 COL7A1 0.00
COL10A1 COL4A1 COL7A1 0.00
COL11A1 COL4A5 COL6A1 0.00
COL15A1 COL3A1 COL6A1 0.00
COL15A1 COL6A5 COL3A1 0.00
COL15A1 COL3A1 COL8A2 0.00
COL15A1 COL18A1 COL3A1 0.00
COL15A1 COL5A2 0.00
COL17A1 COL5A2 0.00
COL17A1 COL18A1 COL3A1 0.00
COL10A1 COL12A1 COL5A2 0.00
COL11A1 COL5A2 0.00
COL11A1 COL15A1 0.00
COL15A1 COL17A1 0.00
COL10A1 COL15A1 COL18A1 0.00
COL17A1 COL3A1 COL6A1 0.00
COL15A1 COL5A1 COL6A1 0.00
COL17A1 COL6A1 COL8A2 0.00
COL3A1 COL5A2 0.00
COL4A5 COL5A2 0.00
COL18A1 COL5A2 COL8A2 0.00
COL1A1 COL5A2 COL8A2 0.00
COL10A1 COL1A1 COL5A2 0.00
COL17A1 COL3A1 COL8A2 0.00
COL11A1 COL17A1 0.00
COL14A1 COL15A1 COL5A3 1.00
COL5A2 COL7A1 0.00
COL5A1 COL5A2 0.00
COL27A1 COL5A3 1.00
COL10A1 COL4A1 COL5A2 0.00
COL10A1 COL18A1 COL5A2 0.00
COL10A1 COL5A2 COL8A2 0.00
COL11A1 COL18A1 COL6A1 0.00
COL11A1 COL4A1 COL4A5 0.00
COL1A2 COL5A3 COL8A2 1.00
COL11A1 COL14A1 COL5A3 1.00
COL14A1 COL16A1 COL5A3 1.00
COL14A1 COL3A1 COL5A3 1.00
COL16A1 COL17A1 COL7A1 0.00
COL10A1 COL16A1 COL7A1 0.00
COL14A1 COL8A2 COL9A2 1.00
COL14A1 COL18A1 COL5A3 1.00
COL14A1 COL5A1 COL5A3 1.00
COL14A1 COL5A3 COL8A2 1.00
COL14A1 COL1A1 COL5A3 1.00
COL14A1 COL17A1 COL5A3 1.00
COL4A2 COL6A2 1.00
COL4A2 COL5A3 COL8A2 1.00
COL5A3 COL6A2 1.00
COL5A2 COL6A1 0.00
COL1A2 COL3A1 COL5A3 1.00
COL14A1 COL5A3 COL6A1 1.00
COL10A1 COL5A2 COL8A1 0.00
COL10A1 COL14A1 COL5A3 1.00
COL14A1 COL6A5 COL5A3 1.00
COL14A1 COL4A2 COL5A3 1.00
COL4A2 COL5A3 COL8A1 1.00
COL1A1 COL5A3 COL8A1 1.00
COL14A1 COL5A3 COL8A1 1.00
COL14A1 COL4A5 COL5A3 1.00
COL14A1 COL5A3 COL9A2 1.00
COL14A1 COL1A2 1.00
Low Risk
HR < 1
High Risk
HR > 1
High Risk
HR > 1

Figure S8C

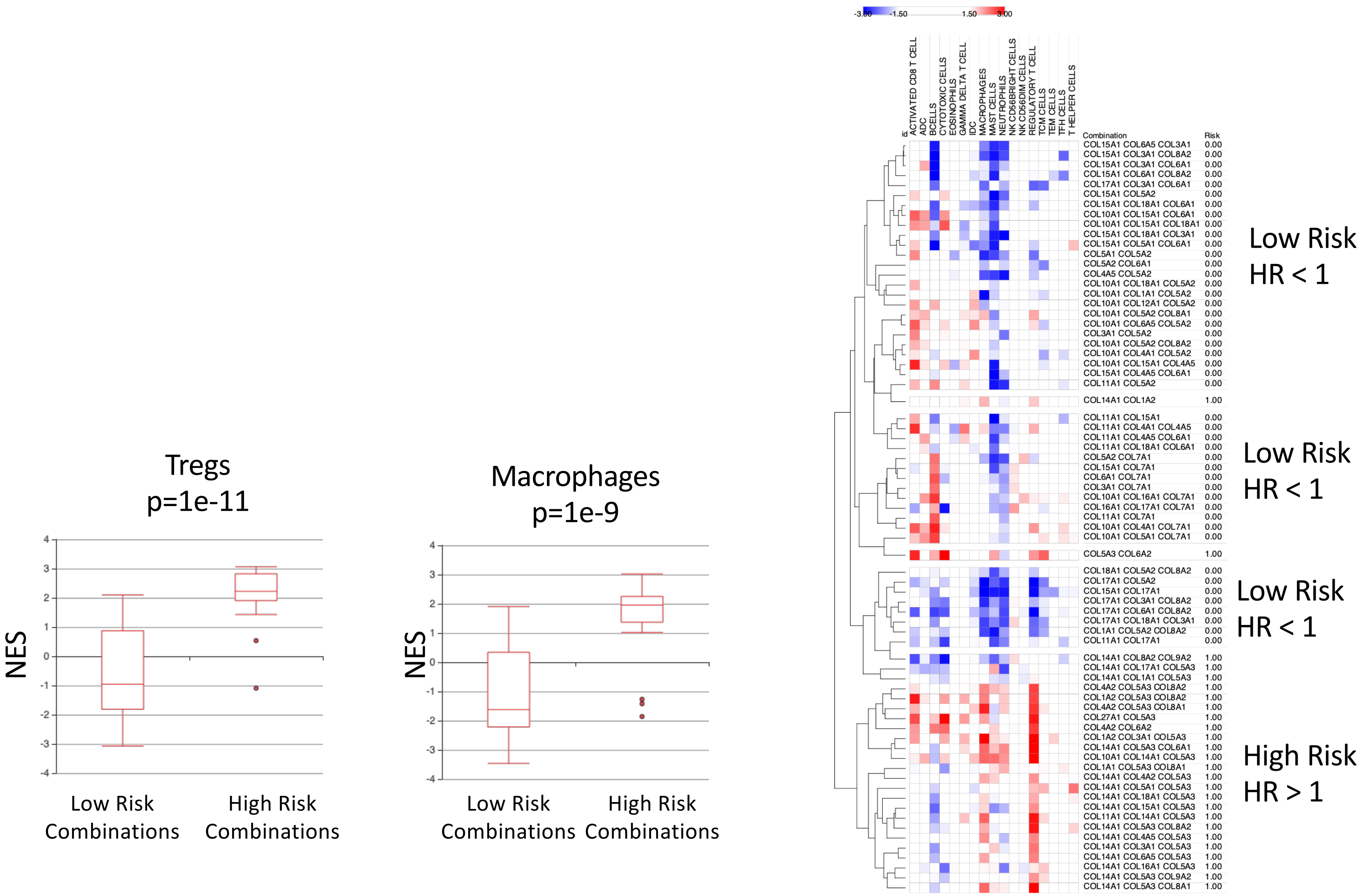

Tregs
p=1e-11
NES
4
3
2
1
0
-1
-2
-3
-4
Low Risk Combinations
High Risk Combinations
Macrophages
p=1e-9
NES
4
3
2
1
0
-1
-2
-3
-4
Low Risk Combinations
High Risk Combinations
-3.00
-1.50
1.50
3.00
id
ACTIVATED CD8 T CELL
ADC
BCELLS
CYTOTOXIC CELLS
EOSINOPHILS
GAMMA DELTA T CELL
IDC
MACROPHAGES
MAST CELLS
NEUTROPHILS
NK CD56BRIGHT CELLS
NK CD56DIM CELLS
REGULATORY T CELL
TCM CELLS
TEM CELLS
TFH CELLS
T HELPER CELLS
Combination
Risk
COL15A1 COL6A5 COL3A1 0.00
COL15A1 COL3A1 COL8A2 0.00
COL15A1 COL3A1 COL6A1 0.00
COL15A1 COL6A1 COL8A2 0.00
COL17A1 COL3A1 COL6A1 0.00
COL15A1 COL5A2 0.00
COL15A1 COL18A1 COL6A1 0.00
COL10A1 COL15A1 COL6A1 0.00
COL10A1 COL15A1 COL18A1 0.00
COL15A1 COL18A1 COL3A1 0.00
COL15A1 COL5A1 COL6A1 0.00
COL5A1 COL5A2 0.00
COL5A2 COL6A1 0.00
COL4A5 COL5A2 0.00
COL10A1 COL18A1 COL5A2 0.00
COL10A1 COL1A1 COL5A2 0.00
COL10A1 COL12A1 COL5A2 0.00
COL10A1 COL5A2 COL8A1 0.00
COL10A1 COL6A5 COL5A2 0.00
COL3A1 COL5A2 0.00
COL10A1 COL5A2 COL8A2 0.00
COL10A1 COL4A1 COL5A2 0.00
COL10A1 COL15A1 COL4A5 0.00
COL15A1 COL4A5 COL6A1 0.00
COL11A1 COL5A2 0.00
COL14A1 COL1A2 1.00
COL11A1 COL15A1 0.00
COL11A1 COL4A1 COL4A5 0.00
COL11A1 COL4A5 COL6A1 0.00
COL11A1 COL18A1 COL6A1 0.00
COL5A2 COL7A1 0.00
COL15A1 COL7A1 0.00
COL6A1 COL7A1 0.00
COL3A1 COL7A1 0.00
COL10A1 COL16A1 COL7A1 0.00
COL16A1 COL17A1 COL7A1 0.00
COL11A1 COL7A1 0.00
COL10A1 COL4A1 COL7A1 0.00
COL10A1 COL5A1 COL7A1 0.00
COL5A3 COL6A2 1.00
COL18A1 COL5A2 COL8A2 0.00
COL17A1 COL5A2 0.00
COL15A1 COL17A1 0.00
COL17A1 COL3A1 COL8A2 0.00
COL17A1 COL6A1 COL8A2 0.00
COL17A1 COL18A1 COL3A1 0.00
COL1A1 COL5A2 COL8A2 0.00
COL11A1 COL17A1 0.00
COL14A1 COL8A2 COL9A2 1.00
COL14A1 COL17A1 COL5A3 1.00
COL14A1 COL1A1 COL5A3 1.00
COL4A2 COL5A3 COL8A2 1.00
COL1A2 COL5A3 COL8A2 1.00
COL4A2 COL5A3 COL8A1 1.00
COL27A1 COL5A3 1.00
COL4A2 COL6A2 1.00
COL1A2 COL3A1 COL5A3 1.00
COL14A1 COL5A3 COL6A1 1.00
COL10A1 COL14A1 COL5A3 1.00
COL1A1 COL5A3 COL8A1 1.00
COL14A1 COL4A2 COL5A3 1.00
COL14A1 COL5A1 COL5A3 1.00
COL14A1 COL18A1 COL5A3 1.00
COL14A1 COL15A1 COL5A3 1.00
COL11A1 COL14A1 COL5A3 1.00
COL14A1 COL5A3 COL8A2 1.00
COL14A1 COL4A5 COL5A3 1.00
COL14A1 COL3A1 COL5A3 1.00
COL14A1 COL6A5 COL5A3 1.00
COL14A1 COL16A1 COL5A3 1.00
COL14A1 COL5A3 COL9A2 1.00
COL14A1 COL5A3 COL8A1 1.00
Low Risk
HR < 1
Low Risk
HR < 1
Low Risk
HR < 1
High Risk
HR > 1

Figure S9A

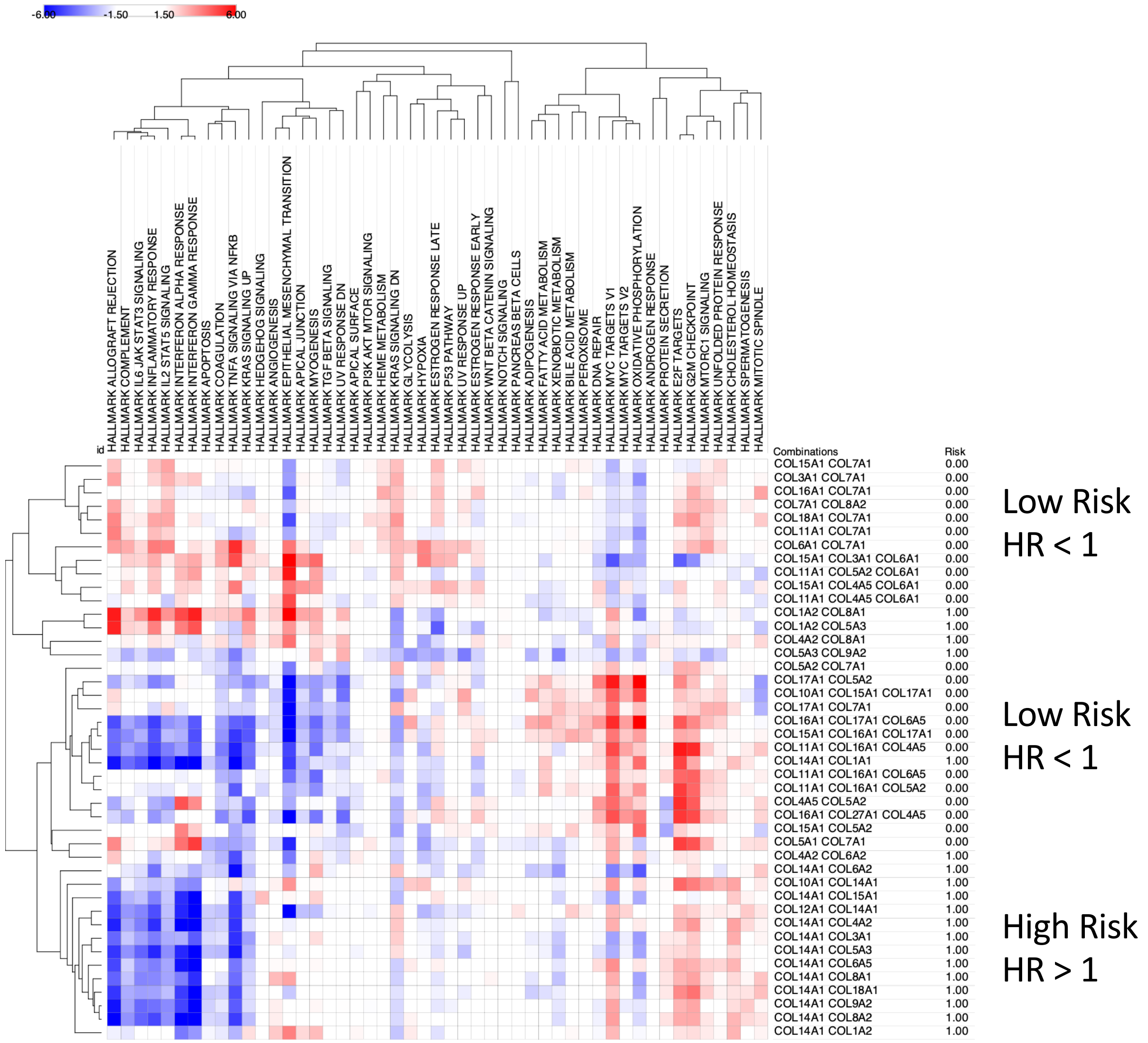
-6.00
-1.50
1.50
6.00
id
HALLMARK ALLOGRAFT REJECTION
HALLMARK COMPLEMENT
HALLMARK IL6 JAK STAT3 SIGNALING
HALLMARK INFLAMMATORY RESPONSE
HALLMARK IL2 STAT5 SIGNALING
HALLMARK INTERFERON ALPHA RESPONSE
HALLMARK INTERFERON GAMMA RESPONSE
HALLMARK APOPTOSIS
HALLMARK COAGULATION
HALLMARK TNFA SIGNALING VIA NFKB
HALLMARK KRAS SIGNALING UP
HALLMARK HEDGEHOG SIGNALING
HALLMARK ANGIOGENESIS
HALLMARK EPITHELIAL MESENCHYMAL TRANSITION
HALLMARK APICAL JUNCTION
HALLMARK MYOGENESIS
HALLMARK TGF BETA SIGNALING
HALLMARK UV RESPONSE DN
HALLMARK APICAL SURFACE
HALLMARK PI3K AKT MTOR SIGNALING
HALLMARK HEME METABOLISM
HALLMARK KRAS SIGNALING DN
HALLMARK GLYCOLYSIS
HALLMARK HYPOXIA
HALLMARK ESTROGEN RESPONSE LATE
HALLMARK P53 PATHWAY
HALLMARK UV RESPONSE UP
HALLMARK ESTROGEN RESPONSE EARLY
HALLMARK WNT BETA CATENIN SIGNALING
HALLMARK NOTCH SIGNALING
HALLMARK PANCREAS BETA CELLS
HALLMARK ADIPOGENESIS
HALLMARK FATTY ACID METABOLISM
HALLMARK XENOBIOTIC METABOLISM
HALLMARK BILE ACID METABOLISM
HALLMARK PEROXISOME
HALLMARK DNA REPAIR
HALLMARK MYC TARGETS V1
HALLMARK MYC TARGETS V2
HALLMARK OXIDATIVE PHOSPHORYLATION
HALLMARK ANDROGEN RESPONSE
HALLMARK PROTEIN SECRETION
HALLMARK E2F TARGETS
HALLMARK G2M CHECKPOINT
HALLMARK MTORC1 SIGNALING
HALLMARK UNFOLDED PROTEIN RESPONSE
HALLMARK CHOLESTEROL HOMEOSTASIS
HALLMARK SPERMATOGENESIS
HALLMARK MITOTIC SPINDLE
Combinations Risk
COL15A1 COL7A1 0.00
COL3A1 COL7A1 0.00
COL16A1 COL7A1 0.00
COL7A1 COL8A2 0.00
COL18A1 COL7A1 0.00
COL11A1 COL7A1 0.00
COL6A1 COL7A1 0.00
COL15A1 COL3A1 COL6A1 0.00
COL11A1 COL5A2 COL6A1 0.00
COL15A1 COL4A5 COL6A1 0.00
COL11A1 COL4A5 COL6A1 0.00
COL1A2 COL8A1 1.00
COL1A2 COL5A3 1.00
COL4A2 COL8A1 1.00
COL5A3 COL9A2 1.00
COL5A2 COL7A1 0.00
COL17A1 COL5A2 0.00
COL10A1 COL15A1 COL17A1 0.00
COL17A1 COL7A1 0.00
COL16A1 COL17A1 COL6A5 0.00
COL15A1 COL16A1 COL17A1 0.00
COL11A1 COL16A1 COL4A5 0.00
COL14A1 COL1A1 1.00
COL11A1 COL16A1 COL6A5 0.00
COL11A1 COL16A1 COL5A2 0.00
COL4A5 COL5A2 0.00
COL16A1 COL27A1 COL4A5 0.00
COL15A1 COL5A2 0.00
COL5A1 COL7A1 0.00
COL4A2 COL6A2 1.00
COL14A1 COL6A2 1.00
COL10A1 COL14A1 1.00
COL14A1 COL15A1 1.00
COL12A1 COL14A1 1.00
COL14A1 COL4A2 1.00
COL14A1 COL3A1 1.00
COL14A1 COL5A3 1.00
COL14A1 COL6A5 1.00
COL14A1 COL8A1 1.00
COL14A1 COL18A1 1.00
COL14A1 COL9A2 1.00
COL14A1 COL8A2 1.00
COL14A1 COL1A2 1.00
Low Risk
HR < 1
Low Risk
HR < 1
High Risk
HR > 1

Figure S9B

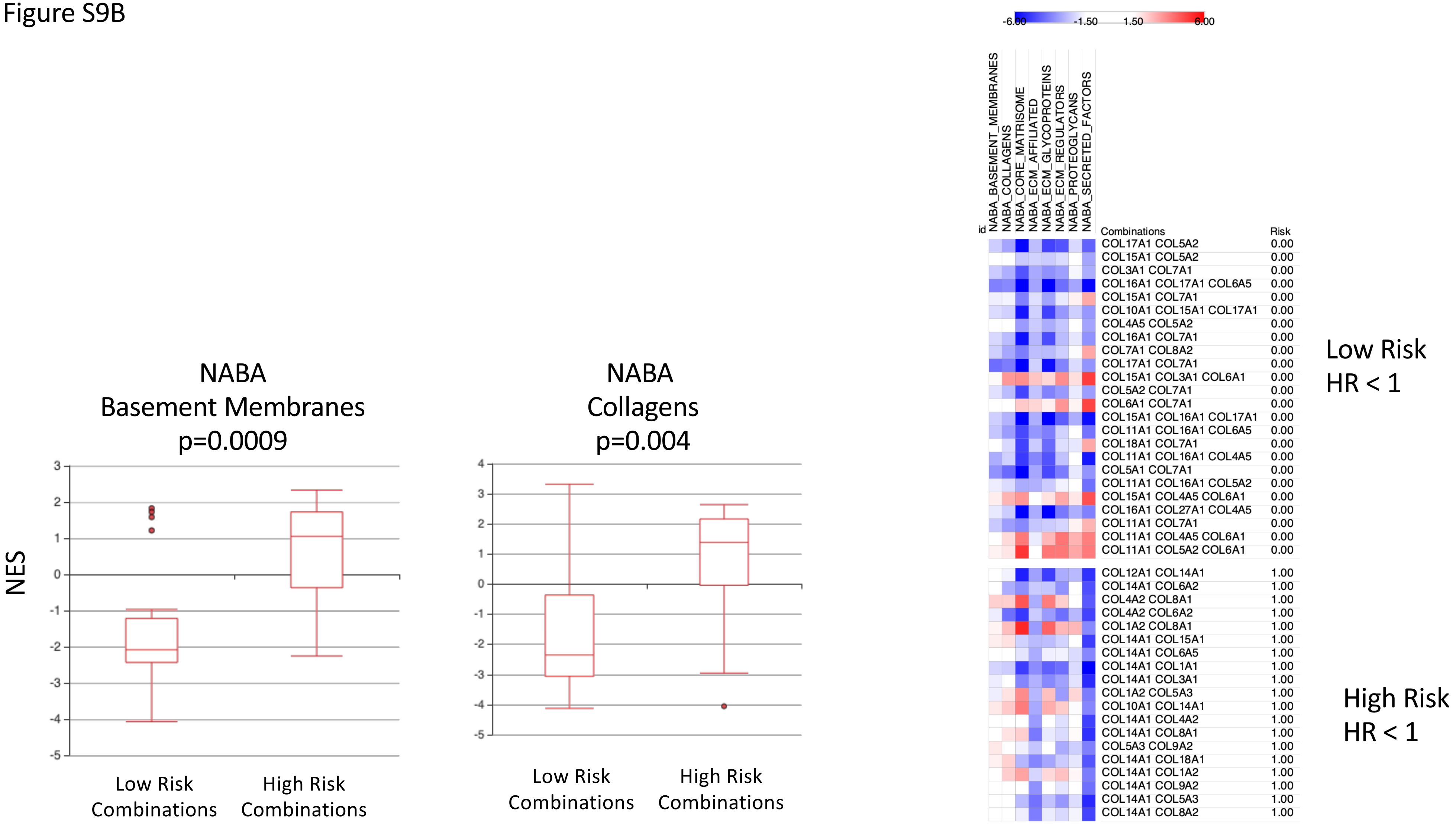
NABA
Basement Membranes
p=0.0009
NES
3
2
1
0
-1
-2
-3
-4
-5
Low Risk
Combinations
High Risk
Combinations
NABA
Collagens
p=0.004
4
3
2
1
0
-1
-2
-3
-4
-5
Low Risk
Combinations
High Risk
Combinations
-6.00
-1.50
1.50
6.00
id
NABA_BASEMENT_MEMBRANES
NABA_COLLAGENS
NABA_CORE_MATRISOME
NABA_ECM_AFFILIATED
NABA_ECM_GLYCOPROTEINS
NABA_ECM_REGULATORS
NABA_PROTEOGLYCANS
NABA_SECRETED_FACTORS
Combinations
Risk
COL17A1 COL5A2 0.00
COL15A1 COL5A2 0.00
COL3A1 COL7A1 0.00
COL16A1 COL17A1 COL6A5 0.00
COL15A1 COL7A1 0.00
COL10A1 COL15A1 COL17A1 0.00
COL4A5 COL5A2 0.00
COL16A1 COL7A1 0.00
COL7A1 COL8A2 0.00
COL17A1 COL7A1 0.00
COL15A1 COL3A1 COL6A1 0.00
COL5A2 COL7A1 0.00
COL6A1 COL7A1 0.00
COL15A1 COL16A1 COL17A1 0.00
COL11A1 COL16A1 COL6A5 0.00
COL18A1 COL7A1 0.00
COL11A1 COL16A1 COL4A5 0.00
COL5A1 COL7A1 0.00
COL11A1 COL16A1 COL5A2 0.00
COL15A1 COL4A5 COL6A1 0.00
COL16A1 COL27A1 COL4A5 0.00
COL11A1 COL7A1 0.00
COL11A1 COL4A5 COL6A1 0.00
COL11A1 COL5A2 COL6A1 0.00
COL12A1 COL14A1 1.00
COL14A1 COL6A2 1.00
COL4A2 COL8A1 1.00
COL4A2 COL6A2 1.00
COL1A2 COL8A1 1.00
COL14A1 COL15A1 1.00
COL14A1 COL6A5 1.00
COL14A1 COL1A1 1.00
COL14A1 COL3A1 1.00
COL1A2 COL5A3 1.00
COL10A1 COL14A1 1.00
COL14A1 COL4A2 1.00
COL14A1 COL8A1 1.00
COL5A3 COL9A2 1.00
COL14A1 COL18A1 1.00
COL14A1 COL1A2 1.00
COL14A1 COL9A2 1.00
COL14A1 COL5A3 1.00
COL14A1 COL8A2 1.00
Low Risk
HR < 1
High Risk
HR < 1

Figure S9C

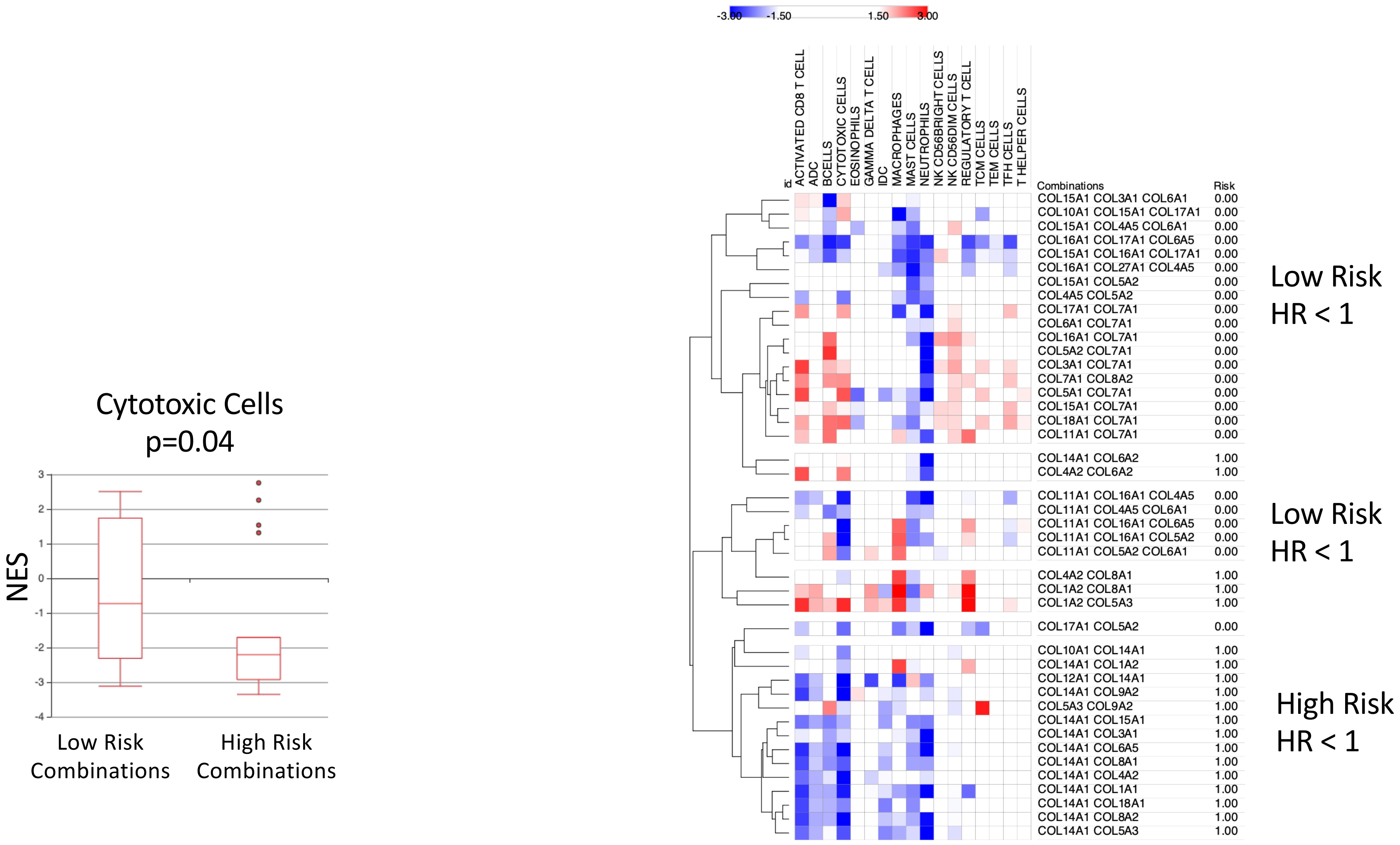
Cytotoxic Cells
p=0.04
NES
3
2
1
0
-1
-2
-3
-4
Low Risk Combinations
High Risk Combinations
-3.00
-1.50
1.50
3.00
id
ACTIVATED CD8 T CELL
ADC
BCELLS
CYTOTOXIC CELLS
EOSINOPHILS
GAMMA DELTA T CELL
IDC
MACROPHAGES
MAST CELLS
NEUTROPHILS
NK CD56BRIGHT CELLS
NK CD56DIM CELLS
REGULATORY T CELL
TCM CELLS
TEM CELLS
TFH CELLS
T HELPER CELLS
Combinations
Risk
COL15A1 COL3A1 COL6A1 0.00
COL10A1 COL15A1 COL17A1 0.00
COL15A1 COL4A5 COL6A1 0.00
COL16A1 COL17A1 COL6A5 0.00
COL15A1 COL16A1 COL17A1 0.00
COL16A1 COL27A1 COL4A5 0.00
COL15A1 COL5A2 0.00
COL4A5 COL5A2 0.00
COL17A1 COL7A1 0.00
COL6A1 COL7A1 0.00
COL16A1 COL7A1 0.00
COL5A2 COL7A1 0.00
COL3A1 COL7A1 0.00
COL7A1 COL8A2 0.00
COL5A1 COL7A1 0.00
COL15A1 COL7A1 0.00
COL18A1 COL7A1 0.00
COL11A1 COL7A1 0.00
COL14A1 COL6A2 1.00
COL4A2 COL6A2 1.00
COL11A1 COL16A1 COL4A5 0.00
COL11A1 COL4A5 COL6A1 0.00
COL11A1 COL16A1 COL6A5 0.00
COL11A1 COL16A1 COL5A2 0.00
COL11A1 COL5A2 COL6A1 0.00
COL4A2 COL8A1 1.00
COL1A2 COL8A1 1.00
COL1A2 COL5A3 1.00
COL17A1 COL5A2 0.00
COL10A1 COL14A1 1.00
COL14A1 COL1A2 1.00
COL12A1 COL14A1 1.00
COL14A1 COL9A2 1.00
COL5A3 COL9A2 1.00
COL14A1 COL15A1 1.00
COL14A1 COL3A1 1.00
COL14A1 COL6A5 1.00
COL14A1 COL8A1 1.00
COL14A1 COL4A2 1.00
COL14A1 COL1A1 1.00
COL14A1 COL18A1 1.00
COL14A1 COL8A2 1.00
COL14A1 COL5A3 1.00
Low Risk
HR < 1
Low Risk
HR < 1
High Risk
HR < 1

Figure S10A

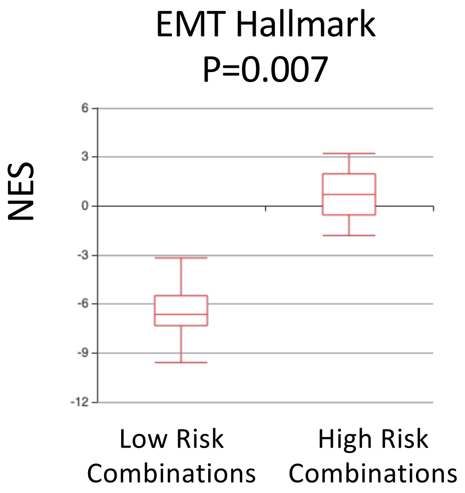
EMT Hallmark
P=0.007
NES
6
3
0
-3
-6
-9
-12
Low Risk Combinations
High Risk Combinations

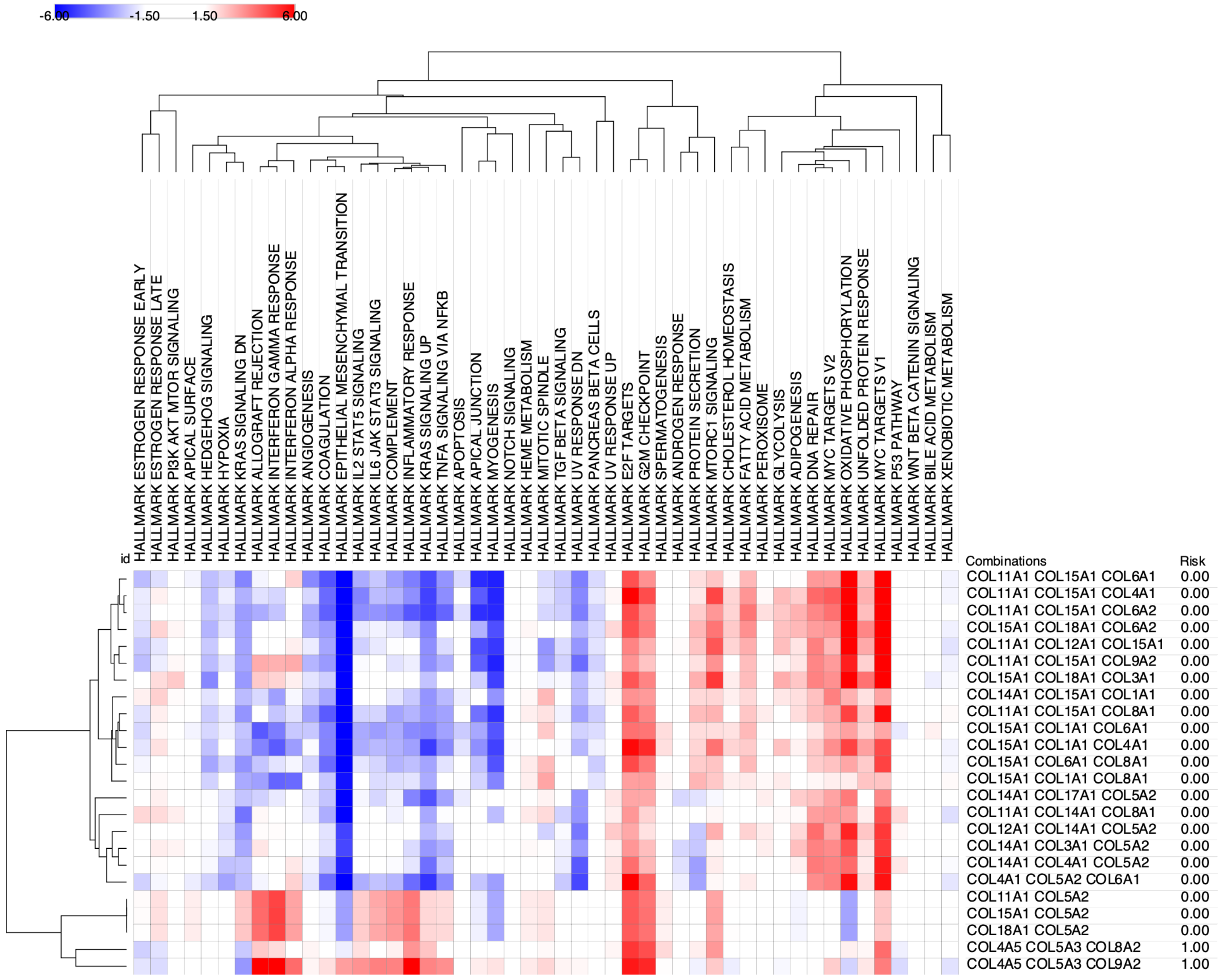
-6.00
-1.50
1.50
6.00
id
HALLMARK ESTROGEN RESPONSE EARLY
HALLMARK ESTROGEN RESPONSE LATE
HALLMARK PI3K AKT MTOR SIGNALING
HALLMARK APICAL SURFACE
HALLMARK HEDGEHOG SIGNALING
HALLMARK HYPOXIA
HALLMARK KRAS SIGNALING DN
HALLMARK ALLOGRAFT REJECTION
HALLMARK INTERFERON GAMMA RESPONSE
HALLMARK INTERFERON ALPHA RESPONSE
HALLMARK ANGIOGENESIS
HALLMARK COAGULATION
HALLMARK EPITHELIAL MESENCHYMAL TRANSITION
HALLMARK IL2 STAT5 SIGNALING
HALLMARK IL6 JAK STAT3 SIGNALING
HALLMARK COMPLEMENT
HALLMARK INFLAMMATORY RESPONSE
HALLMARK KRAS SIGNALING UP
HALLMARK TNFA SIGNALING VIA NFKB
HALLMARK APOPTOSIS
HALLMARK APICAL JUNCTION
HALLMARK MYOGENESIS
HALLMARK NOTCH SIGNALING
HALLMARK HEME METABOLISM
HALLMARK MITOTIC SPINDLE
HALLMARK TGF BETA SIGNALING
HALLMARK UV RESPONSE DN
HALLMARK PANCREAS BETA CELLS
HALLMARK UV RESPONSE UP
HALLMARK E2F TARGETS
HALLMARK G2M CHECKPOINT
HALLMARK SPERMATOGENESIS
HALLMARK ANDROGEN RESPONSE
HALLMARK PROTEIN SECRETION
HALLMARK MTORC1 SIGNALING
HALLMARK CHOLESTEROL HOMEOSTASIS
HALLMARK FATTY ACID METABOLISM
HALLMARK PEROXISOME
HALLMARK GLYCOLYSIS
HALLMARK ADIPOGENESIS
HALLMARK DNA REPAIR
HALLMARK MYC TARGETS V2
HALLMARK OXIDATIVE PHOSPHORYLATION
HALLMARK UNFOLDED PROTEIN RESPONSE
HALLMARK MYC TARGETS V1
HALLMARK P53 PATHWAY
HALLMARK WNT BETA CATENIN SIGNALING
HALLMARK BILE ACID METABOLISM
HALLMARK XENOBIOTIC METABOLISM
Combinations
Risk
COL11A1 COL15A1 COL6A1 0.00
COL11A1 COL15A1 COL4A1 0.00
COL11A1 COL15A1 COL6A2 0.00
COL15A1 COL18A1 COL6A2 0.00
COL11A1 COL12A1 COL15A1 0.00
COL11A1 COL15A1 COL9A2 0.00
COL15A1 COL18A1 COL3A1 0.00
COL14A1 COL15A1 COL1A1 0.00
COL11A1 COL15A1 COL8A1 0.00
COL15A1 COL1A1 COL6A1 0.00
COL15A1 COL1A1 COL4A1 0.00
COL15A1 COL6A1 COL8A1 0.00
COL15A1 COL1A1 COL8A1 0.00
COL14A1 COL17A1 COL5A2 0.00
COL11A1 COL14A1 COL8A1 0.00
COL12A1 COL14A1 COL5A2 0.00
COL14A1 COL3A1 COL5A2 0.00
COL14A1 COL4A1 COL5A2 0.00
COL4A1 COL5A2 COL6A1 0.00
COL11A1 COL5A2 0.00
COL15A1 COL5A2 0.00
COL18A1 COL5A2 0.00
COL4A5 COL5A3 COL8A2 1.00
COL4A5 COL5A3 COL9A2 1.00

Figure S10B

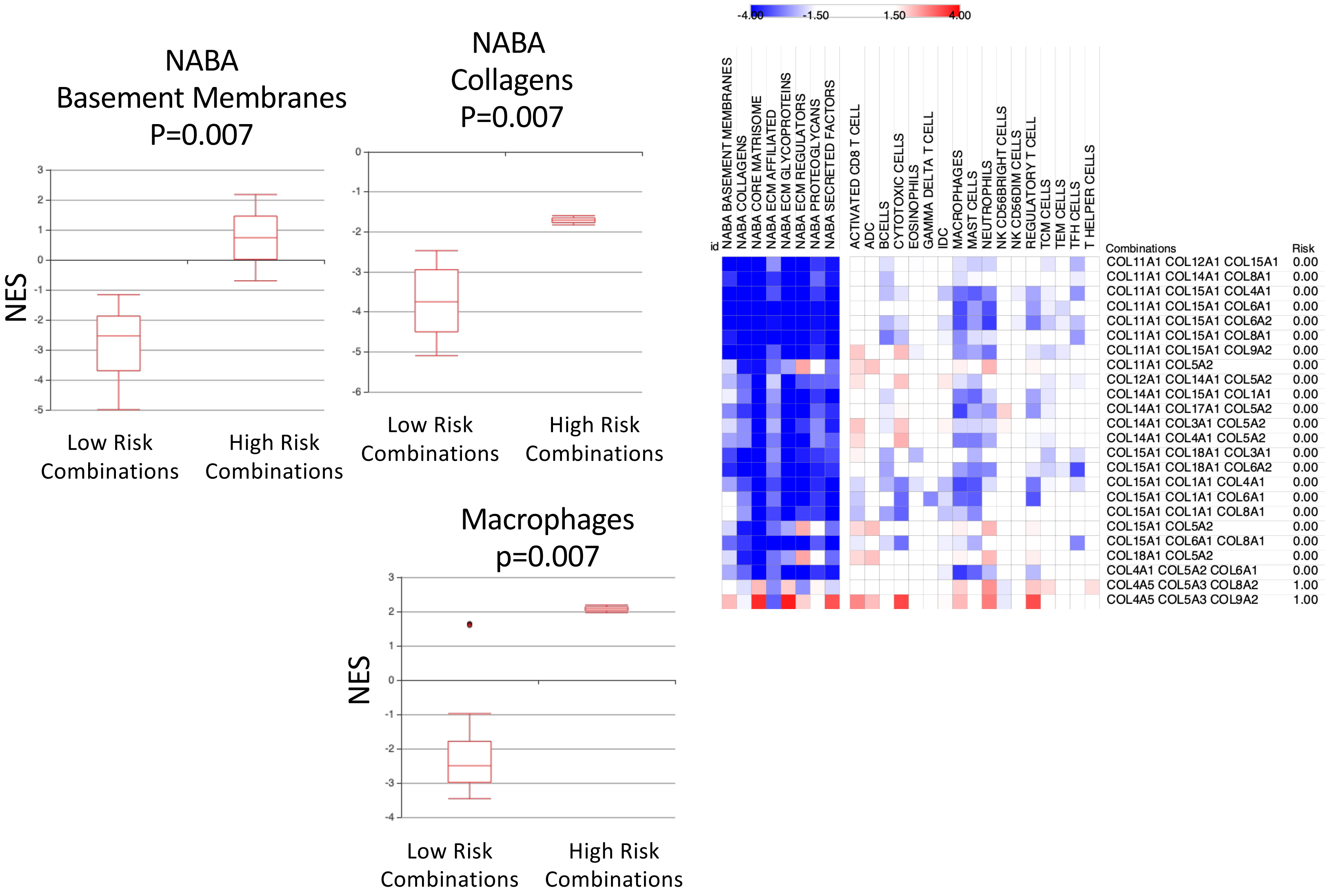
NABA
Basement Membranes
P=0.007
NES
Low Risk Combinations
High Risk Combinations
NABA
Collagens
P=0.007
Low Risk Combinations
High Risk Combinations
Macrophages
p=0.007
NES
Low Risk Combinations
High Risk Combinations
-4.00
-1.50
1.50
4.00
id
NABA BASEMENT MEMBRANES
NABA COLLAGENS
NABA CORE MATRISOME
NABA ECM AFFILIATED
NABA ECM GLYCOPROTEINS
NABA ECM REGULATORS
NABA PROTEOGLYCANS
NABA SECRETED FACTORS
ACTIVATED CD8 T CELL
ADC
BCELLS
CYTOTOXIC CELLS
EOSINOPHILS
GAMMA DELTA T CELL
IDC
MACROPHAGES
MAST CELLS
NEUTROPHILS
NK CD56BRIGHT CELLS
NK CD56DIM CELLS
REGULATORY T CELL
TCM CELLS
TEM CELLS
TFH CELLS
T HELPER CELLS
Combinations
Risk
COL11A1 COL12A1 COL15A1 0.00
COL11A1 COL14A1 COL8A1 0.00
COL11A1 COL15A1 COL4A1 0.00
COL11A1 COL15A1 COL6A1 0.00
COL11A1 COL15A1 COL6A2 0.00
COL11A1 COL15A1 COL8A1 0.00
COL11A1 COL15A1 COL9A2 0.00
COL11A1 COL5A2 0.00
COL12A1 COL14A1 COL5A2 0.00
COL14A1 COL15A1 COL1A1 0.00
COL14A1 COL17A1 COL5A2 0.00
COL14A1 COL3A1 COL5A2 0.00
COL14A1 COL4A1 COL5A2 0.00
COL15A1 COL18A1 COL3A1 0.00
COL15A1 COL18A1 COL6A2 0.00
COL15A1 COL1A1 COL4A1 0.00
COL15A1 COL1A1 COL6A1 0.00
COL15A1 COL1A1 COL8A1 0.00
COL15A1 COL5A2 0.00
COL15A1 COL6A1 COL8A1 0.00
COL18A1 COL5A2 0.00
COL4A1 COL5A2 COL6A1 0.00
COL4A5 COL5A3 COL8A2 1.00
COL4A5 COL5A3 COL9A2 1.00

Figure S11A

A.

B.

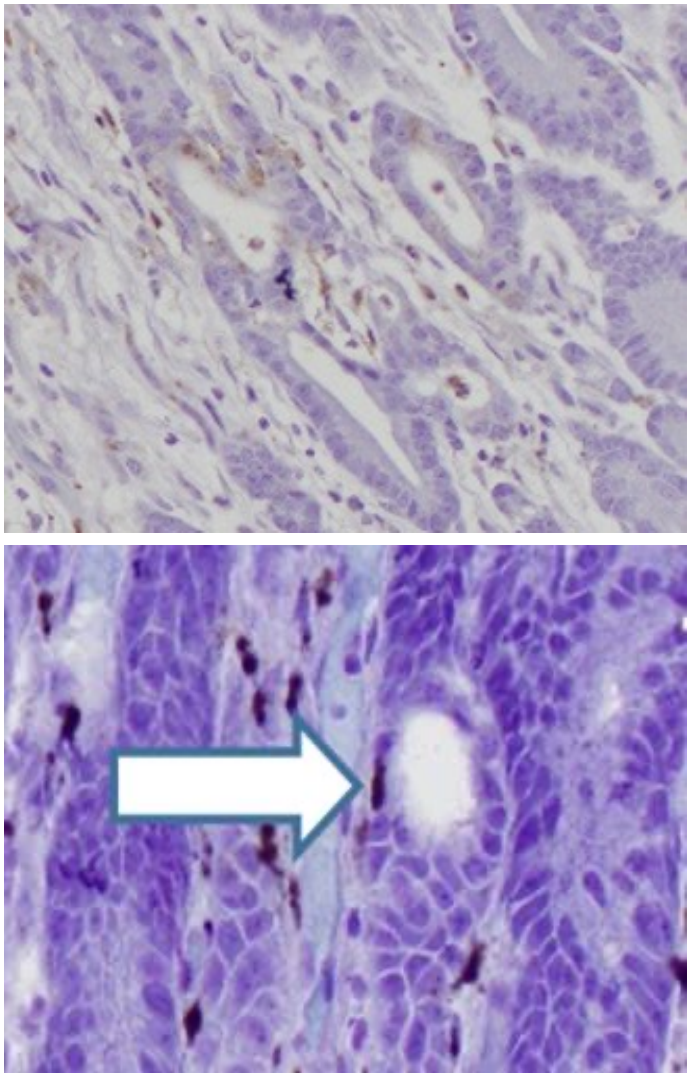

Figure S11B

COL7A1 Epithelium Expression

COL7A1 Stroma Expression

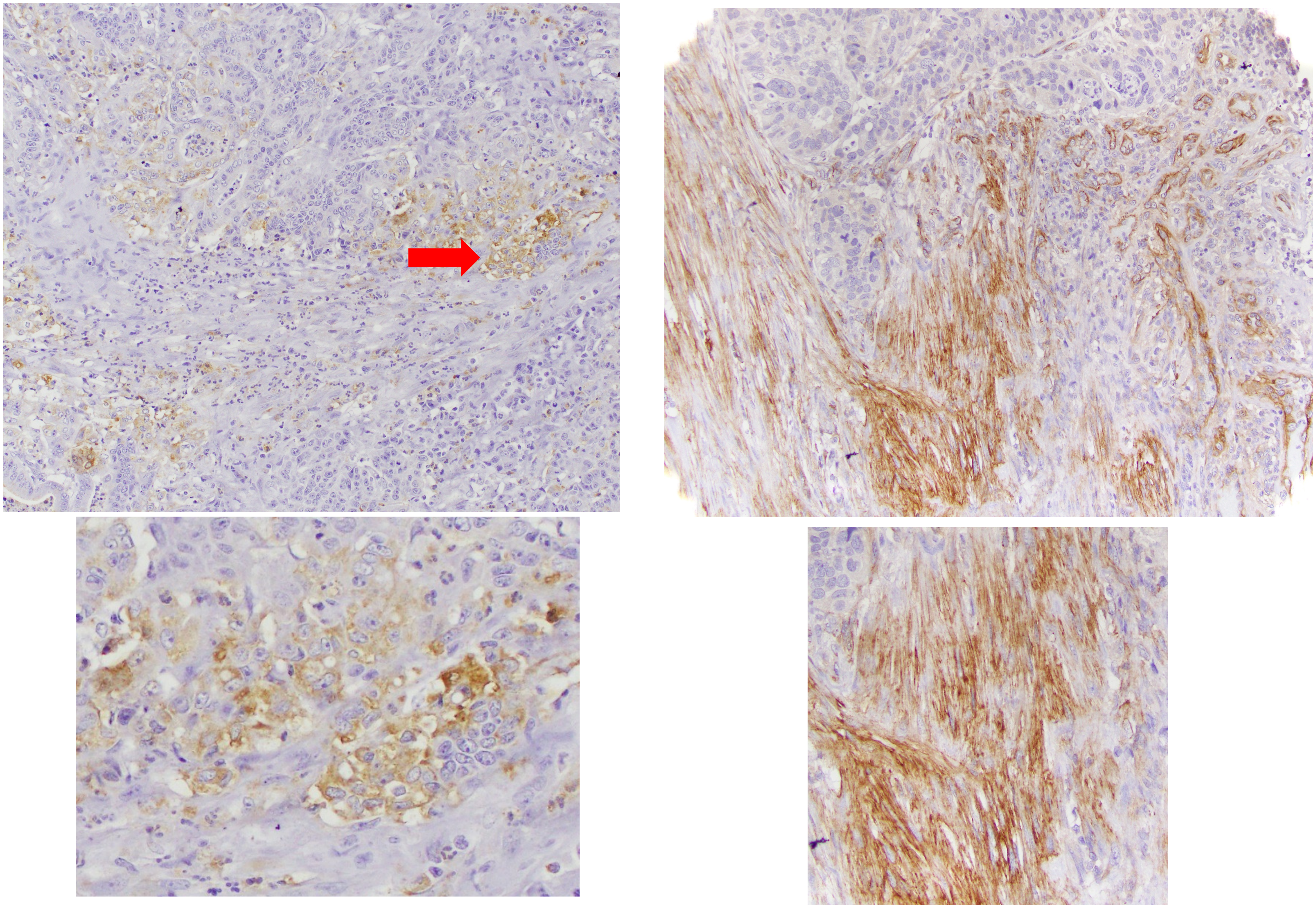

Figure S11C

COL7 expression in normal skin

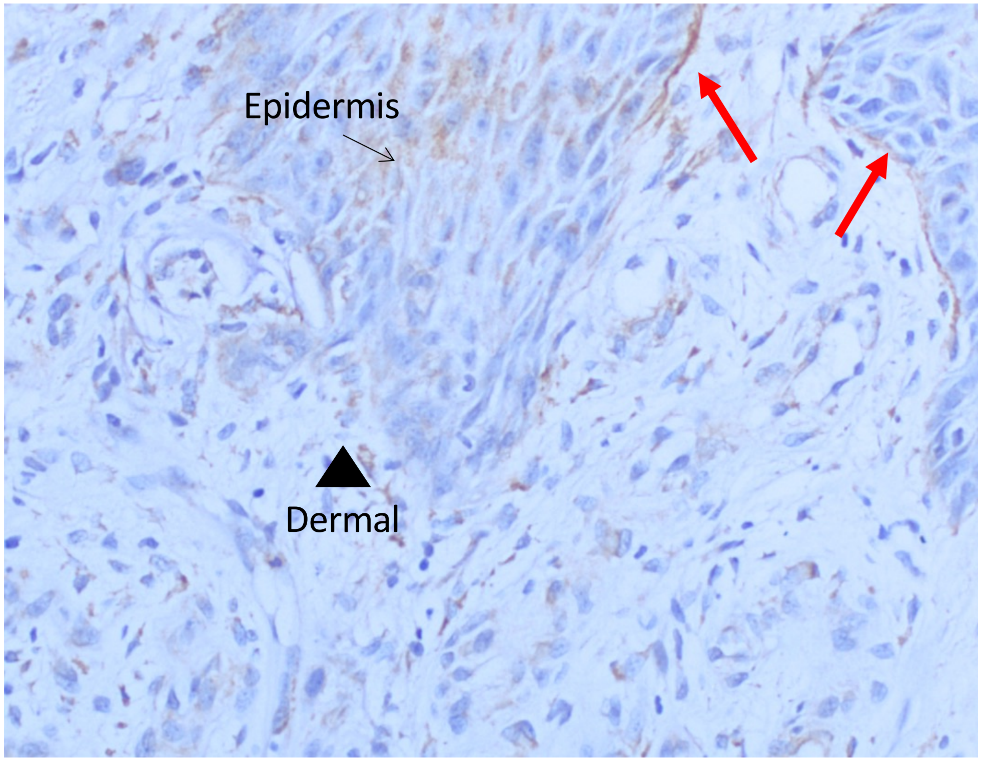

Figure S12A

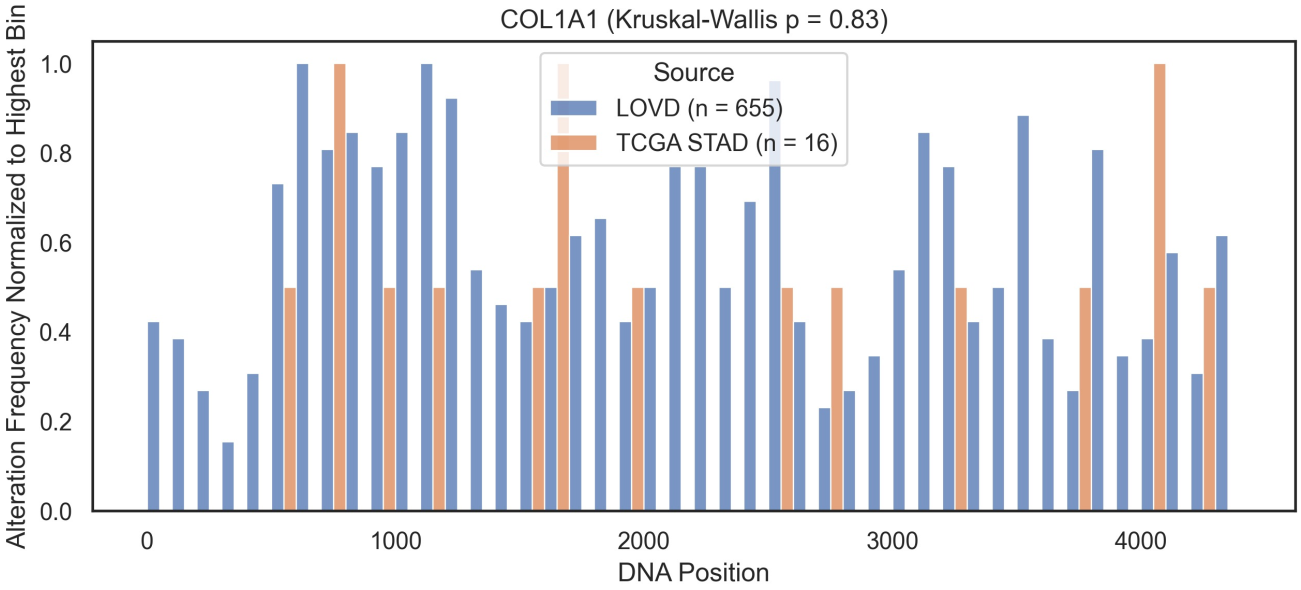
COL1A1 (Kruskal-Wallis p = 0.83)
Alteration Frequency Normalized to Highest Bin
1.0
0.8
0.6
0.4
0.2
0.0
0
1000
2000
3000
4000
DNA Position
Source
LOVD (n = 655)
TCGA STAD (n = 16)

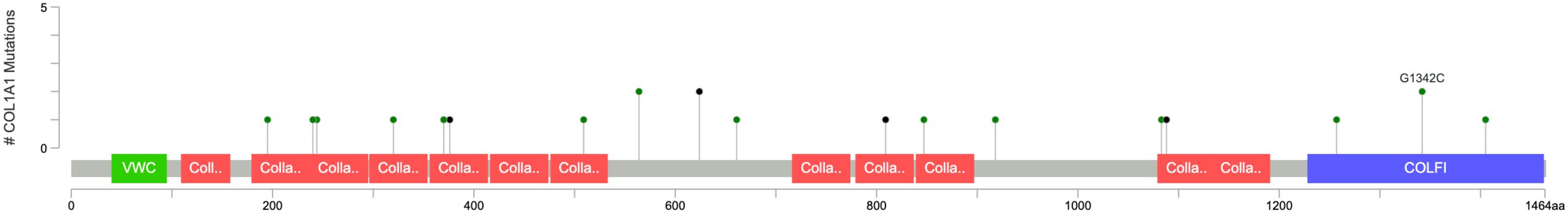
### COL1A1 Mutations
5
0
G1342C
VWC
Coll..
Colla..
COLFI
0
200
400
600
800
1000
1200
1464aa

Figure S12B

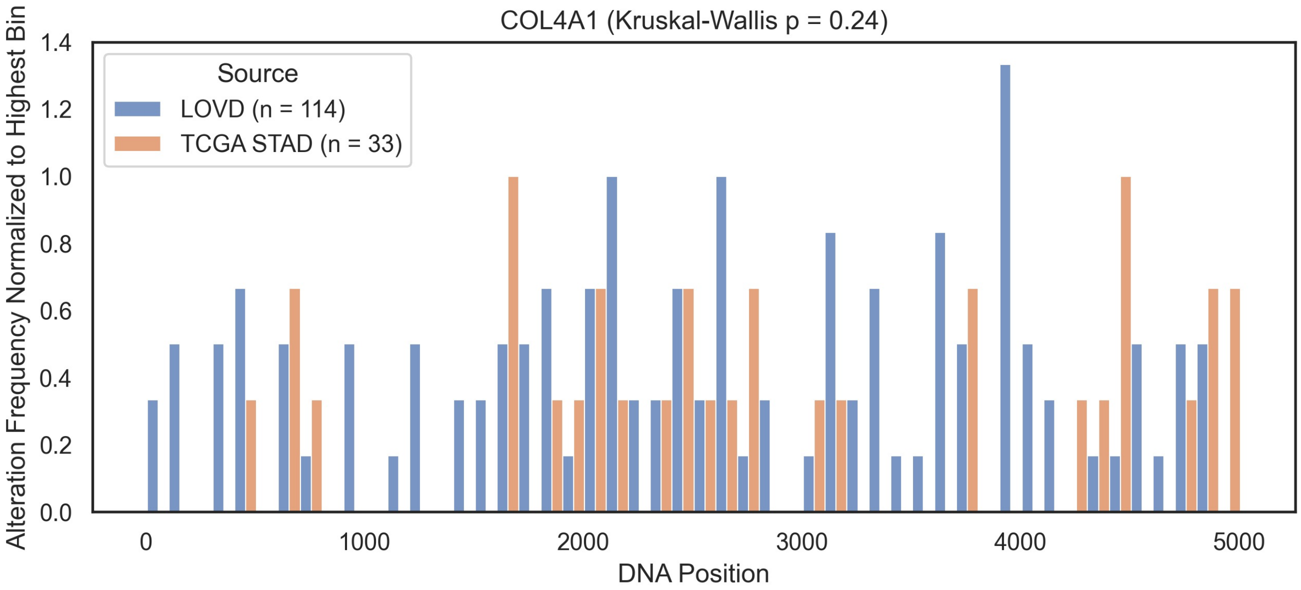

COL4A1 (Kruskal-Wallis p = 0.24)
Source
LOVD (n = 114)
TCGA STAD (n = 33)
Alteration Frequency Normalized to Highest Bin
DNA Position


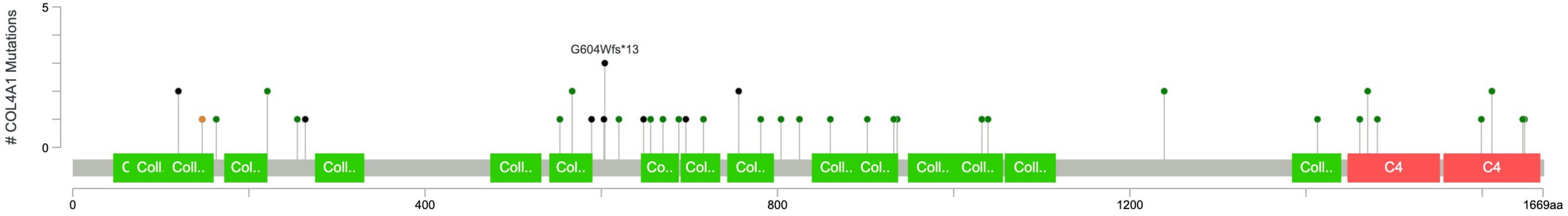

### COL4A1 Mutations
G604Wfs*13
C4
C4
1669aa

Figure S12C

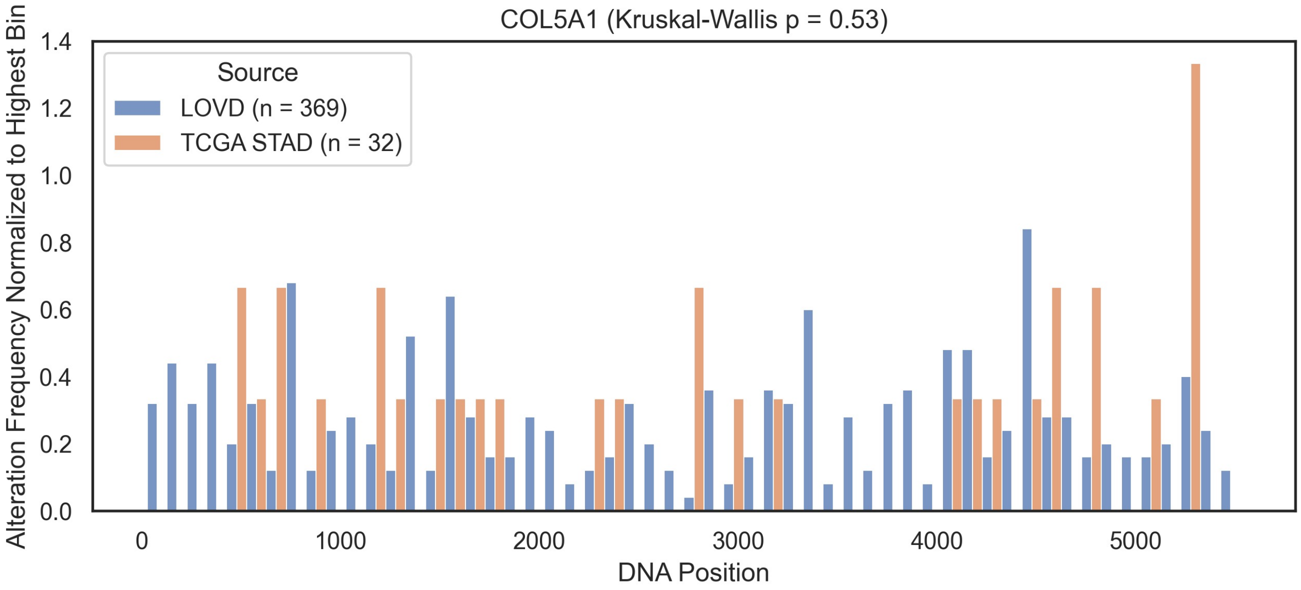

COL5A1 (Kruskal-Wallis p = 0.53)
Source
LOVD (n = 369)
TCGA STAD (n = 32)
Alteration Frequency Normalized to Highest Bin
0.0
0.2
0.4
0.6
0.8
1.0
1.2
1.4
0
1000
2000
3000
4000
5000
DNA Position


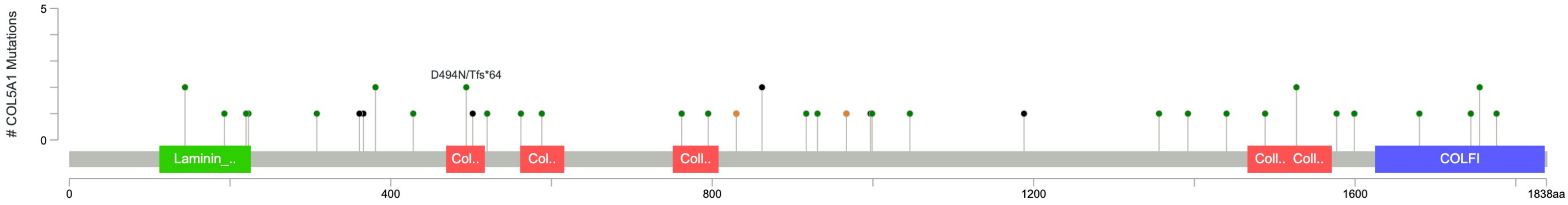

### COL5A1 Mutations
5
0
D494N/Tfs*64
Laminin_..
Col..
Col..
Coll..
Coll..
Coll..
COLFI
0
400
800
1200
1600
1838aa
